# Large-Scale Comparative Analysis of Codon Models Accounting for Protein and Nucleotide Selection

**DOI:** 10.1101/174839

**Authors:** Iakov I. Davydov, Nicolas Salamin, Marc Robinson-Rechavi

## Abstract

There are numerous sources of variation in the rate of synonymous substitutions inside genes, such as direct selection on the nucleotide sequence, or mutation rate variation. Yet scans for positive selection rely on codon models which incorporate an assumption of effectively neutral synonymous substitution rate, constant between sites of each gene. Here we perform a large-scale comparison of approaches which incorporate codon substitution rate variation and propose our own simple yet effective modification of existing models. We find strong effects of substitution rate variation on positive selection inference. More than 70% of the genes detected by the classical branch-site model are presumably false positives caused by the incorrect assumption of uniform synonymous substitution rate. We propose a new model which is strongly favored by the data while remaining computationally tractable. With the new model we can capture signatures of nucleotide level selection acting on translation initiation and on splicing sites within the coding region. Finally, we show that rate variation is highest in the highly recombining regions, and we propose that recombination and mutation rate variation, such as high CpG mutation rate, are the two main sources of nucleotide rate variation. While we detect fewer genes under positive selection in Drosophila than without rate variation, the genes which we detect contain a stronger signal of adaptation of dynein, which could be associated with *Wolbachia* infection. We provide software to perform positive selection analysis using the new model.

## I. Introduction

Detecting the selective pressure affecting protein coding genes is an important component of molecular evolution and evolutionary genomics. Codon models are one of the main tools used to infer selection on protein coding genes (Koonin and Wolf, 2010). This is done by comparing the rate of nonsynonymous substitutions (*d*_*N*_) that are changing the amino acid sequence with the rate of synonymous substitutions (*d*_*S*_) that do not affect this amino acid sequence.

While there is overwhelming evidence of negative and positive selection acting on the amino acid sequence of the proteins (Boyko et al., 2008), synonymous substitutions affecting the protein coding genes are assumed to be effectively neutral in most current models. This is a reasonable first approximation, especially for species with low effective population size, such as many mammals (Keightley et al., 2005; Romiguier et al., 2014). Therefore the synonymous substitution rate can be used as a proxy for the neutral substitution rate, and comparison between *d*_*N*_ and *d*_*S*_ can be used to identify selection acting on the level of amino acids (Yang and Bielawski, 2000).

A corollary of this assumption has been that most codon models assume that synonymous substitution rates are uniform across each gene. Yet there is no biological reason to assume this uniformity, and actually some evidence against it (for a review see Rubinstein and Pupko 2012). This is expected to particularly affect more sophisticated models, where *ω* varies between branches and sites. Indeed, violation of the assumption of uniformity of the synonymous rate can affect the performances of the site-model (Rubinstein et al., 2011).

There are numerous sources of variation in the rate of synonymous substitutions inside genes. First, the raw mutation rate across each genome varies significantly. One of the strongest effects on the mutation rate in mammals is CpG sites. Transitions at CpG sites are more that 10-fold more likely than transversions at non-CpG sites (Leffler et al., 2013) due to spontaneous deamination, which causes a mutation from C to T, or from G to A. Both mutation frequencies and repair efficiency are highly dependent on the context. For example, the mammalian CpG mutation rate is lower in high GC regions (Fryxell and Zuckerkandl, 2000). This is probably related to strand separation and hydrogen bonding in the neighboring region (Segurel et al., 2014). High GC regions themselves are characterized by a higher mutation rate, which is probably caused by less efficient repair by the exonuclease domain. There are other context-dependent effects which are known, many of which lack a mechanistic explanation, such as a higher mutation rate away from T with an increasing number of flanking purines (Hwang and Green 2004, for reviews see Hodgkinson and Eyre-Walker 2011 and Segurel et al. 2014).

Mutation rate is also affected by replication time and it has been shown to be higher in late-replicating regions (Stamatoyannopoulos et al., 2009). This effect has been attributed both to the interference between RNA and DNA polymerases (Jørgensen and Schierup, 2009) and to variation in the efficiency of mismatch repair (Supek and Lehner, 2015). It is not clear that this affects variation within genes, as opposed to between genes, but it could do so in very long genes.

Mutation rates are correlated with recombination rates. Some suggest (Lercher and Hurst, 2002; Hellmann et al., 2003, 2008) that recombination itself can have a mutagenic effect, possibly through an interaction with indels. Evidence from Drosophila suggests that recombination is not mutagenic, but does influence local *N*_*e*_ and thus the efficiency of selection (Castellano et al., 2018). Alternatively, this correlation can be a result of GC-biased gene conversion (GC-BGC), whereby mutations increasing GC content have a higher chance of fixation in the population (Duret and Galtier, 2009). While GC-BGC is a fixation bias, in some cases it can create a pattern which is hard to distinguish from positive selection (Ratnakumar et al., 2010).

Finally, the synonymous substitution rate can be affected by selection at the nucleotide level. First, while synonymous substitutions do not affect the protein sequence, they might affect translation efficiency. This effect is not limited to species with large effective population size, such as Drosophila (Carlini and Stephan, 2003), since selection for codon usage was identified even in *Homo sapiens* (Comeron, 2004) and other mammals, especially for highly expressed genes. It has been suggested that bias in codon usage reflects the abundance of tRNAs, and thereby provides a fitness advantage through increased translation efficiency or accuracy of protein synthesis (Bulmer, 1991), although in many cases there is no dependency between tRNA abundance and codon frequency, and the source of the bias remains unknown (Plotkin and Kudla, 2011). Selection at the nucleotide sequence can be also caused by secondary structure avoidance, as secondary structure can reduce translation efficiency (Kudla et al., 2009; Kertesz et al., 2010). Other potentially important sources of selection on the nucleotide sequence, independent of the coding frame, include splicing motifs located within exons, exon-splicing enhancers (Majewski and Ott, 2002), or genes for functional non-coding RNAs, such as miRNAs or siRNAs, which often reside within coding sequences (Mattick and Makunin, 2006).

Because of all these mutational and selective effects, it is important to model rate variation not only at the level of protein selection, but also at the nucleotide level. There are in principle two different approaches to incorporate rate variation into codon models. We can extend either the Muse and Gaut (1994) model, where both *d*_*N*_ and *d*_*S*_ are estimated as two independent parameters, or the Goldman and Yang (1994) model, where selection pressure on the protein sequence is represented by a single parameter (*ω*) that defines the ratio of nonsynonymous to synonymous substitutions (*d*_*N*_ /*d*_*S*_). First, it is possible to model synonymous (*d*_*S*_) and non-synonymous (*d*_*n*_) substitution rates separately by extending a two-rate model, as in Pond and Muse (2005) and Mayrose et al. (2007). Second, it is possible to incorporate site-specific rates as an independent parameter into single parameter models (Scheffler et al., 2006; Rubinstein et al., 2011). In the second case, the substitution rate parameter captures biological factors acting on all substitutions, both synonymous and nonsynonymous. These factors can include mutation rates, fixation rates, or nucleotide selection.

Here we focus on the second approach, which is traditionally used for large-scale positive selection analyses in eukaryotes (Clark et al., 2007; Markova-Raina and Petrov, 2011; Zhang et al., 2014; Moretti et al., 2014). Spielman et al. (2016) reports superior performance of a compound parameter for the estimation of selection strength on the protein. While we use a single parameter *ω* to model the selection strength at the protein level, *ω* can vary both between alignment sites and between tree branches.

While codon models accounting for nucleotide rate variation have been available for more than a decade, they are still rarely used for large-scale selection analyses, such as Kosiol et al. (2008); Moretti et al. (2014); Zhang et al. (2014). This is probably because these models have even higher computational demands, and the statistical performance of different approaches to nucleotide rate variation was never compared.

Here we extend the Scheffler et al. (2006) model, which captures variation between codons, i.e., uses a single rate per codon, and perform a direct comparison with the model of Rubinstein et al. (2011), which captures variation between nucleotides, i.e., with three rate parameters per codon. Thanks to the computational efficiency of our method, we can show that synonymous rate variation is pervasive, and impacts strongly the detection of branch-site positive selection.

We also assess the impact of nucleotide rate variation on the BS-REL-family model (Murrell et al., 2015). Models based on Goldman and Yang (1994) typically use the maximum likelihood approach on the two nested models in order to detect positive selection. This way the method can identify genes, while individual sites can be detected only using additional posterior analysis. This approach seems to be suitable for large-scale genome analyses, where one is interested in identifying biological functions undergoing positive selection (Clark et al., 2007; Kosiol et al., 2008; Zhang et al., 2014; Daub et al., 2017). We chose Murrell et al. (2015) as a comparison, since it is the only BS-REL model for gene-wide identification of positive selection, while other positive selection models in that family are intended for inference of selection at individual sites, and, thus, cannot be compared directly.

We first use simulations to compare different approaches of modeling synonymous rate variation. Then, we use our model to detect positive selection in twelve Drosophila species and in a vertebrate dataset.

We detect positive selection on genes from those two datasets under our new model, and we demonstrate that it is important to take rate variation into account for such inference of positive selection. We investigate factors affecting the nucleotide substitution rate, and we show that the new model successfully detects synonymous selection acting on regulatory sequences within the coding sequence. We also identify what are the gene features that affect rate heterogeneity the most.

## II. New approaches

We model the process of codon substitution as a Markov process defined by the instantaneous rate matrix *Q*. In a general case, *Q* can be written as (Rubinstein et al., 2011):

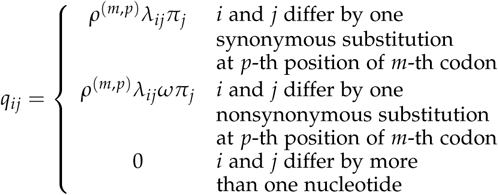

Here *ρ*^(*m,p*)^ is the substitution rate for *ρ*-th position Here r(m,p) is the substitution rate for p-th position (*p* ∈ {1, 2, 3}) of codon site *m* of an alignment (*m* ∈ 1 … *M*, where *M* is the alignment length in codons). The variable *λ_ij_* is the substitution factor to change from codon *i* to codon *j*. It is typically used to account for the difference between transition and transversion rates (Hasegawa et al., 1985). In this case, *λ_ij_* depends on the substitution type but does not depend on the codon position in which substitution occurs. The rate *ρ*^(*m,p*)^ is used to account for various effects that are not captured by the variation in *ω*. In particular, it accounts for variation in mutation rate and selection acting on the nucleotide sequence.

In Rubinstein et al. (2011), *ρ*^(*m,p*)^ is modeled using a one parameter gamma distribution across sites of the alignment, such that the mean relative substitution rate is equal to 1, i.e., *ρ*^(*m,p*)^ ∼ *Gamma*(*α*, 1/*α*). Keeping a mean rate of 1 is important to avoid biases in the estimation of branch lengths. There is no implicit assignment of rates to sites, as in the CAT model (Lartillot and Philippe, 2004). Instead, a random-effect model is used: the gamma distribution is split into *K* equally probable discrete categories *ρ*_*k*_ using quantiles, and the site likelihood is computed as the average of the likelihoods for each possible rate assignment. This approach adds only one extra parameter to the model, but it is computationally intensive. Indeed, in order to compute a likelihood for the *m*-th codon in the alignment, it is necessary to compute likelihoods for this codon given all possible rate assignments, i.e.,

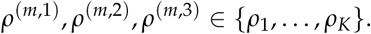

For *K* discrete categories, *K*^3^ likelihoods have thus to be computed per codon site.

In Scheffler et al. (2006), unlike Rubinstein et al. (2011), the three positions of each codon have the same rates, i.e., *ρ*^(*m*,1)^ = *ρ*^(*m*,2)^ = *ρ*^(*m*,3)^; we denote the rate of a codon *m* as *ρ*^(*m*,*)^. Here a codon belongs to one of three categories, each one represented by a single rate value *ρk*_*k*_. The rates and their respective proportions are estimated from the data, which leads to the estimation of four different parameters: two rate parameters *R*_1_ < 1 and *R*_2_ > 1 and two proportion parameters 0 < *p*_1_ < 1 and 0 < *p*_2_ < 1. Effective proportions are computed as follows: 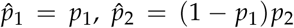, and 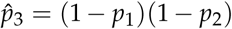. Rates are computed as: *ρ*_1_ = *sR*_1_, *ρ*_2_ = *s*, and *ρ*_3_ = *sR*_2_, where *s* is a scale factor chosen such that the mean rate is equal to one: 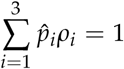. This approach is virtually equivalent to adding a branch length multiplier for certain site classes, and therefore likelihood can be computed efficiently.

Here we propose having one rate per codon *ρ*^(*m*,1)^ = *ρ*^(*m*,2)^ = *ρ*^(*m*,3)^, while allowing this rate to vary following a gamma distribution, *ρ*_*k*_ ∼ *Gamma*(*α*, 1/*α*). The same approach to model rate variation has already been used (Gil et al., 2013; Baele and Lemey, 2013), but it was restricted to the assumption of constant selective pressure (*ω*) across all sites and phylogenetic branches.

Our approach is closely related to Scheffler et al. (2006), as we are modeling a single rate per codon. The distinction is that we are using unit gamma-distribution to model rates because of its flexibility while being controlled by a single parameter, as opposed to four parameters required for the 3-rate model of Scheffler et al. (2006). The approach described in Rubinstein et al. (2011) also uses gamma distribution to model rates, but those rates are associated with a single nucleotide site, not a single codon site, which substantially increases the computational complexity. In the approaches proposed by Pond and Muse (2005) and Mayrose et al. (2007), gamma distributed rates are also assigned to individual codons. The important distinction is that in those cases synonymous and non-synonymous rates are modeled separately, which makes estimation of selection as a ratio between the two rates more challenging (see also Spielman et al., 2016).

Using our approach, we extended two widely used codon models: the site model M8 (Yang et al., 2000) and the branch-site model (Zhang et al., 2005). In principle our approach could be applied to any GY94-based model. In M8, selection pressure represented by the *ω* parameter varies between the sites of an alignment following a beta distribution, while staying constant over the branches of the phylogenetic tree. In this model, a subset of sites can evolve under positive selection. In the branch-site model, *ω* varies both between the sites of the alignment and the branches of the phylogenetic tree. In this model, a subset of sites can thus evolve under positive selection on a predefined subset of branches. These two models were implemented in Godon, a codon model optimizer in Go, in four variants: no rate variation, site rate variation (Rubinstein et al., 2011), codon 3-rate variation (Scheffler et al., 2006), and codon gamma rate variation as described above.

Four of the eight models were implemented and used for the first time to our knowledge: branch-site models with site rate variation similar to Rubinstein et al. (2011), codon 3-rate variation similar to Scheffler et al. (2006), gamma distributed codon rate variation as proposed above, and M8-based model with gamma distributed codon rate variation. All models were implemented within a common framework, ensuring fair comparisons.

## III. Results

### I. Simulations

#### Site models

We have simulated four datasets using various flavors of the M8 model: a dataset without rate variation, a dataset with site rate variation, a dataset with gamma distributed codon rate variation, and a dataset with codon 3-rate variation (Table 1). We then used the four corresponding models to infer positive selection in those datasets. In all four cases, as expected, the model corresponding to the simulations shows the best result in terms of receiver operating characteristic (ROC, Fig. 1, Table 2) as well as accuracy (Supplementary Table S1), and precision versus recall (Supplementary Fig. S1).

**Table 1:**
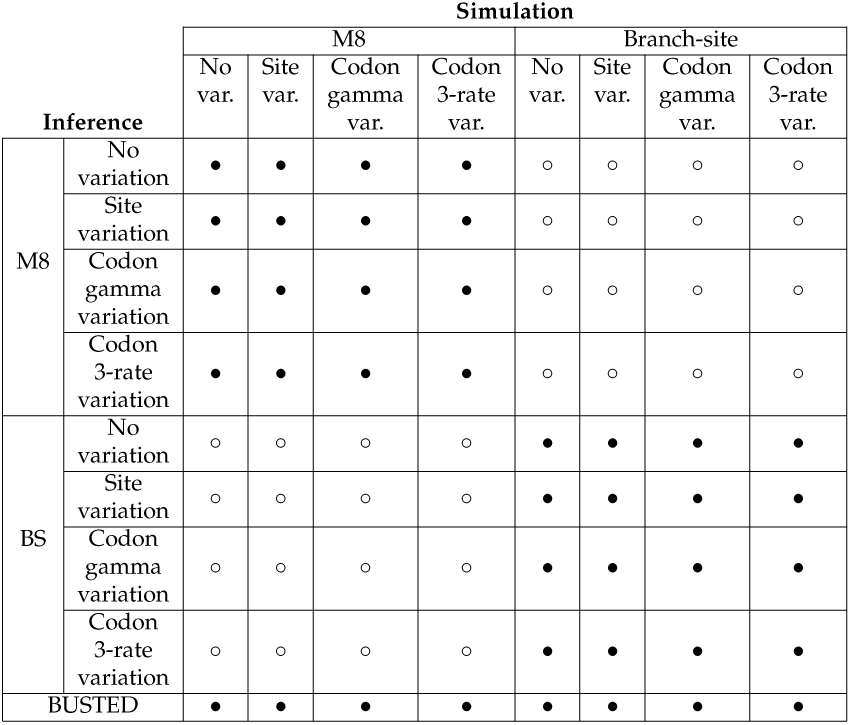
Summary of estimations performed on the simulated datasets; bullets and circles indicate which models were used to simulate datasets and which models were used for inference on these datasets. Combinations indicated with bullets are discussed in the main text, while combinations indicated with circles are discussed in the supplementary. M8: M8 model of Yang et al. (2000); BS: branch-site model of Zhang et al. (2005); BUSTED: BUSTED model from the BS-REL-family (Murrell et al., 2015). Codon variation gamma refers to the proposed parametrization, while codon variation 3-rate refers to the parametrization introduced in Scheffler et al. (2006)

**Table 2:**
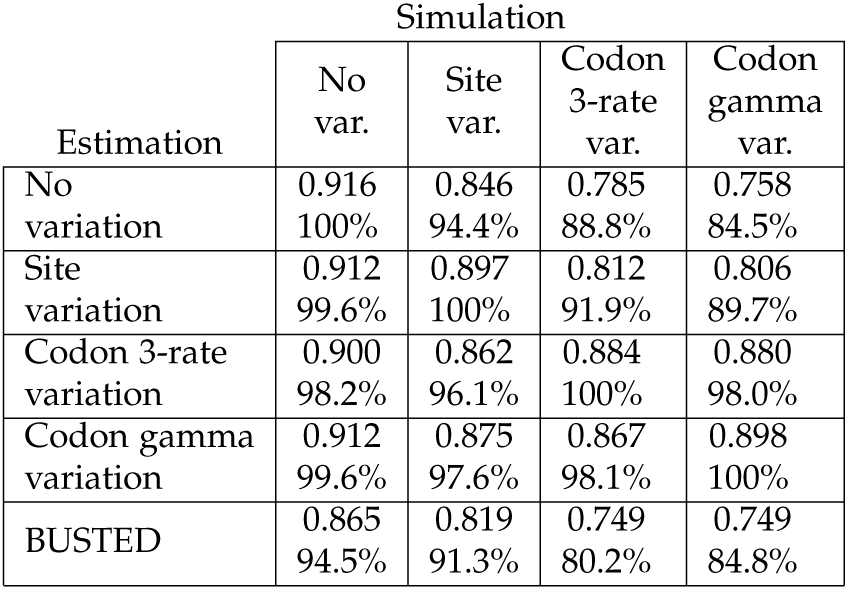
Area under curve (AUC) for all M8-based simulations (see Fig. 1) and for BUSTED. Second number computed as proportion of maximum AUC for a particular simulation.

**Figure 1:**
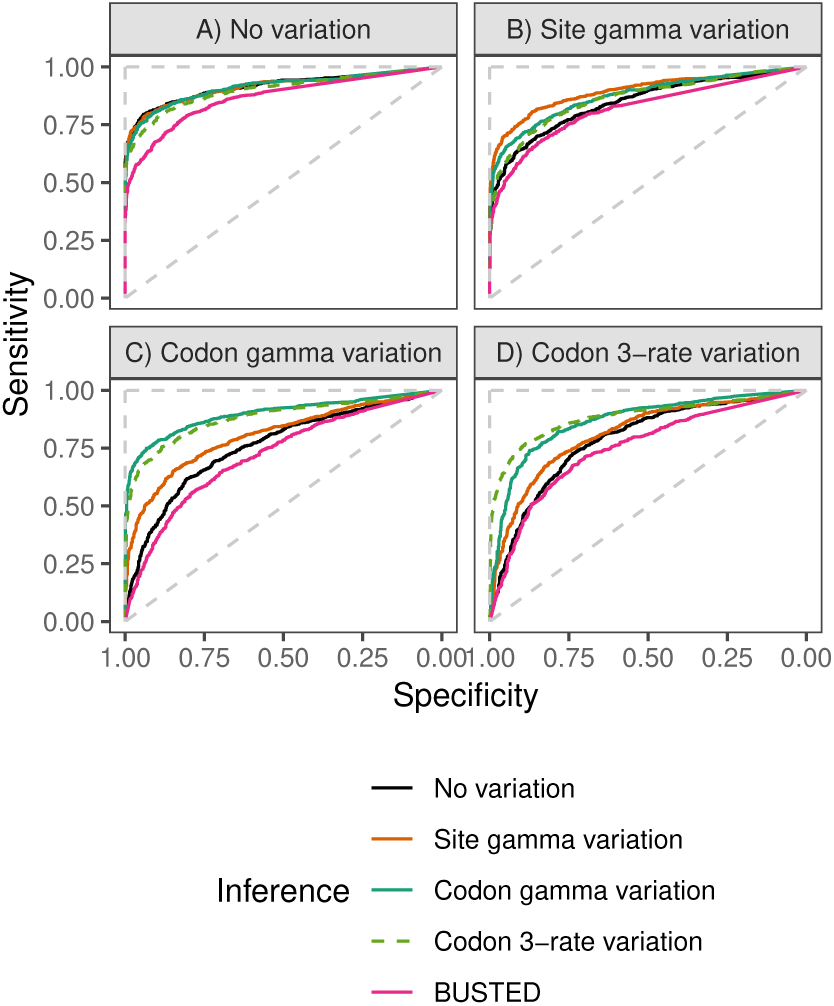
ROC of four M8-based models (M8 with no rate variation, M8 with site rate variation, M8 with codon gamma rate variation, and M8 with codon 3-rate variation) and of BUSTED on datasets A) without rate variation, B) with site rate variation, C) with codon gamma rate variation, and D) with codon 3-rate variation. Specificity is defined as the proportion of correctly identified alignments simulated under a model with positive selection, sensitivity is defined as the proportion of correctly identified alignments simulated without positive selection. The dashed diagonal line shows theoretical performance of the random predictor, the dashed vertical and horizontal lines indicate theoretical performance of the perfect predictor.

In the absence of rate variation, the statistical performance of the four methods is very similar, even though the M8 model without rate variation has a slightly better ROC (Fig. 1A), a false positive rate (FPR) which is closer to the theoretical expectation (Supplementary Fig. S2A), and a slightly higher sensitivity (Supplementary Fig. S3A). Despite increased complexity, the sensitivity of the codon gamma variation model is only marginally reduced relative to the model without rate variation (Supplementary Fig. S3A).

The M8 model without rate variation largely underperforms on the dataset with site rate variation: it has a worse ROC (Fig. 1B) and a higher FPR (Supplementary Fig. S2B). On the other hand, codon gamma variation performs almost as well as site variation, and clearly better than the model with no variation.

For both datasets with codon rate variation, there is a relatively large decrease in the performance of models both without rate variation and with site rate variation. ROC is decreased (Fig. 1C,D) and FPR is inflated (Supplementary Fig. S2C,D). Performance of the two variants of codon rate variation is similar. Codon gamma rate variation even increases the false positive rate above 50% for the model without rate variation at the significance level of 0.05. Stronger rate variation (i.e., smaller *α* value) causes a higher false positive rate (Supplementary Fig. S4).

From this we can conclude that a) the performance of models accounting for codon variation is acceptable in all three scenarios, i.e., no rate variation, site rate variation, and codon rate variation; b) in the presence of codon rate variation in the data, models not accounting for this kind of variation suffer from a notable loss of statistical performance.

We have mainly focused on a realistic scenario in which true branch lengths are unknown. However, to confirm that our results are not biased by the differences in branch lengths, we also fit the models using the true branch lengths. In this case, performance is very similar, both in terms of ROC (Supplementary Fig. S5) and of false positive rates (Supplementary Fig. S6).

#### Branch-site models

The simulations based on the branch-site model show a qualitatively similar behavior to the simulations based on the M8-type models regarding ROC (Fig. 2), FPR (Supplementary Fig. S7), AUC, and precision (Supplementary Tables S2, S3), and precision versus recall (Supplementary Fig. S8), although the performances are more similar between models. As with M8-type models, codon rate variation models perform well in all four cases, while simulating with codon rate variation causes a clear underperformance in both other models. Unlike in the case of M8-type models, false positive rates are only marginally inflated compared to theoretical expectations with smaller values of *α* (Supplementary Fig. S9). Nevertheless, the models with codon rate variation show the best performance.

**Figure 2:**
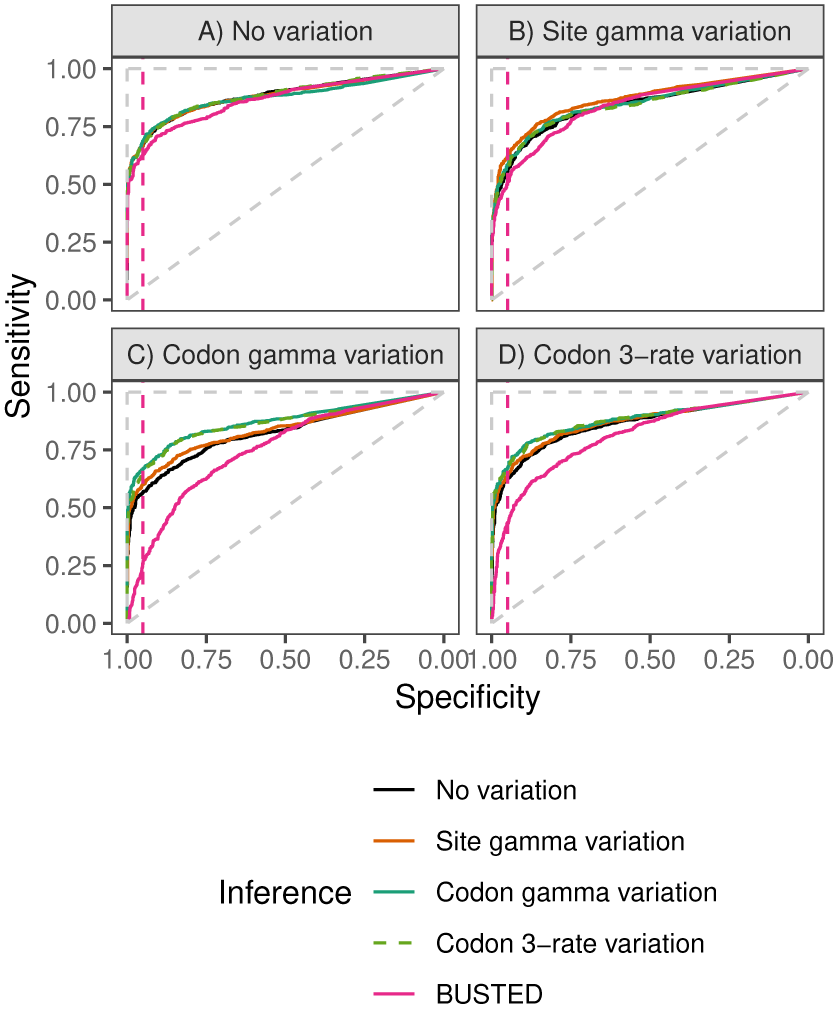
Performance (ROC) of four branch-site-based models (branch-site with no rate variation, branch-site with site rate variation, branch-site with codon gamma rate variation, and branch-site with codon 3-rate variation) and of BUSTED on datasets A) without rate variation, B) with site rate variation, C) with codon gamma rate variation, and D) with codon 3-rate variation. The pink dashed line indicates the 0.95 specificity threshold (i.e., false positive rate of 0.05). The dashed diagonal line shows theoretical performance of the random predictor, the dashed vertical and horizontal lines indicate theoretical performance of the perfect predictor.

When using true or estimated branch lengths performance is very similar (ROC: Supplementary Fig. S10). The main difference is an increase in the false positive rate of the model without rate variation (Supplementary Fig. S11).

More complex models have a computational cost. Analyses with codon gamma rate variation were 3.3 and 2.8 times slower compared to no rate variation for M8 and branch-site models respectively. The codon 3-rate variation model by Scheffler et al. (2006) provides a similar statistical performance while having a higher computational cost: 3.7 and 6.7 times slower compared to no rate variation, respectively. This increased computational load might be explained by a larger dimensionality of parameter space. Site rate variation models were 62.6 and 44.7 times slower, respectively (Table 3). Thus codon rate variation captures biological signal at a much lower computational cost than site rate variation, especially when coupled with a gamma model.

**Table 3:** Inference time of different models. Top lines show total CPU-hours used, bottom lines indicate relative slowdown compared to the fastest model. For the simulated datasets only time used by models of the same family is shown, i.e., comparison between M8 and branch-site models is not included.

| Dataset | Inference model |  |  |  |
| --- | --- | --- | --- | --- |
|  | No var. | Site var. | Codon var. gamma | Codon var. 3-rate |
| M8 simulations | 91h<br>1x | 5,723h<br>63x | 300h<br>3.3x | 339h<br>3.7x |
| Branch-site simulations | 47h<br>1x | 2,126h<br>46x | 132h<br>2.8x | 318h<br>6.7x |
| Vertebrate dataset | 55h<br>1x | 3,837h<br>69x | 258h<br>4.7x | 481h<br>8.7x |
| Drosophila dataset | 966h<br>1x | – | 5,746h<br>5.9x | – |
| Drosophila dataset<br>1,000 alignments | 115h<br>1x | 10,079h<br>88x | 677h<br>5.9x | 1,278h<br>11x |

#### Comparisons with BS-REL

It has been demonstrated (Murrell et al., 2015) that in certain cases the statistical power of BS-REL is superior to other methods, including the branch-site model. Therefore it is important to study how rate variation affects the performance of those models. The only BS-REL model suitable for the gene-wide identification of positive selection is BUSTED (Murrell et al., 2015), and the current implementation supports neither rate variation nor *d*_*S*_ variation (implemented in Pond and Muse 2005).

The BS-REL framework differs from the branch-site and M8 models by the frequency parametrization. BS-REL is based on the approach of Muse and Gaut (1994), while the branch-site and M8 models are based on the approach of Goldman and Yang (1994). During simulations, we used F0 frequencies, i.e., identical frequencies of all the codons (*π*_*i*_ = 1/61). As F0 can be considered a special case of both Muse and Gaut (1994) and Goldman and Yang (1994), we do not expect any bias caused by this difference between the two approaches.

BUSTED shows significantly inflated rates of false positives in the presence of codon gamma rate variation (Supplementary Fig. S2C, S7C, simulations under M8 and branch-site, respectively). At a typical significance level of 0.05, the false positive rate of BUSTED is close to 0.3 and 0.2 for the M8 and branch-site simulations, respectively (i.e. the proportion of false positives can be 4-6 times higher than we expect by chance). This shows that the statistical performance of the BS-REL family models is also affected by not taking into account rate variation.

### II. Vertebrate dataset

Given the good performance of the codon gamma rate model in the simulations, we applied it to real data. First we used 767 one-to-one orthologs from vertebrate species. This represents a set of genes with high divergence (more than 450 My), conservative evolution (Studer et al., 2008), and relatively low effective population sizes (although some vertebrates have high *N*_*e*_, see Gossmann et al. 2012), thus relatively weak impact of natural selection. We analyzed them with four variants of the branch-site model: no rate variation, site rate variation, codon gamma rate variation, and codon 3-rate variation. We used the branch-site model to search for positive selection, as it is more sensitive to episodic positive selection (see Supplementary Materials, page 3).

In most cases (each case corresponding to a single branch of a gene tree), the data supports the codon rate variation models based on Akaike information criterion (AIC): out of 8,907 individual branches tested, data supports codon gamma rate variation in 43% of the tests, codon 3-rate variation in 47%, site rate variation in 9.4%, and no rate variation model was favored only in 12 tests (0.1%).

A large proportion (43%) of branches detected to be under positive selection with the no rate variation model are not detected to be under positive selection with the codon rate variation model (Table 4). This effect is even stronger (72%) when multiple testing correcion is used (Table 4B). The majority of the positive predictions from the standard branch-site model are not supported when both multiple testing and codon rate variation are accounted for. This suggests that evolution on these branches can be explained by nucleotide substitution rate variation without positive selection (see Discussion). We observe relatively high agreement in predictions between the site rate variation and codon rate variation models (after multiple testing correction), as those are two different approaches to model the same evolutionary process. Supplementary Fig. S12 shows the number of genes detected by the branch-site models with codon gamma rate variation and without rate variation, mapped to the vertebrate species phylogeny. There was no positive selection identified using M8 models, with or without codon gamma rate variation.

**Table 4:** Positive selection predictions for the vertebrate dataset. Numbers in each cell indicate how many branches were detected (+) or not detected (-) to be evolving under positive selection by different variants of the branch-site model. A,B) no rate variation versus codon gamma rate variation; C,D) no rate variation versus codon 3-rate variation; E,F) no rate variation versus site rate variation; G,H) site rate variation versus codon rate variation; A,C,E,G) significance threshold of 0.05; B,D,F,H) false discovery rate threshold of 0.1.

|  |  |  |
| --- | --- | --- |
| **A** Codon gamma variation |  |  |
| No variation | – | + |
| – | 7,063 | 58 |
| + | 823 | 963 |
| **B** Codon gamma variation |  |  |
| No variation | – | + |
| – | 7,660 | 6 |
| + | 966 | 275 |
| **C** Codon 3-rate variation |  |  |
| No variation | – | + |
| – | 7,059 | 62 |
| + | 777 | 1,009 |
| **D** Codon 3-rate variation |  |  |
| No variation | – | + |
| – | 7,660 | 6 |
| + | 951 | 290 |
| **E** Site variation |  |  |
| No variation | – | + |
| – | 7,069 | 52 |
| + | 516 | 1,270 |
| **F** Site variation |  |  |
| No variation | – | + |
| – | 7,657 | 9 |
| + | 694 | 547 |
| **G** Codon gamma variation |  |  |
| Site variation | – | + |
| – | 7,513 | 72 |
| + | 373 | 949 |
| **H** Codon gamma variation |  |  |
| Site variation | – | + |
| – | 8,347 | 4 |
| + | 279 | 277 |

Supplementary Table S4 shows prediction agreement between each model and the best supported model out of four, confirming the good performance of the codon variation model. While the codon 3-rate model has a slightly higher proportion of tests which support it, the codon gamma rate variation model has the highest prediction agreement with the best model (Supplementary Table S4B, C).

Codon gamma rate variation model was 4.7 times slower, while codon 3-rate variation model was 8.7 times slower, and site rate variation model was 69 times slower (Table 3) than no rate variation.

With real data, differences between genes are not only stochastic, but more interestingly are expected to be driven by underlying biological differences. It is thus interesting to find which factors affect rate variation as estimated by the model, as well as to know which genes favored the model with codon rate variation the most. We focused on gene features associated with the underlying evolutionary process, such as recombination rate, GC content standard deviation (indicative of shifts in recombination hotspots, Glemin et al. 2015) and expression level (associated with stronger purifying selection, Drummond et al. 2005; Pal et al. 2006; Kryuchkova-Mostacci and Robinson-Rechavi 2015). Parameters which can directly affect the performance of the method were also included in the linear model to avoid potential biases, e.g., number of sequences and alignment length. We also included total intron length and number of exons, since they can affect synonymous selection associated with splicing, or disparity between mutation rates associated with chromosomal localization of exons.

Here and below we used three response variables for our analyses. First, we create linear models using the relative support of the model (based on Akaike weights, see Methods) as a response variable. These models allow us to understand for which categories of genes the effect of rate variation is the strongest. Second, models using the *α* parameter of the gamma distribution (codon rate variation) as a response variable allow us to identify gene properties associated with high substitution rate variance. Finally, a model for the proportion of branches which are inferred to have evolved under positive selection when rate variation is not taken into account, but not when it is, allows us to identify the main causes of discrepancy between the results of the two models.

In this analysis, each group of orthologous genes was treated as a single observation. Estimated parameter values obtained by testing different branches of the tree were averaged.

The relative support of the model with codon gamma rate variation is mostly affected by total branch length, alignment length, and mean GC content of the gene (Supplementary Table S5). The positive correlation with tree length and alignment lengths is probably related to the increase in total amount of information available for the model. The relation to GC content might be due to the relationships between recombination rates, substitution rates, and GC content (Duret and Galtier 2009; Rudolph et al. 2016, see Discussion).

For the shape parameter of the gamma distribution *α*, the strongest explanatory variable is GC content (Supplementary Table S6). As with relative support of the model, this could be related to recombination. We also observe a weak relation with maximal expression level. Highly expressed genes tend to have a higher rate variation, which could be explained by higher nucleotide level selection on certain parts of the gene.

Enrichment analysis did not identify any categories overrepresented among genes detected to be evolving under positive selection with rate variation. This might be due to the small size of the dataset.

## III. Drosophila dataset

The second real dataset we used contains 8,606 one-to-one orthologs from Drosophila genomes. The Drosophila data set is ten-fold larger than the vertebrate dataset. Since analyses on the simulated and vertebrate datasets show a consistent superiority of codon gamma rate over site variation and codon 3-rate variation, with a much lower computational cost, we ran only no variation and codon gamma variation on the full dataset. Therefore for the Drosophila data, we are mainly focusing on comparing models with and without codon rate variation. However, we did run site variation and codon 3-rate variation on a subset of 1,000 genes selected randomly. Drosophila has large effective population sizes on average (Gossmann et al., 2012), thus stronger impact of natural selection; the genes studied are less biased towards core functions than in the vertebrate dataset, and have lower divergence: about 50 mya for Drosophila (Russo et al., 2013) compared to more than 450 mya for the vertebrate dataset (Betancur-R et al., 2015).

In total 66,656 branches were tested for positive selection. The model with codon gamma rate variation was supported by the data in 97% (resp. 96%) of the tests when using AIC (resp. likelihood ratio test, LRT). On the smaller subset, on which all the four approaches were applied, codon gamma rate variation was supported in 48% of the tests, codon 3-rate variation in 23%, site rate variation in 26%, and no variation in 2%. As with the vertebrate dataset, predictions were not consistent between the models (Table 5, comparison of all models for a subset of 1,000 genes in Supplementary Table S7). The site and the codon gamma rate variation models display a stronger consistency in predictions of positive selection relative to the consistency between the model without rate variation and the model with codon gamma rate variation.

**Table 5:**
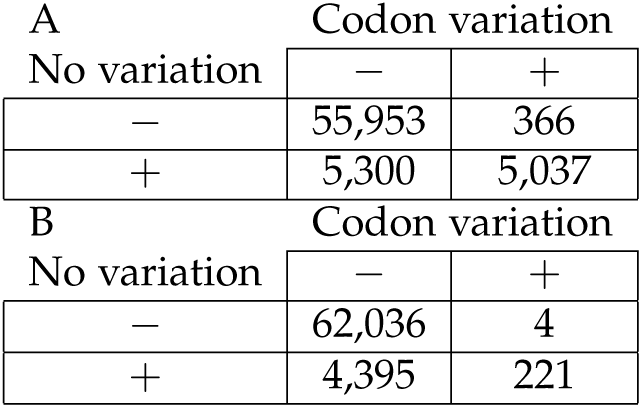
Positive selection predictions for the Drosophila dataset. Numbers in each cell indicate how many branches were detected (+) or not detected (-) to be evolving under positive selection by different variants of the branch-site model. Codon rate variation versus no rate variation. A) statistical significance threshold of 0.05; B) false discovery rate threshold of 0.1.

As in vertebrates, when accounting for multiple testing, the vast majority of predictions of positive selection given by the model without rate variation are not supported by the model accounting for rate variation. Supplementary Fig. S13 shows the number of genes detected by the branch-site models with codon gamma rate variation and without rate variation, mapped to the Drosophila species phylogeny. In addition, there were 4 and 19 genes identified with the M8 models with codon gamma rate variation and without rate variation respectively (for the full lists, see Availability).

Genes identified to be under positive selection with the model accounting for codon gamma rate variation are enriched for molecular function GO categories associated with dynein chain binding (GO:0045503, GO:0045505, for both terms *q*-value= 0.016, Supplementary Table S8). Dynein plays an important role in *Wolbachia* infection (Serbus and Sullivan, 2007), and is thus a likely candidate for strong positive selection (Werren et al., 2008). Surprisingly, there are no significant molecular function GO categories identified using the branch-site model without rate variation (for dynein categories *q*-value=1, Supplementary Table S9). Genes associated with dynein chain binding predicted to have evolved under positive selection, and the relevant amino acid positions, are provided in Supplementary Table S10.

The relative support of the model with codon rate variation is mainly explained by alignment length, number of sequences and coding sequence length (Table 6). Stronger model support associated with increase in the amount of information (increased coding sequence length means less gaps for the same alignment length), expression levels, and mean GC content is consistent with the vertebrate results (Supplementary Table S5).

**Table 6:** Linear model of relative support of model with the codon rate variation; Drosophila dataset. Significant variables (p-value < *0.05) in bold. Model p-value is* < 2.2 · 10-16, adjusted R2 is 0.6133. Model formula: Relative model support ∼ Number of sequences + Total branch length + Alignment length + Length of coding sequence + GC content (mean) + GC content (stdev) + Total intron length + Number of exons + Maximum expression + Mean expression + Recombination rate.

| Variable | Estimate | Std. Error | *t* value | *p*-value |
| --- | --- | --- | --- | --- |
| **Number of sequences** | 0.189 | 0.009 | 22.128 | $< 2 \cdot 10^{-16}$ |
| **Total branch length** | 0.106 | 0.011 | 9.788 | $< 2 \cdot 10^{-16}$ |
| **Alignment length** | 0.574 | 0.030 | 19.138 | $< 2 \cdot 10^{-16}$ |
| **Length of coding sequence** | 0.141 | 0.029 | 4.797 | $1.64 \cdot 10^{-6}$ |
| **GC content (mean)** | 0.084 | 0.009 | 9.789 | $< 2 \cdot 10^{-16}$ |
| GC content (stdev) | 0.011 | 0.007 | 1.445 | 0.1485 |
| **Total intron length** | -0.054 | 0.011 | -4.762 | $1.96 \cdot 10^{-6}$ |
| **Number of exons** | 0.159 | 0.016 | 10.184 | $< 2 \cdot 10^{-16}$ |
| **Maximum expression** | 0.043 | 0.018 | 2.418 | 0.0156 |
| Mean expression | -0.023 | 0.018 | -1.291 | 0.1966 |
| **Recombination rate** | 0.059 | 0.007 | 8.578 | $< 2 \cdot 10^{-16}$ |

We also observe a dependence on the number of exons and on recombination rate. A larger number of exons implies more exon-intron junctions, which might affect variation in levels of nucleotide sequence selection (see below). Recombination might affect GC-BGC, mutation rate, and selection strength acting on synonymous sites (Campos et al., 2014).

The rate variation parameter *α* can be explained by several features of genes (Supplementary Table S11). Most of the effects are not reproduced between the two datasets. While some of them are strongly significant, generally the effect sizes are not very large. The most consistent effect between the two datasets is dependence of the rate variation on GC content.

## IV. Signatures of selection at the nucleotide level

Codon rate variation can be influenced by various factors such as mutation bias, fixation bias (e.g., gene conversion), or selection acting against synonymous substitutions. Notably, it is well known that exon regions adjacent to splicing sites are evolving under purifying selection at the nucleotide level (e.g., see Majewski and Ott, 2002). We determined posterior rates for positions of protein coding gene regions located in the proximity of exon-intron and intron-exon junctions; first exons were excluded from the analysis.

We observe in Drosophila (Fig. 3) that our codon rate variation model captures these selection constraints: the codon substitution rate is lower at the exon-intron junction than at the intron-exon junction, and both have lower rates than the rest of the exon. This is in agreement with splicing motif conservation scores (e.g., see Cartegni et al., 2002), and consistent with negative selection acting on splicing sites.

**Figure 3:**
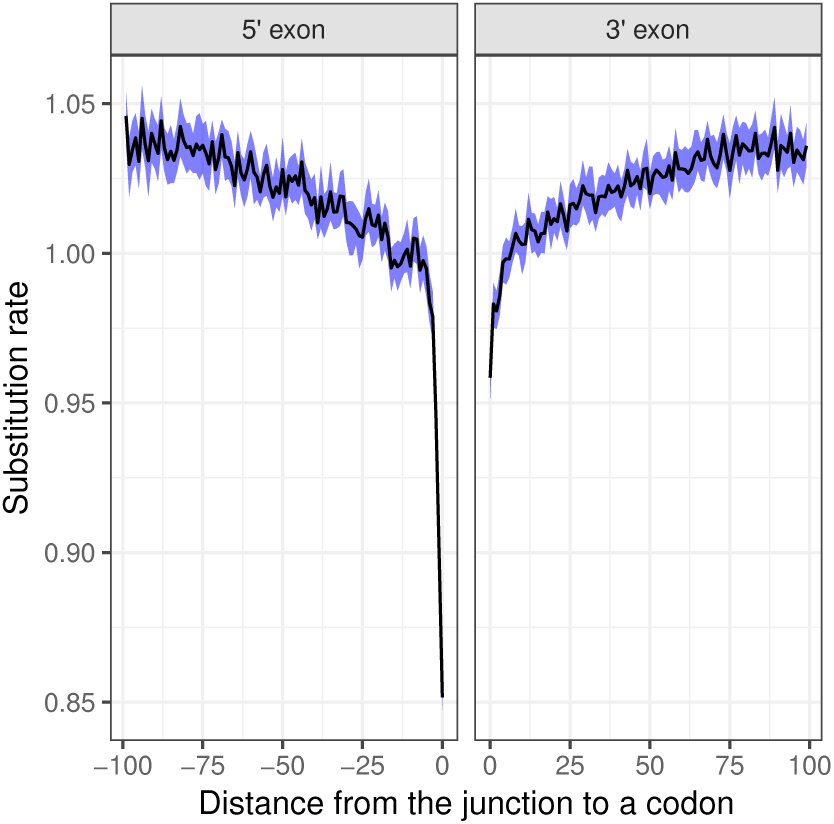
Relative substitution rate as a function of proximity to the exon-intron and intron-exon junctions in the Drosophila dataset. The rates were estimated using the model M8 with codon gamma rate variation. The left panel depicts rates in 5’ exon (prior to the exon-intron junction, negative distances), while the right panel depicts 3’ exon (rates after the intron-exon junction, positive distances). A rate of 1 corresponds to the average rate of substitution over the gene; thus values above 1 do not indicate positive selection, but simply a rate higher than average for this gene. The blue ribbon indicates the 98% confidence interval of mean estimate. Only alignment positions with less than 30% of gaps were used in the plot.

We also used the model M8 with codon gamma rate variation to simultaneously estimate the effect of factors which affect substitution rates of nucleotide and protein sequences, again in Drosophila. We observed that the model is able to recover opposing trends acting on the 5’ region of the protein coding gene (Fig. 4). These trends are probably a result of the high functional importance of the 5’ nucleotide sequence, but low functional importance of the corresponding amino acid sequence (see Discussion). Comparing
nucleotide and inverse protein rates (Supplementary Fig. S14) indicates that the effect is slightly stronger at the protein level; however the difference is only marginal. It is worth noting that this effect cannot be explained by a dependence between the *ρ* and the *ω* estimates, as such a dependence is not observed in other regions (Supplementary Fig. S15).

**Figure 4:**
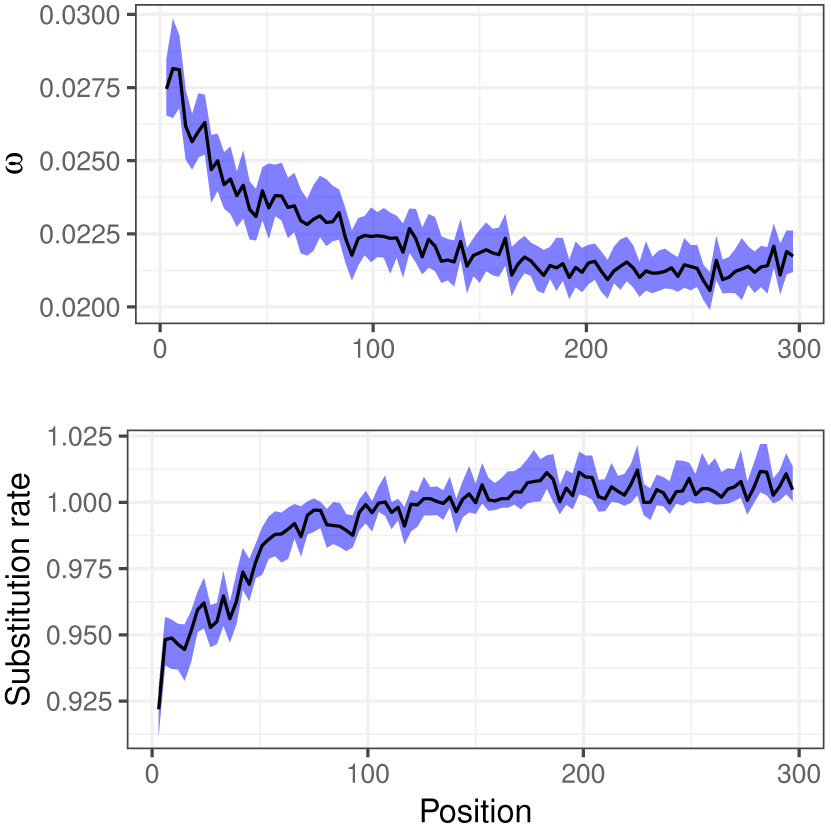
Posterior estimates of median ω (dN/dS, top panel) and codon substitution rate ρ (bottom panel) as a function of distance from the start codon expressed in the number of nucleotides in Drosophila. The model M8 with codon gamma rate variation was used to estimate both parameters simultaneously. Smaller values of ω (top panel) indicate stronger negative selection acting on the protein sequence. A substitution rate of 1 (bottom panel) corresponds to the average rate of substitution over the gene; thus values above 1 do not indicate positive selection, but simply a rate higher than average for this gene. The blue ribbons indicate 98% confidence intervals of median estimates. Start codons and alignment positions with less than three sequences were excluded from the plot.

We observe that the top 25% most highly expressed genes show both stronger conservation of the amino acid sequences (Supplementary Fig. S16) and more pronounced decrease in the substitution rate of the 5’-region (Supplementary Fig. S17). Stronger purifying selection acting on highly expressed proteins was previously observed across domains of life (Pál et al., 2001; Rocha and Danchin, 2004; Drummond et al., 2005; Kryuchkova-Mostacci and Robinson-Rechavi, 2015). As for the decrease in substitution rate, *ρ* is defined relative to the gene-wide average substitution rate. Therefore a stronger relative selection on the 5’-region can be detected above the average increase in purifying selection on highly expressed genes. The relation with expression levels is consistent with the assumption that we are measuring natural selection on gene sequences in this case, rather than mutation rates.

## IV. Discussion

### I. Nucleotide level selection in coding regions

There is strong evidence of selection acting on synonymous substitutions within protein coding sequences, and the strength of this selection is expected to vary across coding regions (Chamary et al., 2006). In particular, negative selection strongly affects regulatory sequences, such as exonic splicing enhancers or exon junction regulatory sequences (Cartegni et al., 2002). Variation in selection over both synonymous and nonsynonymous substitutions can affect the performance of codon models (Rubinstein et al., 2011), and we show that indeed it strongly impacts the results of the popular branch-site model, as well as the simpler site model (M8). While there are multiple ways to account for this variation, for instance by modeling the synonymous and non-synonymous rates separately (Pond and Muse, 2005), here we focused on modeling the ratio of non-synonymous vs synonymous rates as a single parameter (*ω*), while allowing the substitution rate (*ρ*) to vary along the sequence.

Our approach succeeds in recovering a signal of splicing motif conservation jointly with negative and positive selection acting on the protein sequence.

We also demonstrate that our model is able to disentangle opposite trends acting on the same sequence, i.e., stronger negative selection acting on the nucleotide sequence combined with weaker amino-acid selection towards the beginning of the reading frame.

Selection on the 5’ nucleotide sequence is probably due to selection for translation initiation efficiency (Bentele et al., 2013), and is perhaps related to suppression of mRNA structures at the ribosome binding site. At the same time N-terminal amino acids are more likely to be unstructured, and they are relatively less important to protein function and stability compared to the core (Guharoy and Chakrabarti, 2005).

### II. Determinants of rate variation

The majority of gene alignments in the study indicated better support for the model with codon gamma rate variation. Moreover, the relative probability of the models incorporating codon gamma rate variation increases with the amount of information available, be it number of sequences, alignment length, or total number of substitutions. This indicates that these models are better in describing the underlying evolutionary process, and if we have enough data, these models are favored. We detect a strong signal of nucleotide variation in two quite different datasets. Flies have high effective population size, thus natural selection is relatively strong, including on codon usage or splicing. Vertebrates have higher sequence divergence, which does not appear to mask the signal of nucleotide evolution, despite lower effective population sizes in many species (Gossmann et al., 2012; Romiguier et al., 2014). Thus, the effect of nucleotide rate variation appears quite general, and will probably be found in many other species.

The strongest determinant of the relative support of the model with codon rate variation is GC content. A strong effect of GC content on synonymous rate variation was reported by (Dimitrieva and Anisimova, 2014), based on an analysis of protein domain coding sequences with a modified site-model. It is well known (Fullerton et al., 2001; Marais et al., 2003; Chamary et al., 2006) that regions with high recombination rates have higher GC as a result of GC-BGC, notably in the species studied here. It has also been shown that models accounting for rate variation show significantly better performance than simpler models in the presence of recombination (Scheffler et al., 2006), even if the true tree topology is used.

The effect of recombination rate as measured in Comeron et al. (2012) on the relative support of codon rate variation is comparable to the effect of GC content alone. In mammals, the higher CpG dinucleotide mutation rate (Kong et al., 2012) can increase the disparity in substitution rates and therefore contribute to the dependence between the GC content and the relative model support.

Yet, we observe this dependence in Drosophila, where there are no significant neighboring base contextual effects on the mutation rate (Keightley et al., 2009). One explanaion is that the GC content could be acting as a proxy for the average recombination rate over time via GC-BGC. Considering the rapid evolution of recombination hotspots (Ptak et al., 2005), GC content probably captures historical recombination rates, while the direct measurement of recombination rate captures only the current state.

Low and moderate levels of recombination are in general tolerated by codon models, especially models that account for rate variation (Anisimova et al., 2003; Scheffler et al., 2006). However, high levels of recombination could inflate the false positive rate of such models. Therefore, for genes showing the highest rates of recombination, positive selection predictions should be interpreted with caution. Removing the top 30% highest recombination rates has almost no effect on the linear model (Supplementary Tables S12, S13).

In both datasets, we observe a significant positive association of rate variation with the maximal expression level. Pressure for translational robustness increases with expression levels (Drummond et al., 2005), and codon choice affects expression level (Bentele et al., 2013). One of the main causes of selection on the codon sequence of highly expressed genes is protein misfolding avoidance (Yang et al., 2010), but there is also selection for efficient translation initiation (Pop et al., 2014). Consistent with this, Dimitrieva and Anisimova (2014) reported more evidence for synonymous rate variation in genes expressed in the brain. Brain expressed genes are known to be more sensitive to misfolding and under stronger purifying selection (Kryuchkova-Mostacci and Robinson-Rechavi, 2015; Drummond and Wilke, 2008; Roux et al., 2017). It is reasonable to assume that only certain parts of protein coding genes will be affected by strong nucleotide sequence selection, and that this selection will be stronger on more expressed genes. Indeed our results show strong negative selection acting on the coding sequence of translation initiation regions, and the relative selection strength is higher for the 25% highest expressed genes (Supplementary Fig. S17). This can lead to an overall increase in substitution rate variation.

Given that highly expressed genes have both stronger negative selection and stronger variation in substitution rate on the coding sequence, it is especially important to take this variation into account. These genes can easily have a combination of a low average *d*_*S*_, pulled down by strong purifying selection on some regions of the gene, a subset of codons evolving faster than this mean, and a low average *ω*. This can be mistakenly interpreted as positive selection by models without rate variation. On the other hand, with our model we find a positive correlation between support for rate variation and positive selection in Drosophila, even after correcting for confounding variables such as gene length or GC content (linear model of relative model support with proportion of branches under positive selection: Supplementary Table S14). It is possible that these genes are evolving under very strong selection, both positive and negative, or that strong recombination affects the performance of the model. We did not find a significant association in vertebrates (*p*-value=0.24 for the term in a linear model), which could be explained by a smaller size of the dataset. In any case, we cannot confirm a previous report that the genes with the strongest evidence for synonymous rate variation had the less evidence for positive selection (Dimitrieva and Anisimova, 2014).

### III. Codon models and rate variation

Widely used mechanistic codon models rely on the assumption of constant synonymous substitution rates. This assumption is often violated due to factors such as mutation bias or nucleotide selection, which vary across the gene. While substitution rate variation can be caused by multiple factors, we use a single compound rate parameter to model this variation.

We demonstrate that this simple model captures such rate variation, and that it both detects new biological signal, and substantially decreases the false positive rate in positive selection detection. Not only do we observe this effect in simulations (Fig. 1, 2, FPR: Supplementary Fig. S2), but inconsistency between models is even higher when applied to the vertebrate and fly datasets. The vast majority of the predictions of positive selection obtained using models without rate variation were not supported by the model with codon gamma rate variation. The comparable power of the two methods (sensitivity: Supplementary Fig. S3A), and the strong support for rate variation from the data, suggest that most of those positive selection predictions can be explained by nucleotide rate variation (Tables 4, 5). Thus they should be considered as probable false positives. The proportion of those potential false positives per gene alignment is positively associated with the amount of available information, such as alignment length (linear models: Supplementary Tables S15, S16, and S17). This confirms that loss of power is not the main cause for the lack of detection by the codon rate model, but rather that the issue is false positive detection by the classical branch-site.

The effect of rate variation on the model performance is stronger in Drosophila than in vertebrates. On the vertebrate dataset, after multiple testing correction, about 80% of the positive predictions might be false positives (Table 4B), while in Drosophila it is 95% (Table 5B). This might be a consequence of the higher effective population size (Gossmann et al., 2012) and thus stronger selection at the nucleotide level in Drosophila.

Incorporating rate variation into the model allowed us to identify a strong signal of positive selection acting on the inner dynein arm, which could be a result of selection against *Wolbachia* infection in Drosophila. With the model without rate variation, this signal is masked by other, presumably false positive, genes. While there have already been several large-scale analyses of positive selection in Drosophila (Clark et al., 2007; Markova-Raina and Petrov, 2011; Cicconardi et al., 2017), none of them reported positive selection affecting inner dynein arm. An association between positive selection on dynein and *Wolbachia* needs more evidence before it can be confirmed, however it is promising that this association is better supported with a more stringent and realistic model. While the gene list obtained with rate variation is shorter and thus provides less signal for anatomical enrichment, testis notably remains highly significant, as expected for selection on sexual conflict in flies (Supplementary Table S18). Indeed, sexual selection has been repeatedly demonstrated using various approaches (e.g., Lupold et al., 2016).

Both versions of the branch-site test detect a small number of genes under positive selection, relative to expectations from some other approaches (Bierne and Eyre-Walker, 2004). It has been suggested that the presence of positive selection on background branches can cause a decrease in power of the branch-site models (Kosakovsky Pond et al., 2011). However, we observe only a marginal decrease of power in our simulations (see Supplementary Materials, “Branch-site model and background positive selection”). This decrease is small both under the classical branch-site model and under rate variation. Thus, this background selection effect does not seem to explain the small number of genes detected.

An important question is why accounting for rate variation changes the statistical properties of the test. For models with a single *ω* = *d*_*N*_ /*d*_*S*_ value per alignment, comparison between *d*_*N*_ and *d*_*S*_ can be viewed as a contrast between the rates before and after the action of selection on the protein, and should not be significantly biased by nucleotide rate variation (Yang, 2014). However, when *ω* is allowed to vary, *d*_*N*_ /*d*_*S*_ overestimation could be caused not only by the variation in *d*_*N*_, but also by codon-specific substitution rates. Indeed, having a small percentage of rapidly evolving codons in the gene would not be captured by an overall rate for *d*_*S*_, and therefore would be interpreted as positive selection by models with protein level but without nucleotide level rate variation. Conversely, fully accounting for rate variation allows detecting these codons as rapidly evolving by the signatures of both synonymous and nonsynonymous substitutions.

There is recent evidence that double mutations in coding sequences increase the branch-site model false positive rate from 1.1% to 8.6% in similar datasets to those investigated here (Venkat et al., 2018). The interaction between this effect and rate variation along the gene is worth investigating.

We compared three different models accounting for rate variation: the site variation model of Rubinstein et al. (2011), the codon 3-rate variation of Scheffler et al. (2006), and our new codon gamma variation model which extends Scheffler et al. (2006). The codon rate variation model can be informally thought of as a special case of the site rate variation model. Despite that, the codon gamma rate variation performs better both in the simulations (M8: Table 2, branch-site AUC: Supplementary Table S2) and on the vertebrate dataset (positive selection predictions: Supplementary Table S4). There are probably two reasons for that. First, the fact that we can assign a rate to a particular nucleotide position does not necessary mean that we can reliably estimate it. Only two amino acids allow single nucleotide synonymous substitution associated with the first or second codon positions. This means that individual position rates can be estimated mostly through non-synonymous substitutions, which are typically rare compared to synonymous ones. Moreover, the branch-site and M8 models allow variation in the nonsynonymous rate over codon positions, which means estimates of *ω* and site rates are not independent. However, there is no visible dependency between codon rate variation (Fig. 3) and *ω* variation (Supplementary Fig. S15).

Secondly, we expect site rates to be autocorrelated along the sequence, since many factors, such as GC content, recombination rate, or chromatin state change slowly over the gene. Indeed we see a signal of such autocorrelation in our data (effect size 0.018, *p*-value < 0.0001). Therefore having an independent rate for every site is probably redundant.

Statistical performance of codon gamma rate variation and codon 3-rate variation by Scheffler et al. (2006) is comparable (M8 based models: Table 2, branch-site based models: Supplementary Table S2). Also, they are similar in terms of the model support provided by the data. However, codon gamma rate variation provides two important advantages. First, it is up to two times faster than codon 3-rate variation implemented using the same optimizations in Godon (Table 3). This is probably caused by a larger dimensionality of parameter space and by non-independence between model parameters, which can slow down the likelihood optimization. Second, comparison of positive selection predictions between codon gamma variation and the model with the highest support suggests (Supplementary Table S4) that the codon gamma rate variation model provides a better detection of positive selection even in case of model misspecification.

One of the key advantages of codon variation relative to site variation is computational performance. Having a distinct rate for every position increases the number of site classes for which likelihood computations have to be performed by a factor of *K*^3^, where *K* is the number of discrete categories for gamma distribution. Having a rate only for each codon increases the number of site classes by a factor of *K*. This means that even for four discrete categories, the slowdown of likelihood computation for site rate gamma model will be about 64 times, vs. only 4 times for codon rate variation model. In practice this ratio between the two models was respected in simulated and in vertebrate data. This makes codon rate variation models usable in large phylogenomics datasets, as we demonstrate by analyzing 12 Drosophila genomes.

Unlike traditional mechanistic codon models, our new models allow independent estimations of substitution rate at the nucleotide level and of selective pressure on amino acid sequences. It should be noted that individual site rates estimates may be still noisy because of the amount of data available. But given enough data it is possible to have accurate estimates of selection acting on specific regions, e.g., splicing motifs, within coding sequences (Fig. 3).

## V. Conclusions

We have performed a large-scale comparison of different approaches to model rate variation. Failure to account for rate variation leads to both type I and type II errors. We also propose an extension to the model of Scheffler et al. (2006), which has a good statistical performance both in the presence and in the absence of rate variation. We also provide a software implementation of the new models. Rate variation is strongly supported by homologous genes both from species with larger (flies) and smaller (vertebrates) effective population sizes. We are able to capture differences in substitution rates caused by nucleotide selection. Importantly, while being more complex these models remain computationally tractable and therefore can be applied to large-scale datasets. These models and their efficient implementation open the opportunity of simultaneous analysis of different layers of selection in phylogenomics.

## VI. Methods

### I. Sequence simulations

We simulated eight datasets (Table 1) that include either no rate variation across sites (corresponding to the GY94 model), variation between sites (corresponding to the Rubinstein et al. 2011 model), 3-rate variation between codons (corresponding to Scheffler et al. 2006 parametrization), or gamma variation between codons (corresponding to our new approach). Each dataset contains 1,000 alignments simulated under the null hypothesis H0 with no positive selection (all *ω* ≤ 1) as well as 1,000 alignments under the alternative hypothesis H1 with positive selection (some *ω* > 1). Models used in the study are mainly focusing on detecting alignments (i.e., genes) under positive selection, rather than individual sites. Since our dataset is balanced in terms of alignments, we used ROC and AUC as our main performance metrics. All the datasets had between 8 and 12 sequences composed of 100 to 400 codons and were simulated using our software named cosim (see Availability). The parameters of each simulation, including the alignment length and the number of species, were generated at random from their respective distribution (Supplementary Table S19, Supplementary Fig. S18). Maximum values of *ω* > 1 for H1 were 79 and 15 for the branch-site and M8 models, respectively. Values of *α* were within the range of values estimated from the real data (Supplementary Fig. S19), with an emphasis on smaller values where the variation is stronger. For the simulations including rate variation, we used four discrete gamma categories that we assigned either to sites or to codons. The M8 model assumes that the neutral sites and those under purifying selection have an *ω* drawn from a beta distribution and we represented this distribution using five discrete categories. Finally, to simulate evolution under the branch-site model, we randomly selected one “foreground” branch of the phylogenetic tree (either internal or terminal) for every simulated alignment.

### II. Vertebrate and Drosophila datasets

We analyzed two biological datasets. Our goals were to compare the fit of the different models on real data, and to study which gene features are contributing to the variation of the substitution rate. First, we used a vertebrate one-to-one orthologs dataset (Studer et al. 2008, available at http://bioinfo.unil.ch/supdata/positiveselection/Singleton.html) consisting of 767 genes (singleton dataset). This dataset was already used in previous studies of codon models (Fletcher and Yang, 2010; Gharib and Robinson-Rechavi, 2013; Davydov et al., 2017).

We also used a subset of one-to-one orthologs from 12 Drosophila species from the Selectome database (release 6, http://selectome.unil.ch/). This dataset consists of 8,606 genes, and the alignments were filtered to remove unreliably aligned codons; the Selectome filtering procedure is based notably on GUIDANCE (Penn et al., 2010) and is described on the Selectome web-site and the corresponding publication (Moretti et al., 2014). Phylogenetic trees for Drosophila and vertebrates were acquired from TimeTree (Hedges et al., 2006).

### III. Model parameters inference

For all the tests on simulated data we used the correct (i.e., simulated) tree topology, but starting branch lengths were estimated using PhyML v. 20131022 (Guindon et al., 2010) with the model HKY85 (Hasegawa et al., 1985). We did not start the optimization from the true branch lengths, by similarity to a real use-case, when only gene sequences are available, and the true branch lengths are unknown. Additionally, we also show results of estimations when true branch lengths were used. While tree topology is also inferred in real use-cases, and wrong topology could impact the inference of positive selection (Diekmann and Pereira-Leal, 2015), investigating this is outside the scope of our study.

Optimization of all model parameters jointly with branch lengths is not practical and substantially increases the computational load. We instead first estimated branch lengths using the simpler M0 model, which assumes a constant *ω* across branches and sites, and optimized in a second step the model parameters of the M8 or branch-site models with or without rate variation, while fixing branch lengths. A similar approach was used in previous studies (Scheffler et al., 2006; Moretti et al., 2014).

We show that this approach at least in the case of the absence of variation does not decrease significantly the statistical properties of the positive selection inference (Supplementary Fig. S20).

All model optimizations with the exception of BUSTED were performed in Godon, followed by model selection (see below).

For BUSTED we used an implementation available in HyPhy v. 2.2.6 (Pond et al., 2005). When running BUSTED on M8 simulations, positive selection was tested on all the branches jointly. In the case of branch-site model simulations, only the foreground branch (*ω* ≥ 1) was tested for positive selection.

For the biological datasets, all the internal branches were tested using the branch-site model for positive selection. Tip branches were not tested to reduce the potential effect of sequencing errors. The M8 model was applied to estimate substitution rates and *ω* for individual sites.

### IV. Model selection

During model selection we had eight models to choose from: four rate variation approaches and, for each, the absence or presence of positive selection. Although LRT can be used to test for positive selection, it is not possible to use it to compare across all eight models that we tested (i.e., any pair of codon rate variation and site rate variation models cannot be represented as a nested pair).

We thus first used the AIC on the alternative model to select one of the four approaches to model rate variation: no rate variation, site rate variation, codon 3-rate variation, or codon gamma rate variation. For the Drosophila dataset, when only the no rate variation and codon gamma rate variation models were compared, we used both AIC and LRT. Fig. 5 shows the scheme of model selection.

**Figure 5:**
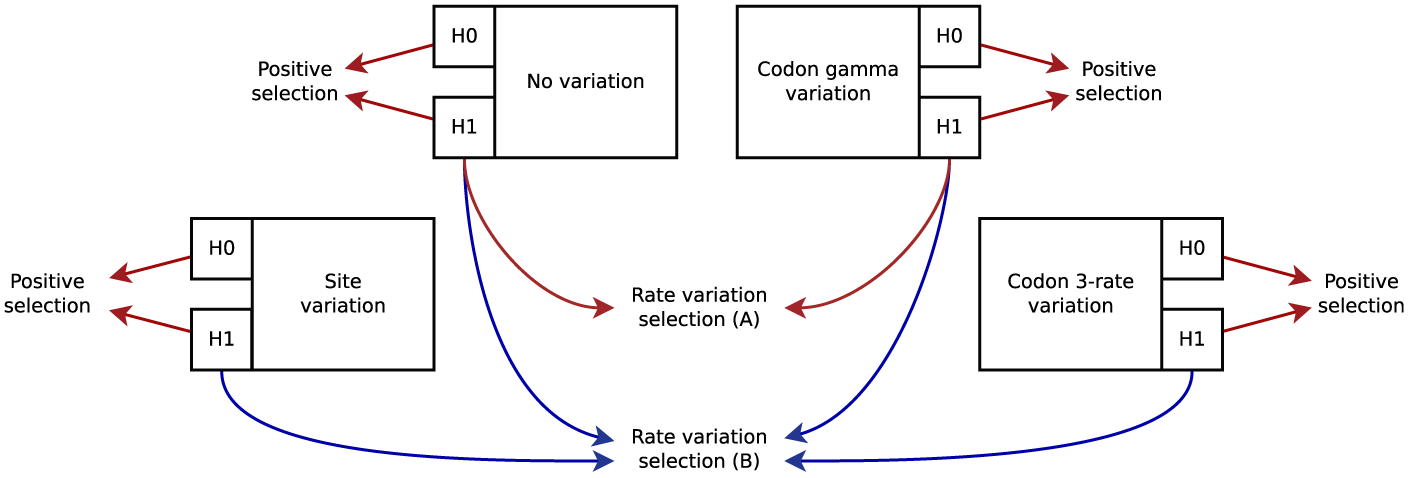
Model selection scheme. Red arrows correspond to LRT and blue arrows correspond to AIC. Rate variation selection (A) was performed only on the full Drosophila dataset, while (B) was used on all three datasets: the vertebrate dataset, the Drosophila dataset, and a subset of 1,000 genes from the Drosophila dataset.

Once the rate variation model was selected, we performed LRT to detect positive selection on the corresponding pair of models, i.e., model with *ω* ≤ 1 and model without this constraint. A 50:50 mix of a *χ*^2^ distribution with one degree of freedom and of a point mass of 0 was used as a null distribution (Yang and dos Reis, 2011). False discovery rate was computed using the qvalue package (Storey et al., 2004).

### V. Posterior rates inference

In order to estimate substitution rates for individual codons we used an approach similar to Mayrose et al. (2007) and Rubinstein et al. (2011). First, we estimated the probability of a codon belonging to each rate as *P* = *Pr*(*ρ*^(*m*,*)^ = *ρ*_*k*_ |*x*_*m*_, *η*), where *ρ*^(*m*,*)^ is the rate of codon *m, ρk* is the *k*-th discrete gamma rate, *x*_*m*_ is the data observed at codon *m*, and *η* are the parameters of the model (e.g., for M8 *η* = {*p*_0_, *p, q, ω*}). In this approach, *η* is replaced with the maximum likelihood estimate of model parameters 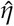. Thus codon rates can be estimated as a weighted sum

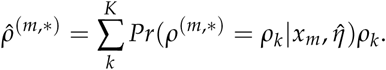

An alternative would be to use Bayes empirical Bayes (BEB, Yang et al., 2005), instead. However, BEB was developed and tested for site detection in particular codon models, and we do not know how well it is applicable to rate variation. On top of that given the increased parametric space of the model, BEB would be computationally intensive. Since we are averaging rates over multiple sites, random noise should not introduce a substantial bias.

Codon site *d*_*N*_ /*d*_*S*_ ratios in the M8 model can be estimated using a similar approach, while replacing codon rate categories with the *ω* categories.

Posterior codon rate and *d*_*N*_ /*d*_*S*_ estimation is implemented in Godon. In all cases we used an alternative model for posterior estimation. Since the null model for every pair is a special case of the alternative, we can use the later for parameter estimation without any significant loss of precision.

To test for autocorrelation, we used the average absolute difference between the posterior rates of the neighboring codon positions as a statistic:

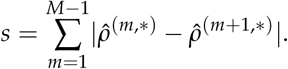

The null distribution was computed by shuffling the rates within all the genes 10,000 times.

### VI. Enrichment analysis

GO-enrichment analysis was performed using the topGO package (Alexa and Rahnenfuhrer, 2016). We used Fisher’s exact test and the graph decorrelation technique named weight01. We used TopAnat (https://bgee.org/?page=top_anat#/) from Bgee 14.0 (Komljenovic et al., 2018) for expression enrichment analysis.

### VII. Regression analysis

To estimate dependencies between various parameters (variables) we used linear models (lm function, R version 3.5.1). Variables were transformed to have a bell shaped distribution if possible (see Supplementary Table S20, Supplementary Fig. S21). Subsequently, parameters were centered at zero and scaled so that standard deviation was equal to one. This transformation allowed us to compare the estimates of the effects. Because in some cases residuals showed strong heteroscedasticity (Supplementary Fig. S22, S23 for vertebrates and Drosophila), we used White standard errors (White, 1980) implemented in the `sandwich` R package.

We used expression data for *H. sapiens* from Fagerberg et al. (2014), acquired from Kryuchkova-Mostacci and Robinson-Rechavi (2015). For *D. melanogaster* we used data from Li et al. (2014), available at http://www.stat.ucla.edu/∼jingyi.li/software-and-data.html. Recombination rates for genes were computed using Recombination Rate Calculator (ver. 2.3, Fiston-Lavier et al., 2010) using dataset from Comeron et al. (2012).

The relative support of codon gamma rate model was computed as a log ratio between Akaike weights (Wagenmakers and Farrell, 2004) of the model with codon gamma rate variation and the model without rate variation.

### VIII. Availability

All the code and the full lists of detected genes are available from https://bitbucket.org/Davydov/codon.rate.variation. Sequence simulator cosim is available from http://bitbucket.org/Davydov/cosim. Codon model parameter estimator Godon is available from https://bitbucket.org/Davydov/godon. We provide source code as well as precompiled binaries for GNU/Linux (64bit).

To our best knowledge Godon provides the first implementation of branch-site models (Zhang et al., 2005) incorporating codon rate variation using approaches of Scheffler et al. (2006), Rubinstein et al. (2011), and our own gamma rates extension.

#### Godon usage

Godon provides an easy way to perform an analysis given a FASTA-file and a newick-file. E.g., the following command will perform branch-site test for positive selection with gamma distributed codon rates on every branch of a tree:

~~~
godon test BSG –all-branches–ncat-codon-rate 4 input.fst input.nwk
~~~

Often branch length optimization is performed once using a simpler model and then for every test the branch length parameters are fixed. In Godon there is an easy way to achieve this behavior:

~~~
godon test BSG –m0-tree –all-branches–ncat-codon-rate 4 input.fst input.nwk
~~~

## VII. Acknowledgments

We would like to thank Peer Community In Evolutionary Biology (https://evolbiol.peercommunityin.org/), Julien Yann Dutheil, and David Enard for useful comments. We also thank Evgeniy Riabenko for helpful suggestions regarding linear modeling. This work was supported by the Swiss National Science Foundation (grant number IZLRZ3_163872). The computations were performed at the Vital-IT (http://www.vital-it.ch) center for high-performance computing of the Swiss Institute of Bioinformatics.

## References

Alexa, A. and Rahnenfuhrer, J. topGO: Enrichment Analysis for Gene Ontology, 2016. R package version 2.28.0.

Anisimova, M., Nielsen, R. and Yang, Z. Effect of recombination on the accuracy of the likelihood method for detecting positive selection at amino acid sites. Genetics, 164(3): 1229–1236, Jul 2003.

Baele, G. and Lemey, P. Bayesian evolutionary model testing in the phylogenomics era: matching model complexity with computational efficiency. Bioinformatics, 29(16):1970–1979, Aug 2013.

Bentele, K. et al. Efficient translation initiation dictates codon usage at gene start. Mol. Syst. Biol., 9:675, Jun 2013.

Betancur-R, R., Orti, G. and Pyron, R.A. Fossil-based comparative analyses reveal ancient marine ancestry erased by extinction in ray-finned fishes. Ecol. Lett., 18(5):441–450, May 2015.

Bierne, N. and Eyre-Walker, A. The genomic rate of adaptive amino acid substitution in Drosophila. Mol. Biol. Evol., 21(7):1350–1360, Jul 2004.

Boyko, A.R. et al. Assessing the evolutionary impact of amino acid mutations in the human genome. PLoS Genet., 4(5):e1000083, May 2008.

Bulmer, M. The selection-mutation-drift theory of synonymous codon usage. Genetics, 129 (3):897–907, Nov 1991.

Campos, J.L. et al. The relation between recombination rate and patterns of molecular evolution and variation *in Drosophila melanogaster*. Mol. Biol. Evol., 31(4):1010–1028, Apr 2014.

Carlini, D.B. and Stephan, W. In vivo introduction of unpreferred synonymous codons into the Drosophila Adh gene results in reduced levels of ADH protein. Genetics, 163 (1):239–243, Jan 2003.

Cartegni, L., Chew, S.L. and Krainer, A.R. Listening to silence and understanding non-sense: exonic mutations that affect splicing. Nat. Rev. Genet., 3(4):285–298, Apr 2002.

Castellano, D., James, J. and Eyre-Walker, A. Nearly Neutral Evolution across the *Drosophila melanogaster* Genome. Mol. Biol. Evol., 35(11):2685–2694, Nov 2018.

Chamary, J.V., Parmley, J.L. and Hurst, L.D. Hearing silence: non-neutral evolution at synonymous sites in mammals. Nat. Rev. Genet., 7(2):98–108, Feb 2006.

Cicconardi, F. et al. Positive diversifying selection is a pervasive adaptive force throughout the Drosophila radiation. Mol. Phylogenet. Evol., 112:230–243, Jul 2017.

Clark, A.G. et al. Evolution of genes and genomes on the Drosophila phylogeny. Nature, 450(7167):203–218, Nov 2007.

Comeron, J.M. Selective and mutational patterns associated with gene expression in humans: influences on synonymous composition and intron presence. Genetics, 167(3): 1293–1304, Jul 2004.

Comeron, J.M., Ratnappan, R. and Bailin, S. The many landscapes of recombination in *Drosophila melanogaster*. PLoS Genet., 8(10): e1002905, 2012.

Daub, J.T. et al. Detection of Pathways Affected by Positive Selection in Primate Lineages Ancestral to Humans. Mol. Biol. Evol., 34(6): 1391–1402, Jun 2017.

Davydov, I.I., Robinson-Rechavi, M. and Salamin, N. State aggregation for fast likelihood computations in molecular evolution. Bioinformatics, 33(3):354–362, 02 2017.

Diekmann, Y. and Pereira-Leal, J.B. Gene Tree Affects Inference of Sites Under Selection by the Branch-Site Test of Positive Selection. Evol. Bioinform. Online, 11(Suppl 2): 11–17, 2015.

Dimitrieva, S. and Anisimova, M. Unraveling patterns of site-to-site synonymous rates variation and associated gene properties of protein domains and families. PLoS ONE, 9 (6):e95034, 2014.

Drummond, D.A. and Wilke, C.O. Mistranslation-induced protein misfolding as a dominant constraint on coding-sequence evolution. Cell, 134(2):341–352, Jul 2008.

Drummond, D.A. et al. Why highly expressed proteins evolve slowly. Proc. Natl. Acad. Sci. U.S.A., 102(40):14338–14343, Oct 2005.

Duret, L. and Galtier, N. Biased gene conversion and the evolution of mammalian genomic landscapes. Annu Rev Genomics Hum Genet, 10:285–311, 2009.

Fagerberg, L. et al. Analysis of the human tissue-specific expression by genome-wide integration of transcriptomics and antibody-based proteomics. Mol. Cell Proteomics, 13(2): 397–406, Feb 2014.

Fiston-Lavier, A.S. et al. *Drosophila melanogaster* recombination rate calculator. Gene, 463(1-2): 18–20, Sep 2010.

Fletcher, W. and Yang, Z. The effect of insertions, deletions, and alignment errors on the branch-site test of positive selection. Mol. Biol. Evol., 27(10):2257–2267, Oct 2010.

Fryxell, K.J. and Zuckerkandl, E. Cytosine deamination plays a primary role in the evolution of mammalian isochores. Mol. Biol. Evol., 17(9):1371–1383, Sep 2000.

Fullerton, S.M., Bernardo Carvalho, A. and Clark, A.G. Local rates of recombination are positively correlated with GC content in the human genome. Mol. Biol. Evol., 18(6): 1139–1142, Jun 2001.

Gharib, W.H. and Robinson-Rechavi, M. The branch-site test of positive selection is surprisingly robust but lacks power under synonymous substitution saturation and variation in GC. Mol. Biol. Evol., 30(7):1675–1686, Jul 2013.

Gil, M. et al. CodonPhyML: fast maximum likelihood phylogeny estimation under codon substitution models. Mol. Biol. Evol., 30(6): 1270–1280, Jun 2013.

Glemin, S. et al. Quantification of GC-biased gene conversion in the human genome. Genome Res., 25(8):1215–1228, Aug 2015.

Goldman, N. and Yang, Z. A codon-based model of nucleotide substitution for protein-coding DNA sequences. Mol. Biol. Evol., 11 (5):725–736, Sep 1994.

Gossmann, T.I., Keightley, P.D. and Eyre-Walker, A. The effect of variation in the effective population size on the rate of adaptive molecular evolution in eukaryotes. Genome Biol Evol, 4(5):658–667, 2012.

Guharoy, M. and Chakrabarti, P. Conservation and relative importance of residues across protein-protein interfaces. Proc. Natl. Acad. Sci. U.S.A., 102(43):15447–15452, Oct 2005.

Guindon, S. et al. New algorithms and methods to estimate maximum-likelihood phylogenies: assessing the performance of PhyML 3.0. Syst. Biol., 59(3):307–321, May 2010.

Hasegawa, M., Kishino, H. and Yano, T. Dating of the human-ape splitting by a molecular clock of mitochondrial DNA. J. Mol. Evol., 22 (2):160–174, 1985.

Hedges, S.B., Dudley, J. and Kumar, S. Time-Tree: a public knowledge-base of divergence times among organisms. Bioinformatics, 22 (23):2971–2972, Dec 2006.

Hellmann, I. et al. A neutral explanation for the correlation of diversity with recombination rates in humans. Am. J. Hum. Genet., 72(6): 1527–1535, Jun 2003.

Hellmann, I. et al. Population genetic analysis of shotgun assemblies of genomic sequences from multiple individuals. Genome Res., 18 (7):1020–1029, Jul 2008.

Hodgkinson, A. and Eyre-Walker, A. Variation in the mutation rate across mammalian genomes. Nat. Rev. Genet., 12(11):756–766, Oct 2011.

Hwang, D.G. and Green, P. Bayesian Markov chain Monte Carlo sequence analysis reveals varying neutral substitution patterns in mammalian evolution. Proc. Natl. Acad. Sci. U.S.A., 101(39):13994–14001, Sep 2004.

Jørgensen, F.G. and Schierup, M.H. Increased rate of human mutations where DNA and RNA polymerases collide. Trends Genet., 25 (12):523–527, Dec 2009.

Keightley, P.D., Lercher, M.J. and Eyre-Walker, A. Evidence for widespread degradation of gene control regions in hominid genomes. PLoS Biol., 3(2):e42, Feb 2005.

Keightley, P.D. et al. Analysis of the genome sequences of three *Drosophila melanogaster* spontaneous mutation accumulation lines. Genome Res., 19(7):1195–1201, Jul 2009.

Kertesz, M. et al. Genome-wide measurement of RNA secondary structure in yeast. Nature, 467(7311):103–107, Sep 2010.

Komljenovic, A. et al. BgeeDB, an R package for retrieval of curated expression datasets and for gene list expression localization enrichment tests [version 2; referees 2 approved, 1 approved with reservations]. F1000Research, 5, 2018.

Kong, A. et al. Rate of de novo mutations and the importance of father’s age to disease risk. Nature, 488(7412):471–475, Aug 2012.

Koonin, E.V. and Wolf, Y.I. Constraints and plasticity in genome and molecular-phenome evolution. Nat. Rev. Genet., 11(7): 487–498, Jul 2010.

Kosakovsky Pond, S.L. et al. A random effects branch-site model for detecting episodic diversifying selection. Mol. Biol. Evol., 28(11): 3033–3043, Nov 2011.

Kosiol, C. et al. Patterns of positive selection in six Mammalian genomes. PLoS Genet., 4 (8):e1000144, Aug 2008.

Kryuchkova-Mostacci, N. and Robinson-Rechavi, M. Tissue-Specific Evolution of Protein Coding Genes in Human and Mouse. PLoS ONE, 10(6):e0131673, 2015.

Kudla, G. et al. Coding-sequence determinants of gene expression in *Escherichia coli*. Science, 324(5924):255–258, Apr 2009.

Lartillot, N. and Philippe, H. A Bayesian mixture model for across-site heterogeneities in the amino-acid replacement process. Mol. Biol. Evol., 21(6):1095–1109, Jun 2004.

Leffler, E.M. et al. Multiple instances of ancient balancing selection shared between humans and chimpanzees. Science, 339(6127):1578–1582, Mar 2013.

Lercher, M.J. and Hurst, L.D. Human SNP variability and mutation rate are higher in regions of high recombination. Trends Genet., 18(7):337–340, Jul 2002.

Li, J.J. et al. Comparison of *D. melanogaster* and *C. elegans* developmental stages, tissues, and cells by modENCODE RNA-seq data. Genome Res., 24(7):1086–1101, Jul 2014.

Lupold, S. et al. How sexual selection can drive the evolution of costly sperm ornamentation. Nature, 533(7604):535–538, 05 2016.

Majewski, J. and Ott, J. Distribution and characterization of regulatory elements in the human genome. Genome Res., 12(12):1827–1836, Dec 2002.

Marais, G., Mouchiroud, D. and Duret, L. Neutral effect of recombination on base composition in Drosophila. Genet. Res., 81(2):79–87, Apr 2003.

Markova-Raina, P. and Petrov, D. High sensitivity to aligner and high rate of false positives in the estimates of positive selection in the 12 Drosophila genomes. Genome Res., 21(6): 863–874, Jun 2011.

Mattick, J.S. and Makunin, I.V. Non-coding RNA. Hum. Mol. Genet., 15 Spec No 1:17–29, Apr 2006.

Mayrose, I. et al. Towards realistic codon models: among site variability and dependency of synonymous and non-synonymous rates. Bioinformatics, 23(13):i319–327, Jul 2007.

Moretti, S. et al. Selectome update: quality control and computational improvements to a database of positive selection. Nucleic Acids Res., 42(Database issue):D917–921, Jan 2014.

Murrell, B. et al. Gene-wide identification of episodic selection. Mol. Biol. Evol., 32(5):1365–1371, May 2015.

Muse, S.V. and Gaut, B.S. A likelihood approach for comparing synonymous and non-synonymous nucleotide substitution rates, with application to the chloroplast genome. Mol. Biol. Evol., 11(5):715–724, Sep 1994.

Pál, C., Papp, B. and Hurst, L.D. Highly expressed genes in yeast evolve slowly. Genetics, 158(2):927–931, Jun 2001.

Pal, C., Papp, B. and Lercher, M.J. An integrated view of protein evolution. Nat. Rev. Genet., 7(5):337–348, May 2006.

Penn, O. et al. GUIDANCE: a web server for assessing alignment confidence scores. Nucleic Acids Res., 38(Web Server issue):W23–28, Jul 2010.

Plotkin, J.B. and Kudla, G. Synonymous but not the same: the causes and consequences of codon bias. Nat. Rev. Genet., 12(1):32–42, Jan 2011.

Pond, S.K. and Muse, S.V. Site-to-site variation of synonymous substitution rates. Mol. Biol. Evol., 22(12):2375–2385, Dec 2005.

Pond, S.L., Frost, S.D. and Muse, S.V. HyPhy: hypothesis testing using phylogenies. Bioinformatics, 21(5):676–679, Mar 2005.

Pop, C. et al. Causal signals between codon bias, mRNA structure, and the efficiency of translation and elongation. Mol. Syst. Biol., 10:770, Dec 2014.

Ptak, S.E. et al. Fine-scale recombination patterns differ between chimpanzees and humans. Nat. Genet., 37(4):429–434, Apr 2005.

Ratnakumar, A. et al. Detecting positive selection within genomes: the problem of biased gene conversion. Philos. Trans. R. Soc. Lond., B, Biol. Sci., 365(1552):2571–2580, Aug 2010.

Rocha, E.P. and Danchin, A. An analysis of determinants of amino acids substitution rates in bacterial proteins. Mol. Biol. Evol., 21(1): 108–116, Jan 2004.

Romiguier, J. et al. Comparative population genomics in animals uncovers the determinants of genetic diversity. Nature, 515(7526): 261–263, Nov 2014.

Roux, J., Liu, J. and Robinson-Rechavi, M. Selective Constraints on Coding Sequences of Nervous System Genes Are a Major Determinant of Duplicate Gene Retention in Vertebrates. Mol. Biol. Evol., 34(11):2773–2791, Nov 2017.

Rubinstein, N.D. and Pupko, T. Detection and analysis of conservation at synonymous sites. Codon Evolution: Mechanisms and Models, pages 218–228, 2012.

Rubinstein, N.D. et al. Evolutionary models accounting for layers of selection in protein-coding genes and their impact on the inference of positive selection. Mol. Biol. Evol., 28 (12):3297–3308, Dec 2011.

Rudolph, K.L. et al. Codon-Driven Translational Efficiency Is Stable across Diverse Mammalian Cell States. PLoS Genet., 12(5): e1006024, May 2016.

Russo, C.A. et al. Phylogenetic analysis and a time tree for a large drosophilid data set (diptera: Drosophilidae). Zoological Journal of the Linnean Society, 169(4):765–775, 2013.

Scheffler, K., Martin, D.P. and Seoighe, C. Robust inference of positive selection from recombining coding sequences. Bioinformatics, 22(20):2493–2499, Oct 2006.

Segurel, L., Wyman, M.J. and Przeworski, M. Determinants of mutation rate variation in the human germline. Annu Rev Genomics Hum Genet, 15:47–70, 2014.

Serbus, L.R. and Sullivan, W. A cellular basis for *Wolbachia* recruitment to the host germline. PLoS Pathog., 3(12):e190, Dec 2007.

Spielman, S.J., Wan, S. and Wilke, C.O. A Comparison of One-Rate and Two-Rate Inference Frameworks for Site-Specific dN/dS Estimation. Genetics, 204(2):499–511, Oct 2016.

Stamatoyannopoulos, J.A. et al. Human mutation rate associated with DNA replication timing. Nat. Genet., 41(4):393–395, Apr 2009.

Storey, J.D., Taylor, J.E. and Siegmund, D. Strong control, conservative point estimation and simultaneous conservative consistency of false discovery rates: a unified approach. Journal of the Royal Statistical Society: Series B (Statistical Methodology), 66(1):187–205, 2004.

Studer, R.A. et al. Pervasive positive selection on duplicated and nonduplicated vertebrate protein coding genes. Genome Res., 18(9): 1393–1402, Sep 2008.

Supek, F. and Lehner, B. Differential DNA mismatch repair underlies mutation rate variation across the human genome. Nature, 521 (7550):81–84, May 2015.

Venkat, A., Hahn, M.W. and Thornton, J.W. Multinucleotide mutations cause false inferences of lineage-specific positive selection. Nat Ecol Evol, 2(8):1280–1288, Aug 2018.

Wagenmakers, E.J. and Farrell, S. AIC model selection using Akaike weights. Psychonomic bulletin & review, 11(1):192–196, 2004.

Werren, J.H., Baldo, L. and Clark, M.E. *Wolbachia*: master manipulators of invertebrate biology. Nat. Rev. Microbiol., 6(10):741–751, Oct 2008.

White, H. A heteroskedasticity-consistent co-variance matrix estimator and a direct test for heteroskedasticity. Econometrica: Journal of the Econometric Society, pages 817–838, 1980.

Yang, J.R., Zhuang, S.M. and Zhang, J. Impact of translational error-induced and error-free misfolding on the rate of protein evolution. Mol. Syst. Biol., 6:421, Oct 2010.

Yang, Z. Molecular evolution: a statistical approach, page 61. Oxford University Press, 2014.

Yang, Z. and Bielawski, J.P. Statistical methods for detecting molecular adaptation. Trends Ecol. Evol. (Amst.), 15(12):496–503, Dec 2000.

Yang, Z. and dos Reis, M. Statistical properties of the branch-site test of positive selection. Mol. Biol. Evol., 28(3):1217–1228, Mar 2011.

Yang, Z. et al. Codon-substitution models for heterogeneous selection pressure at amino acid sites. Genetics, 155(1):431–449, May 2000.

Yang, Z., Wong, W.S. and Nielsen, R. Bayes empirical bayes inference of amino acid sites under positive selection. Mol. Biol. Evol., 22 (4):1107–1118, Apr 2005.

Zhang, G. et al. Comparative genomics reveals insights into avian genome evolution and adaptation. Science, 346(6215):1311–1320, Dec 2014.

Zhang, J., Nielsen, R. and Yang, Z. Evaluation of an improved branch-site likelihood method for detecting positive selection at the molecular level. Mol. Biol. Evol., 22(12): 2472–2479, Dec 2005.

