## Supplementary Material for "Large-Scale Comparative Analysis of Codon Models Accounting for Protein and Nucleotide Selection"

### Contents

**1 Comparison of M8 and branch-site models** **3**
1.1 M8 . . . 3
1.2 Branch-site model . . . 5

**2 Branch-site model and background positive selection** **7**

### 1 Comparison of M8 and branch-site models

We used simulations to compare the performance of M8 and branch-site models. First, we were interested if rate variation in the data inflates the false positive rate. Second, how sensitive is the model M8 in the case of data simulated using the branch-site model, i.e., if positive selection affects only a single branch. Third, if we can detect positive selection in the data simulated under the M8 model by testing individual branches with the branch-site model. In other words, if selection affecting all branches can be identified by testing branches one by one.

We used the same dataset as described in the Methods section of the main text. For the branch-site model, we tested all branches one by one and integrated results using either Fisher's method or minimum $p$-value.

## 1.1 M8

The M8 model is sensitive to variation in the substitution rate (Fig. 1). This result is similar to the behavior of the model M8 under M8 simulations (Fig. S2); false positive rate inflation is especially strong in case of codon gamma rate variation. The performance of M8 when detecting short episodes of positive selection is extremely low (Fig. 2). Thus it is not recommended to use M8 if you expect positive selection to be episodic.

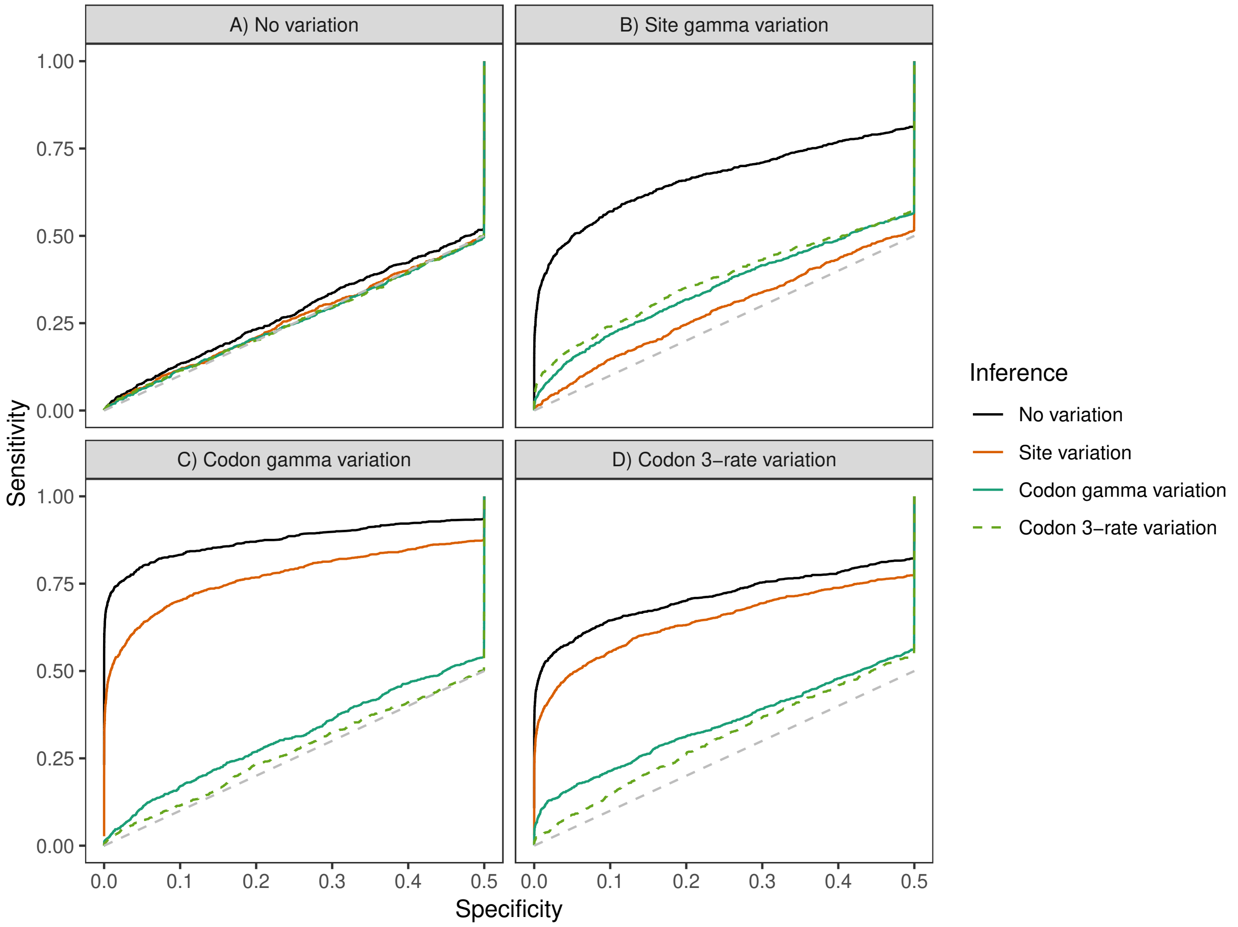

Figure 1: False positive rate as a function of significance level for four M8-based models (M8 with no rate variation, M8 with site rate variation, M8 with codon gamma rate variation, and M8 with codon 3-rate variation) on datasets simulated with the branch-site model A) without rate variation, B) with site rate variation, C) with codon gamma rate variation, and D) with codon 3-rate variation. The dashed diagonal line corresponds to the expected false-positive rate under the null hypothesis.

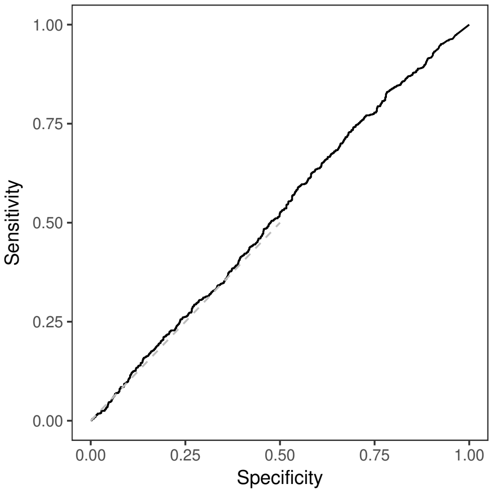

Figure 2: Receiver operating characteristic (ROC) of the model M8 (no rate variation) on simulations performed using branch-site model (no rate variation).

#### 1.2 Branch-site model

The false positive rate of the branch-site test is not affected by rate variation (Fig. 3). Similarly, the branch-site test was less affected by rate variation in the branch-site based simulations (Fig. S7).

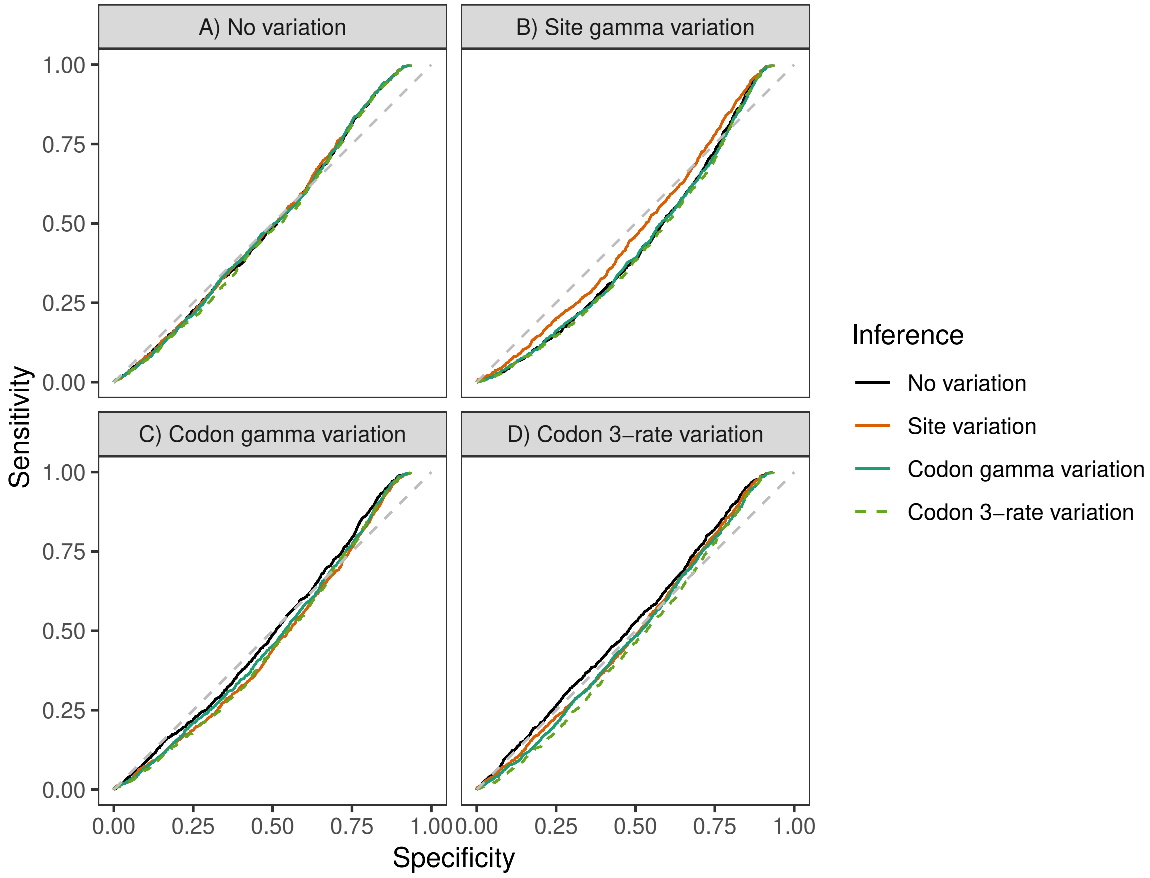

Figure 3: False positive rate as a function of significance level for four branch-site based models (branch-site model with no rate variation, branch-site model with site rate variation, branch-site model with codon gamma variation, and branch-site model with codon 3-rate rate variation) on datasets simulated under the M8 model A) without rate variation, B) with site rate variation, C) with codon gamma rate variation, and D) with codon 3-rate variation. The dashed diagonal line corresponds to the expected false-positive rate under the null hypothesis. All the branches were tested, and $p$-values were aggregated using Fisher's method.

The branch-site model is able to detect positive selection which is affecting all branches, as in the M8 model (Fig. 4). However, statistical performance is substantially degraded compared to the M8 model. There can be two reasons for this. First, while the branch-site model used this way tests every branch, sites evolving under positive selection do not have to be the same for different individual tests. Therefore the aggregated branch-site test, unlike M8, cannot correctly integrate a weak signal of positive selection coming from the same sites. Second, the M8 model has a more complex distribution of $\omega$ values among the sites compared to the branch-site model. This mismatch to the models' assumptions can also play a role in the statistical performance degradation.

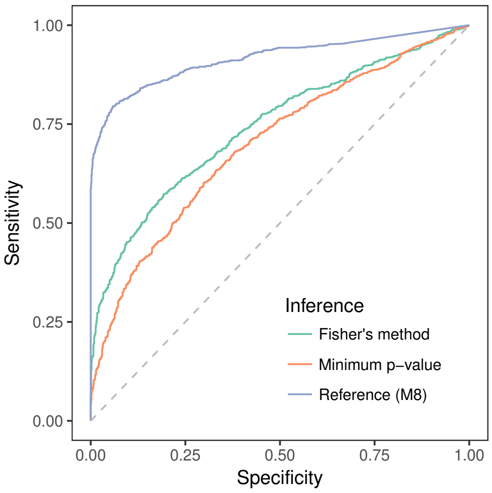

Figure 4: ROC of the branch-site model (no rate variation) on simulations performed using the model M8 (no rate variation). Two $p$-value aggregation methods were used: Fisher's method and minimum $p$-value. Inference using M8 is shown as a reference.

### 2 Branch-site model and background positive selection

We studied the performance of the branch-site model in the presence of strong positive selection along the background branches. For that we chose a single tree (`0176.nwk` in the git repository). We then randomly assigned some background branches to have positive selection ($\omega_3 = 20$) in 30% of alignment codon sites ($p_3 = 0.3$); sites with background positive selection were chosen at random independently for each background branch. The models used for these simulations were an extension of the branch-site model. To simulate a worst-case scenario the foreground positive selection was weaker than background positive selection ($\omega_2 = 10$), with the proportion of sites with foreground positive selection $p_2 = 0.05$. The number of branches with background positive selection was incremented from zero to fifteen (i.e., all background branches). Both the simulations and the inference were performed either with no rate variation or with codon gamma rate variation. For each number of branches with background positive selection and type of rate variation, we performed 1,000 simulations. The inference in all cases was performed on the simulated foreground branch.

We observe that the probability of foreground positive selection detection decreases with the increase in the amount of background positive selection, i.e., with the number of positively selected branches in the background (Fig. 5, 6; Pearson's $r = -0.85$, $p$-value$= 3.7 \cdot 10^{-5}$ and $r = -0.76$, $p$-value$= 0.0007$ for no variation and codon gamma rate variation, respectively). Yet, even though we simulate a large proportion of sites evolving under strong background positive selection, the decrease of the method's power is rather weak, from 56% to 47% and from 37% to 30% in the worst case for no variation and codon gamma rate variation, respectively.

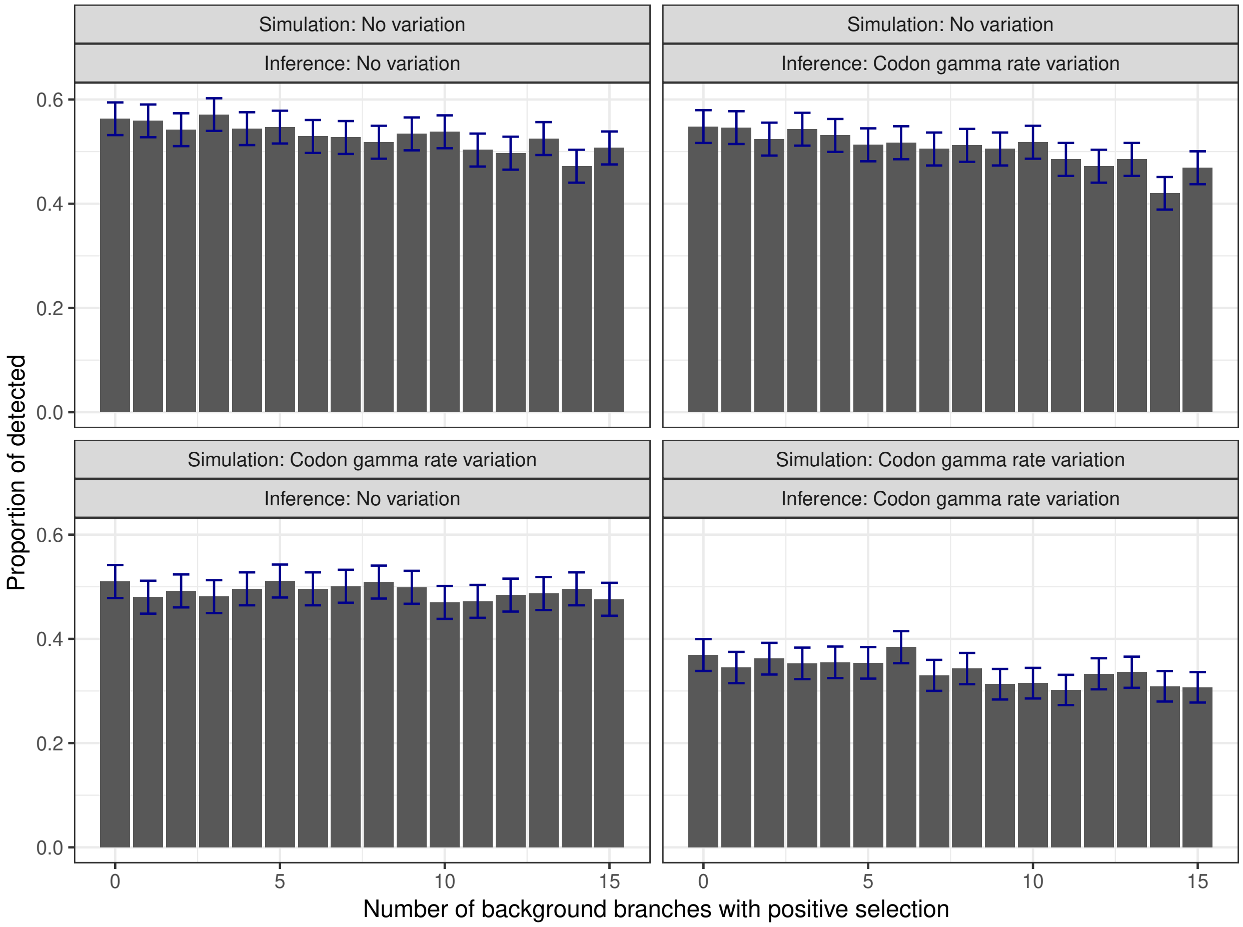

Figure 5: Proportion of simulations with foreground positive selection successfully detected as a function of the number of background branches with positive selection. Each panel corresponds to a combination of rate variation used for simulation and inference. Blue error bars show the 95% confidence intervals for population proportion (normal approximation).

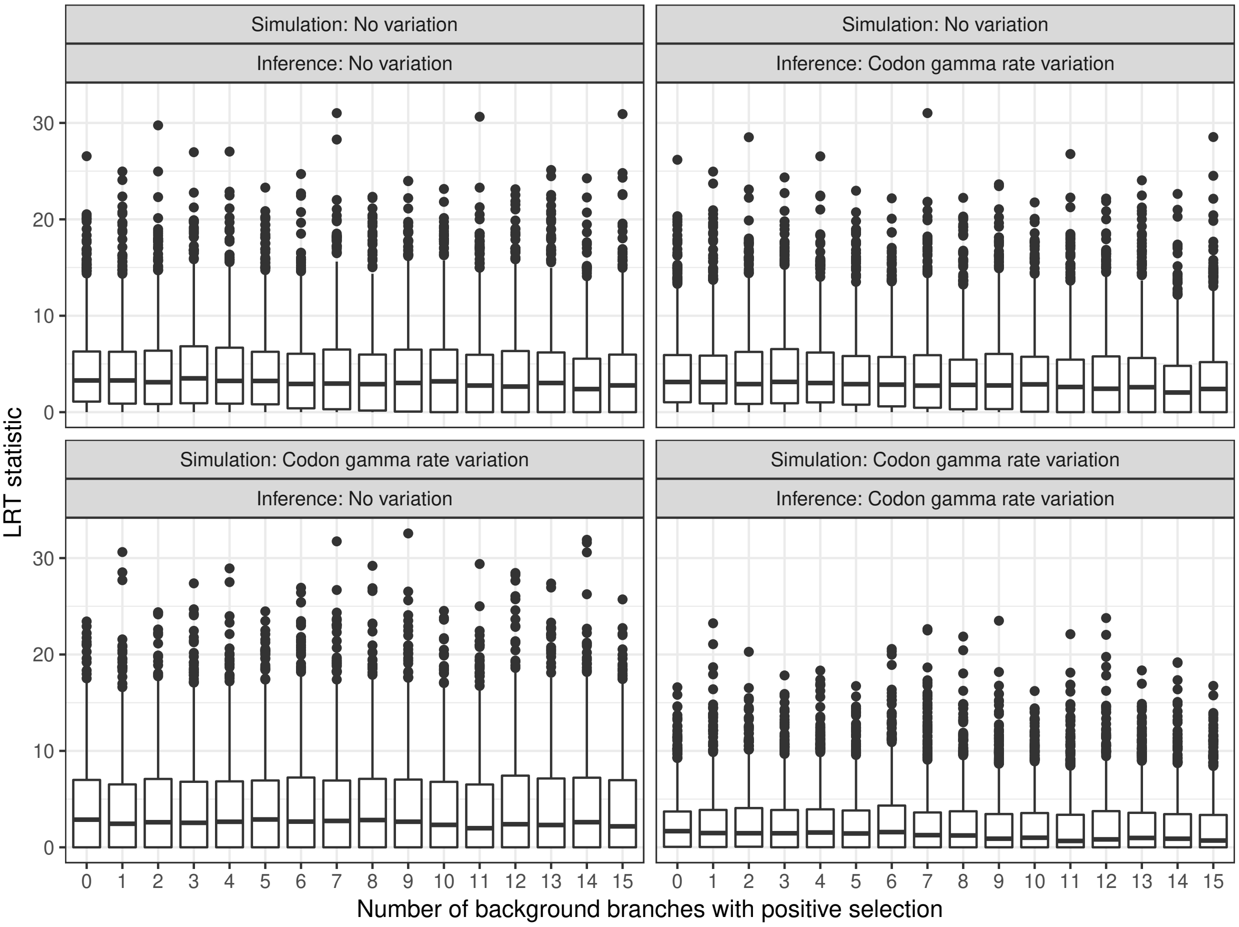

Figure 6: Value of the LRT statistic for the foreground branch as a function of the number of background branches with positive selection. Each panel corresponds to a combination of rate variation used for simulation and inference.

### Tables

| Estimation | Simulation: No variation | Site variation | Codon 3-rate variation | Codon gamma variation |
|---|---|---|---|---|
| No var. | 0.856/ 100% | 0.755/92.8% | 0.704/88.1% | 0.638/77.0% |
| Site var. | 0.852/99.5% | 0.815/ 100% | 0.729/91.2% | 0.694/83.8% |
| Codon 3-rate var. | 0.806/94.2% | 0.775/95.2% | 0.787/98.6% | 0.790/95.4% |
| Codon gamma var. | 0.825/96.3% | 0.800/98.3% | 0.798/ 100% | 0.829/ 100% |
| BUSTED | 0.719/84.0% | 0.669/82.2% | 0.670/80.9% | 0.697/87.3% |

Table S1: Accuracy for all M8-based simulations and for BUSTED. Second number computed as proportion of maximum accuracy for a particular simulation (i.e., maximum value for a column).

| | Simulation | | | |
|---|---|---|---|---|
| Estimation | No variation | Site variation | Codon 3-rate variation | Codon gamma variation |
| No var. | 0.882/99.8% | 0.842/97.2% | 0.863/98.0% | 0.821/94.2% |
| Site var. | 0.880/99.6% | 0.866/ 100% | 0.871/98.9% | 0.833/95.6% |
| Codon 3-rate var. | 0.884/ 100% | 0.841/97.1% | 0.880/ 100% | 0.864/99.2% |
| Codon gamma var. | 0.875/99.0% | 0.847/97.7% | 0.880/ 100% | 0.871/ 100% |

Table S2: Area under curve (AUC) for all branch-site model-based simulations. Second number computed as proportion of maximum AUC for a particular simulation.

| Estimation | Simulation: No variation | Simulation: Site variation | Simulation: Codon 3-rate variation | Simulation: Codon gamma variation |
|---|---|---|---|---|
| No var. | 0.815/99.9% | 0.737/97.4% | 0.795/98.2% | 0.763/95.3% |
| Site var. | 0.815/99.9% | 0.756/ 100% | 0.802/99.1% | 0.777/97.1% |
| Codon 3-rate var. | 0.811/ 99.4% | 0.730/96.4% | 0.804/99.3% | 0.792/98.9% |
| Codon gamma var. | 0.816/ 100% | 0.737/97.4% | 0.809/ 100% | 0.801/ 100% |

Table S3: Accuracy for all branch-site model-based simulations. Second number computed as proportion of maximum accuracy for a particular simulation (i.e., maximum value for a column).

| **A** No variation | Best supported − | Best supported + |
|---|---|---|
| − | 7,062 | 59 |
| + | 817 | 969 |

| **B** Site variation | Best supported − | Best supported + |
|---|---|---|
| − | 7,515 | 70 |
| + | 364 | 958 |

| **C** Codon 3-rate variation | Best supported − | Best supported + |
|---|---|---|
| − | 7,755 | 81 |
| + | 124 | 947 |

| **D** Codon gamma variation | Best supported − | Best supported + |
|---|---|---|
| − | 7,795 | 91 |
| + | 84 | 937 |

Table S4: Positive selection predictions compared to the best supported model for the vertebrate dataset. A) Branch-site without rate variation B) branch-site with site rate variation C) branch-site with codon 3-rate variation D) branch-site with codon gamma rate variation.

| Variable | Estimate | Std. Error | $t$ value | $p$-value |
|---|---|---|---|---|
| Number of sequences | 0.051 | 0.033 | 1.554 | 0.12061 |
| **Total branch length** | 0.123 | 0.028 | 4.339 | $1.66 \cdot 10^{-5}$ |
| **Alignment length** | 0.791 | 0.041 | 19.382 | $< 2 \cdot 10^{-16}$ |
| Length of coding sequence | -0.029 | 0.045 | -0.653 | 0.51371 |
| **GC content (mean)** | 0.175 | 0.028 | 6.227 | $8.53 \cdot 10^{-10}$ |
| **GC content (stdev)** | -0.075 | 0.025 | -3.014 | 0.00268 |
| Total intron length | -0.016 | 0.030 | -0.540 | 0.58940 |
| Number of exons | 0.052 | 0.048 | 1.080 | 0.28057 |
| **Maximum expression** | 0.075 | 0.031 | 2.425 | 0.01558 |
| Mean expression | -0.048 | 0.029 | -1.657 | 0.09803 |

Table S5: Linear model of relative support of model with the codon gamma rate variation; vertebrate dataset. Significant variables ($p$-value $< 0.05$) in bold. Model $p$-value is $< 2.2 \cdot 10^{-16}$, adjusted $R^2$ is 0.6247. Model formula: `Relative model support ~ Number of sequences + Total branch length + Alignment length + Length of coding sequence + GC content (mean) + GC content (stdev) + Total intron length + Number of exons + Maximum expression + Mean expression`.

| Variable | Estimate | Std. Error | $t$ value | $p$-value |
|---|---|---|---|---|
| Number of sequences | 0.101 | 0.064 | 1.578 | 0.11513 |
| Total branch length | 0.013 | 0.044 | 0.299 | 0.76512 |
| Alignment length | 0.044 | 0.050 | 0.874 | 0.38250 |
| Length of coding sequence | 0.025 | 0.061 | 0.415 | 0.67848 |
| **GC content (mean)** | -0.228 | 0.042 | -5.402 | $9.23 \cdot 10^{-8}$ |
| **GC content (stdev)** | 0.197 | 0.038 | 5.176 | $3.01 \cdot 10^{-7}$ |
| Total intron length | 0.004 | 0.035 | 0.110 | 0.91210 |
| Number of exons | 0.019 | 0.064 | 0.293 | 0.76954 |
| **Maximum expression** | -0.115 | 0.043 | -2.672 | 0.00773 |
| Mean expression | 0.037 | 0.043 | 0.850 | 0.39578 |

Table S6: Linear model of $\alpha$ parameter of gamma distribution with the codon rate variation; vertebrate dataset. Significant variables ($p$-value $< 0.05$) in bold. Model $p$-value is $< 2.2 \cdot 10^{-16}$, adjusted $R^2$ is 0.1275. Model formula: `α ~ Number of sequences + Total branch length + Alignment length + Length of coding sequence + GC content (mean) + GC content (stdev) + Total intron length + Number of exons + Maximum expression + Mean expression`.

| significance threshold of 0.05 | | | FDR threshold of 0.1 | | |
|---|---|---|---|---|---|
| **A** | Codon gamma variation | | **B** | Codon gamma variation | |
| No variation | − | + | No variation | − | + |
| − | 6,452 | 36 | − | 7,138 | 2 |
| + | 607 | 580 | + | 494 | 41 |
| **C** | Codon 3-rate variation | | **D** | Codon 3-rate variation | |
| No variation | − | + | No variation | − | + |
| − | 6,449 | 39 | − | 7,140 | 0 |
| + | 558 | 629 | + | 505 | 30 |
| **E** | Site variation | | **F** | Site variation | |
| No variation | − | + | No variation | − | + |
| − | 6,457 | 31 | − | 7,140 | 0 |
| + | 408 | 779 | + | 397 | 138 |
| **G** | Codon gamma variation | | **H** | Codon gamma variation | |
| Site variation | − | + | Site variation | − | + |
| − | 6,822 | 43 | − | 7,533 | 4 |
| + | 237 | 573 | + | 99 | 39 |

Table S7: Positive selection predictions for a random subset (1,000 genes) of the Drosophila dataset. Numbers in each cell indicate how many branches were detected (+) or not detected (-) to be evolving under positive selection by different variants of the branch-site model. A,B no rate variation versus codon gamma rate variation; C,D) no rate variation versus codon 3-rate variation; E,F) no rate variation versus site rate variation; G,H) site rate variation versus codon gamma rate variation; A,C,E,G) significance threshold of 0.05; B,D,F,H) false discovery rate threshold of 0.1.

| A | GO.ID | Term | Annotated | Significant | Expected | $p$-value | $q$-value |
|---|---|---|---|---|---|---|---|
| 1 | GO:0036156 | inner dynein arm | 7 | 3 | 0.17 | 0.00049 | 0.39245 |
| 2 | GO:0005587 | collagen type IV trimer | 2 | 2 | 0.05 | 0.00062 | 0.39245 |
| 3 | GO:0005700 | polytene chromosome | 148 | 9 | 3.69 | 0.00356 | 1.00000 |
| 4 | GO:0016442 | RISC complex | 5 | 2 | 0.12 | 0.00587 | 1.00000 |
| 5 | GO:0016456 | X chromosome located dosage compensation... | 5 | 2 | 0.12 | 0.00587 | 1.00000 |

| B | GO.ID | Term | Annotated | Significant | Expected | $p$-value | $q$-value |
|---|---|---|---|---|---|---|---|
| 1 | GO:0045466 | R7 cell differentiation | 54 | 7 | 1.37 | $4.4 \cdot 10^{-5}$ | 0.29912 |
| 2 | GO:0007605 | sensory perception of sound | 57 | 8 | 1.45 | $8.3 \cdot 10^{-5}$ | 0.29912 |
| 3 | GO:1900182 | positive regulation of protein localization to nucleus | 7 | 3 | 0.18 | 0.00052 | 0.96243 |
| 4 | GO:0007403 | glial cell fate determination | 2 | 2 | 0.05 | 0.00064 | 0.96243 |
| 5 | GO:0036159 | inner dynein arm assembly | 8 | 3 | 0.20 | 0.00082 | 0.96243 |

| C | GO.ID | Term | Annotated | Significant | Expected | $p$-value | $q$-value |
|---|---|---|---|---|---|---|---|
| 1 | GO:0045503 | dynein light chain binding | 13 | 5 | 0.32 | $8.7 \cdot 10^{-6}$ | 0.015925 |
| 2 | GO:0045505 | dynein intermediate chain binding | 14 | 5 | 0.34 | $1.3 \cdot 10^{-5}$ | 0.015925 |
| 3 | GO:0005524 | ATP binding | 462 | 26 | 11.21 | $3.8 \cdot 10^{-5}$ | 0.030900 |
| 4 | GO:0051959 | dynein light intermediate chain binding | 12 | 4 | 0.29 | 0.00014 | 0.085418 |
| 5 | GO:0008569 | ATP-dependent microtubule motor activity, minus-end-directed | 14 | 4 | 0.34 | 0.00028 | 0.133041 |

Table S8: Top GO categories for genes with positive selection detected using the branch-site model with codon rate variation (Drosophila dataset). A) Cellular component; B) Biological process; C) Molecular function.

| A | GO.ID | Term | Annotated | Significant | Expected | $p$-value | $q$-value |
|---|---|---|---|---|---|---|---|
| 1 | GO:0005700 | polytene chromosome | 148 | 85 | 55.04 | $8.2 \cdot 10^{-6}$ | 0.010414 |
| 2 | GO:0035097 | histone methyltransferase complex | 31 | 18 | 11.53 | 0.00036 | 0.229780 |
| 3 | GO:0034388 | Pwp2p-containing subcomplex of 90S preribosome | 6 | 6 | 2.23 | 0.00263 | 1.000000 |
| 4 | GO:0000784 | nuclear chromosome, telomeric region | 9 | 8 | 3.35 | 0.00527 | 1.000000 |
| 5 | GO:0008305 | integrin complex | 5 | 5 | 1.86 | 0.00709 | 1.000000 |

| B | GO.ID | Term | Annotated | Significant | Expected | $p$-value | $q$-value |
|---|---|---|---|---|---|---|---|
| 1 | GO:0055088 | lipid homeostasis | 46 | 31 | 17.16 | 0.00018 | 1 |
| 2 | GO:0043484 | regulation of RNA splicing | 70 | 25 | 26.12 | 0.00102 | 1 |
| 3 | GO:0055114 | oxidation-reduction process | 293 | 130 | 109.32 | 0.00131 | 1 |
| 4 | GO:0000723 | telomere maintenance | 24 | 19 | 8.95 | 0.00147 | 1 |
| 5 | GO:0034976 | response to endoplasmic reticulum stress | 69 | 35 | 25.75 | 0.00187 | 1 |

| C | GO.ID | Term | Annotated | Significant | Expected | $p$-value | $q$-value |
|---|---|---|---|---|---|---|---|
| 1 | GO:0005524 | ATP binding | 462 | 218 | 171.74 | $3.0 \cdot 10^{-6}$ | 0.0072793 |
| 2 | GO:0003682 | chromatin binding | 148 | 79 | 55.01 | $4.2 \cdot 10^{-5}$ | 0.0504813 |
| 3 | GO:0004003 | ATP-dependent DNA helicase activity | 21 | 15 | 7.81 | 0.00089 | 0.7130061 |
| 4 | GO:0000166 | nucleotide binding | 666 | 295 | 247.57 | 0.00401 | 1.0000000 |
| 5 | GO:0045503 | dynein light chain binding | 13 | 10 | 4.83 | 0.00417 | 1.0000000 |

Table S9: Top GO categories for genes with positive selection detected using the branch-site model without rate variation (Drosophila dataset). A) Cellular component; B) Biological process; C) Molecular function.

| | Selectome ID | Gene ID | Gene symbol | Gene name | $\omega_2$ | positions |
|---|---|---|---|---|---|---|
| 1 | EMGT00050000003039.1 | FBgn0035799 | CG14838 | uncharacterized protein | 6.62 | 247, 726, 738, 741, 938, 1000 |
| 2 | EMGT00180000072377.2 | FBgn0013813 | Dhc98D | Dynein heavy chain at 89D | 5.51 | 964, 1184, 1422, 1553, 1828, 2108, 2226, 2311, 2541, 3329, 3646, 4031, 4039, 4565, 4687, 4933 |
| 3 | EMGT00180000072377.2 | FBgn0013813 | Dhc98D | Dynein heavy chain at 89D | 4.29 | 388, 1166, 1181, 1986, 4214, 4762, 5071 |
| 4 | EMGT00180000072377.4 | FBgn0039510 | CG3339 | CG3339 | 8.80 | 716, 1029, 1581, 1707 |
| 5 | EMGT00180000072378.1 | FBgn0013811 | Dhc62B | Dynein heavy chain at 62B | 3.08 | 35, 1967, 2481 |
| 6 | EMGT00180000072378.8 | FBgn0013810 | Dhc36C | Dynein heavy chain at 36C | 4.46 | 27, 227, 242, 443, 488, 521, 822, 848, 1337, 1527, 1980, 2939, 2946, 3668, 4118 |

Table S10: A list of genes under positive selection detected by the branch-site model with codon gamma rate variation (Drosophila dataset). Amino acid positions with posterior probability $> 0.9$ are specified as in Selectome release 6 alignments.

| Variable | Estimate | Std. Error | $t$ value | $p$-value |
|---|---|---|---|---|
| **Number of sequences** | -0.291 | 0.022 | -13.193 | $< 2 \cdot 10^{-16}$ |
| **Total branch length** | 0.103 | 0.022 | 4.644 | $3.48 \cdot 10^{-6}$ |
| Alignment length | -0.125 | 0.071 | -1.772 | 0.0765 |
| Length of coding sequence | 0.092 | 0.068 | 1.355 | 0.1755 |
| **GC content (mean)** | -0.106 | 0.017 | -6.247 | $4.39 \cdot 10^{-10}$ |
| GC content (stdev) | -0.023 | 0.013 | -1.760 | 0.0784 |
| **Total intron length** | -0.133 | 0.019 | -6.995 | $2.87 \cdot 10^{-12}$ |
| Number of exons | -0.000 | 0.021 | -0.015 | 0.9883 |
| Maximum expression | 0.008 | 0.030 | 0.265 | 0.7910 |
| Mean expression | -0.024 | 0.029 | -0.823 | 0.4103 |
| **Recombination rate** | -0.052 | 0.010 | -5.054 | $4.43 \cdot 10^{-7}$ |

Table S11: Linear model of $\alpha$ parameter of gamma distribution with the codon rate variation; Drosophila dataset. Significant variables ($p$-value $< 0.05$) in bold. Model $p$-value is $< 2.2 \cdot 10^{-16}$, adjusted $R^2$ is 0.1498. Model formula: `α ~ Number of sequences + Total branch length + Alignment length + Length of coding sequence + GC content (mean) + GC content (stdev) + Total intron length + Number of exons + Maximum expression + Mean expression + Recombination rate`.

| Variable | Estimate | Std. Error | $t$ value | $p$-value |
|---|---|---|---|---|
| **Number of sequences** | 0.177 | 0.010 | 17.430 | $< 2 \cdot 10^{-16}$ |
| **Total branch length** | 0.118 | 0.012 | 9.683 | $< 2 \cdot 10^{-16}$ |
| **Alignment length** | 0.600 | 0.034 | 17.446 | $< 2 \cdot 10^{-16}$ |
| **Length of coding sequence** | 0.104 | 0.034 | 3.099 | 0.001952 |
| **GC content (mean)** | 0.089 | 0.010 | 8.651 | $< 2 \cdot 10^{-16}$ |
| GC content (stdev) | 0.008 | 0.009 | 0.883 | 0.377265 |
| **Total intron length** | -0.046 | 0.014 | -3.364 | 0.000773 |
| **Number of exons** | 0.152 | 0.019 | 8.144 | $4.67 \cdot 10^{-16}$ |
| Maximum expression | 0.036 | 0.021 | 1.683 | 0.092409 |
| Mean expression | -0.015 | 0.021 | -0.706 | 0.480022 |
| **Recombination rate** | 0.070 | 0.011 | 6.327 | $2.70 \cdot 10^{-10}$ |

Table S12: Linear model of relative support of model with the codon rate variation; Drosophila dataset with 30% of the observations with the highest recombination rates excluded. Significant variables ($p$-value $< 0.05$) in bold. Model $p$-value is $< 2.2 \cdot 10^{-16}$, adjusted $R^2$ is 0.6407. Model formula: `Relative model support ~ Number of sequences + Total branch length + Alignment length + Length of coding sequence + GC content (mean) + GC content (stdev) + Total intron length + Number of exons + Maximum expression + Mean expression + Recombination rate`.

| Variable | Estimate | Std. Error | $t$ value | $p$-value |
|---|---|---|---|---|
| **Number of sequences** | -0.273 | 0.025 | -10.714 | $< 2 \cdot 10^{-16}$ |
| **Total branch length** | 0.096 | 0.024 | 3.951 | $7.88 \cdot 10^{-5}$ |
| Alignment length | -0.140 | 0.087 | -1.616 | 0.106 |
| Length of coding sequence | 0.095 | 0.083 | 1.144 | 0.253 |
| **GC content (mean)** | -0.113 | 0.020 | -5.668 | $1.52 \cdot 10^{-8}$ |
| GC content (stdev) | -0.021 | 0.015 | -1.371 | 0.171 |
| **Total intron length** | -0.125 | 0.023 | -5.552 | $2.95 \cdot 10^{-8}$ |
| Number of exons | 0.006 | 0.024 | 0.231 | 0.817 |
| Maximum expression | -0.014 | 0.035 | -0.400 | 0.689 |
| Mean expression | -0.006 | 0.036 | -0.181 | 0.857 |
| **Recombination rate** | -0.064 | 0.016 | -3.976 | $7.08 \cdot 10^{-5}$ |

Table S13: Linear model of $\alpha$ parameter of gamma distribution with the codon rate variation; Drosophila dataset with 30% of the observations with the highest recombination rates excluded. Significant variables ($p$-value $< 0.05$) in bold. Model $p$-value is $< 2.2 \cdot 10^{-16}$, adjusted $R^2$ is 0.1411. Model formula: `α ~ Number of sequences + Total branch length + Alignment length + Length of coding sequence + GC content (mean) + GC content (stdev) + Total intron length + Number of exons + Maximum expression + Mean expression + Recombination rate`.

| Variable | Estimate | Std. Error | $t$ value | $p$-value |
|---|---|---|---|---|
| **Number of sequences** | 0.194 | 0.008 | 22.865 | $< 2 \cdot 10^{-16}$ |
| **Total branch length** | 0.094 | 0.011 | 8.664 | $< 2 \cdot 10^{-16}$ |
| **Alignment length** | 0.569 | 0.030 | 19.010 | $< 2 \cdot 10^{-16}$ |
| **Length of coding sequence** | 0.128 | 0.029 | 4.366 | $1.28 \cdot 10^{-5}$ |
| **GC content (mean)** | 0.081 | 0.009 | 9.456 | $< 2 \cdot 10^{-16}$ |
| GC content (stdev) | 0.010 | 0.007 | 1.380 | 0.1676 |
| **Total intron length** | -0.052 | 0.011 | -4.566 | $5.03 \cdot 10^{-6}$ |
| **Number of exons** | 0.157 | 0.015 | 10.250 | $< 2 \cdot 10^{-16}$ |
| **Maximum expression** | 0.036 | 0.018 | 2.039 | 0.0414 |
| Mean expression | -0.021 | 0.018 | -1.161 | 0.2455 |
| **Recombination rate** | 0.057 | 0.007 | 8.267 | $< 2 \cdot 10^{-16}$ |
| **Proportion of branches with positive selection** | 0.056 | 0.009 | 6.477 | $9.93 \cdot 10^{-11}$ |

Table S14: Linear model of relative support of model with codon rate variation; Drosophila dataset. Explanatory variables include the proportion of branches with positive selection detected using the branch-site model with codon gamma rate variation. Significant variables ($p$-value $< 0.05$) in bold. Model $p$-value is $< 2.2 \cdot 10^{-16}$, adjusted $R^2$ is 0.6505. Model formula: `Relative model support ~ Number of sequences + Total branch length + Alignment length + Length of coding sequence + GC content (mean) + GC content (stdev) + Total intron length + Number of exons + Maximum expression + Mean expression + Recombination rate + Proportion of branches with positive selection`.

| Variable | Estimate | Std. Error | $t$ value | $p$-value |
|---|---|---|---|---|
| **Number of sequences** | -0.122 | 0.034 | -3.569 | 0.000385 |
| **Total branch length** | 0.156 | 0.038 | 4.075 | $5.17 \cdot 10^{-5}$ |
| **Alignment length** | 0.154 | 0.045 | 3.376 | 0.000778 |
| Length of coding sequence | 0.033 | 0.066 | 0.500 | 0.617029 |
| **GC content (mean)** | 0.168 | 0.046 | 3.686 | 0.000246 |
| GC content (stdev) | 0.031 | 0.038 | 0.826 | 0.409009 |
| Total intron length | -0.057 | 0.057 | -1.000 | 0.317704 |
| Number of exons | -0.038 | 0.071 | -0.537 | 0.591548 |
| Maximum expression | -0.076 | 0.052 | -1.468 | 0.142588 |
| Mean expression | 0.013 | 0.050 | 0.253 | 0.800026 |

Table S15: Linear model of proportion of branches for an alignment identified to be under positive selection by the branch-site model without rate variation, but not identified using branch-site model with codon rate variation, vertebrate dataset. Significant variables ($p$-value $< 0.05$) in bold. Model $p$-value= $3.036 \cdot 10^{-16}$, adjusted $R^2$ is 0.1231. Model formula: `Proportion of false positives ~ Number of sequences + Total branch length + Alignment length + Length of coding sequence + GC content (mean) + GC content (stdev) + Total intron length + Number of exons + Maximum expression + Mean expression`.

| Variable | Estimate | Std. Error | $t$ value | $p$-value |
|---|---|---|---|---|
| Number of sequences | 0.007 | 0.012 | 0.532 | 0.59489 |
| **Total branch length** | 0.241 | 0.014 | 17.376 | $< 2 \cdot 10^{-16}$ |
| **Alignment length** | 0.207 | 0.037 | 5.656 | $1.60 \cdot 10^{-8}$ |
| **Length of coding sequence** | 0.083 | 0.037 | 2.213 | 0.02689 |
| **GC content (mean)** | 0.044 | 0.011 | 3.828 | 0.00013 |
| GC content (stdev) | -0.021 | 0.011 | -1.899 | 0.05766 |
| Total intron length | 0.029 | 0.017 | 1.743 | 0.08140 |
| **Number of exons** | -0.095 | 0.020 | -4.801 | $1.61 \cdot 10^{-6}$ |
| **Maximum expression** | 0.063 | 0.030 | 2.064 | 0.03902 |
| Mean expression | -0.048 | 0.030 | -1.630 | 0.10315 |
| **Recombination rate** | 0.048 | 0.011 | 4.494 | $7.08 \cdot 10^{-6}$ |

Table S16: Linear model of proportion of branches for an alignment identified to be under positive selection by the branch-site model without rate variation, but not identified using branch-site model with codon rate variation, Drosophila dataset. Significant variables ($p$-value $< 0.05$) in bold. Model $p$-value is $< 2.2 \cdot 10^{-16}$, adjusted $R^2$ is 0.113. Model formula: `Proportion of false positives ~ Number of sequences + Total branch length + Alignment length + Length of coding sequence + GC content (mean) + GC content (stdev) + Total intron length + Number of exons + Maximum expression + Mean expression + Recombination rate`.

| Variable | Estimate | Std. Error | $t$ value | $p$-value |
|---|---|---|---|---|
| Number of sequences | -0.000 | 0.015 | -0.019 | 0.98481 |
| **Total branch length** | 0.239 | 0.016 | 14.866 | $< 2 \cdot 10^{-16}$ |
| **Alignment length** | 0.197 | 0.044 | 4.509 | $6.64 \cdot 10^{-6}$ |
| **Length of coding sequence** | 0.102 | 0.044 | 2.295 | 0.02176 |
| **GC content (mean)** | 0.042 | 0.014 | 2.982 | 0.00287 |
| GC content (stdev) | -0.011 | 0.013 | -0.789 | 0.42998 |
| Total intron length | 0.021 | 0.019 | 1.078 | 0.28107 |
| **Number of exons** | -0.100 | 0.023 | -4.376 | $1.23 \cdot 10^{-5}$ |
| Maximum expression | 0.046 | 0.036 | 1.281 | 0.20010 |
| Mean expression | -0.036 | 0.035 | -1.032 | 0.30234 |
| **Recombination rate** | 0.075 | 0.017 | 4.288 | $1.83 \cdot 10^{-5}$ |

Table S17: Linear model of proportion of branches for an alignment identified to be under positive selection by the branch-site model without rate variation, but not identified using branch-site model with codon rate variation; Drosophila dataset with 30% of the observations with the highest recombination rates excluded. Significant variables ($p$-value $< 0.05$) in bold. Model $p$-value is $< 2.2 \cdot 10^{-16}$, adjusted $R^2$ is 0.115. Model formula: `Proportion of false positives ~ Number of sequences + Total branch length + Alignment length + Length of coding sequence + GC content (mean) + GC content (stdev) + Total intron length + Number of exons + Maximum expression + Mean expression + Recombination rate`.

| A | ID | Anatomical entity | Annotated | Significant | Expected | $p$-value | $q$-value |
|---|---|---|---|---|---|---|---|
| FBbt:00003023 | adult abdomen (Drosophila) | 4773 | 1871 | 1747.54 | 1.07 | $1.98 \cdot 10^{-9}$ | $5.04 \cdot 10^{-7}$ |
| FBbt:00001684 | embryonic/larval hemocyte (Drosophila) | 4574 | 1789 | 1674.68 | 1.07 | $3.48 \cdot 10^{-8}$ | 0.00000444 |
| UBERON:0000473 | testis | 5903 | 2258 | 2161.27 | 1.04 | 1e-7 | 0.00000852 |
| UBERON:0001054 | Malpighian tubule | 4706 | 1829 | 1723.01 | 1.06 | $2.47 \cdot 10^{-7}$ | 0.0000158 |
| UBERON:0003917 | arthropod fat body | 5358 | 2060 | 1961.73 | 1.05 | $4.7 \cdot 10^{-7}$ | 0.000024 |
| CL:0000023 | oocyte | 4400 | 1713 | 1610.98 | 1.06 | $8.89 \cdot 10^{-7}$ | 0.0000378 |
| FBbt:00001778 | wing disc (Drosophila) | 4625 | 1791 | 1693.36 | 1.06 | 0.000002 | 0.000073 |
| FBbt:00003143 | adult hindgut (Drosophila) | 4898 | 1880 | 1793.31 | 1.05 | 0.0000164 | 0.000522 |
| FBbt:00000016 | thoracic segment (Drosophila) | 5070 | 1974 | 1856.29 | 1.06 | 0.000234 | 0.00662 |
| UBERON:0000972 | antenna | 6057 | 2277 | 2217.66 | 1.03 | 0.000561 | 0.0143 |
| FBbt:00003133 | crop (Drosophila) | 4741 | 1800 | 1735.83 | 1.04 | 0.00119 | 0.0254 |
| FBbt:00001768 | eye disc (Drosophila) | 4730 | 1796 | 1731.8 | 1.04 | 0.0012 | 0.0254 |
| FBbt:00003138 | adult midgut (Drosophila) | 4284 | 1632 | 1568.51 | 1.04 | 0.00157 | 0.0309 |
| UBERON:0000994 | spermathecum | 4318 | 1640 | 1580.95 | 1.04 | 0.00301 | 0.0548 |
| UBERON:0001017 | central nervous system | 5708 | 2139 | 2089.88 | 1.02 | 0.00556 | 0.0944 |
| B | ID | Anatomical entity | Annotated | Significant | Expected | $p$-value | $q$-value |
| FBbt:00001684 | embryonic/larval hemocyte (Drosophila) | 4574 | 140 | 111.65 | 1.25 | 0.0000132 | 0.00337 |
| UBERON:0000473 | testis | 5903 | 167 | 144.08 | 1.16 | 0.0000338 | 0.0043 |

Table S18: Top anatomical expression enrichment entities for genes with positive selection detected (Drosophila). A) Branch-site model without rate variation B) Branch-site model with codon rate variation. Only anatomical entities with $q$-value $< 0.1$ are shown.

| Parameter | Distribution |
|---|---|
| Number of sequences | DiscreteUniform(8, 12) |
| Alignment length (in codons) | DiscreteUniform(100, 400) |
| $\kappa$ (transition/transversion ratio) | Exponential(1) |
| $\alpha$ (gamma distribution) | 0.1 + Exponential(1) |
| codon 3-rate variation | |
| $p_1$ | Uniform(0, 1) |
| $p_2$ | Uniform(0, 1) |
| $R_1$ | 1/(1+Exponential(1/3)) |
| $R_1$ | 1+Exponential(1/3) |
| M8 | |
| Total branch length | Exponential(1) |
| $\omega$ (positive selection) | Gamma(5, 1) |
| $p$ | 0.2 + Exponential(2) |
| $q$ | 0.2 + Exponential(2) |
| $p_0$ | Beta(8, 2) |
| Branch-site | |
| Total branch length | 10 $\times$ Exponential(1) |
| $\omega_0$ | Beta(1, 5) |
| $p_0 + p_1$ | Beta(10, 1.5) |
| $\frac{p_0}{p_0+p_1}$ | Beta(8, 2) |
| $\omega_2$ (positive selection) | Gamma(50, 1) |

Table S19: Statistical distributions used for simulation parameters.

| Parameter | Transformation |
|---|---|
| Total intron length | $\log(x+1)$ |
| Number of exons | $\sqrt{x}$ |
| Maximum expression | $\log x$ |
| Mean expression | $\log x$ |
| Length of coding sequence | $\log x$ |
| Total branch length | $\log x$ |
| Alignment length (codons) | $\log x$ |
| Relative model support | $\sqrt{\max(x, 0)}$ |
| $\alpha$ (gamma distribution) | $\log x$ |
| $\omega_0$ (branch-site model) | $\log x$ |
| Recombination rate | $\sqrt{x}$ |

Table S20: Variable transformations used for the linear model.

#### Figures

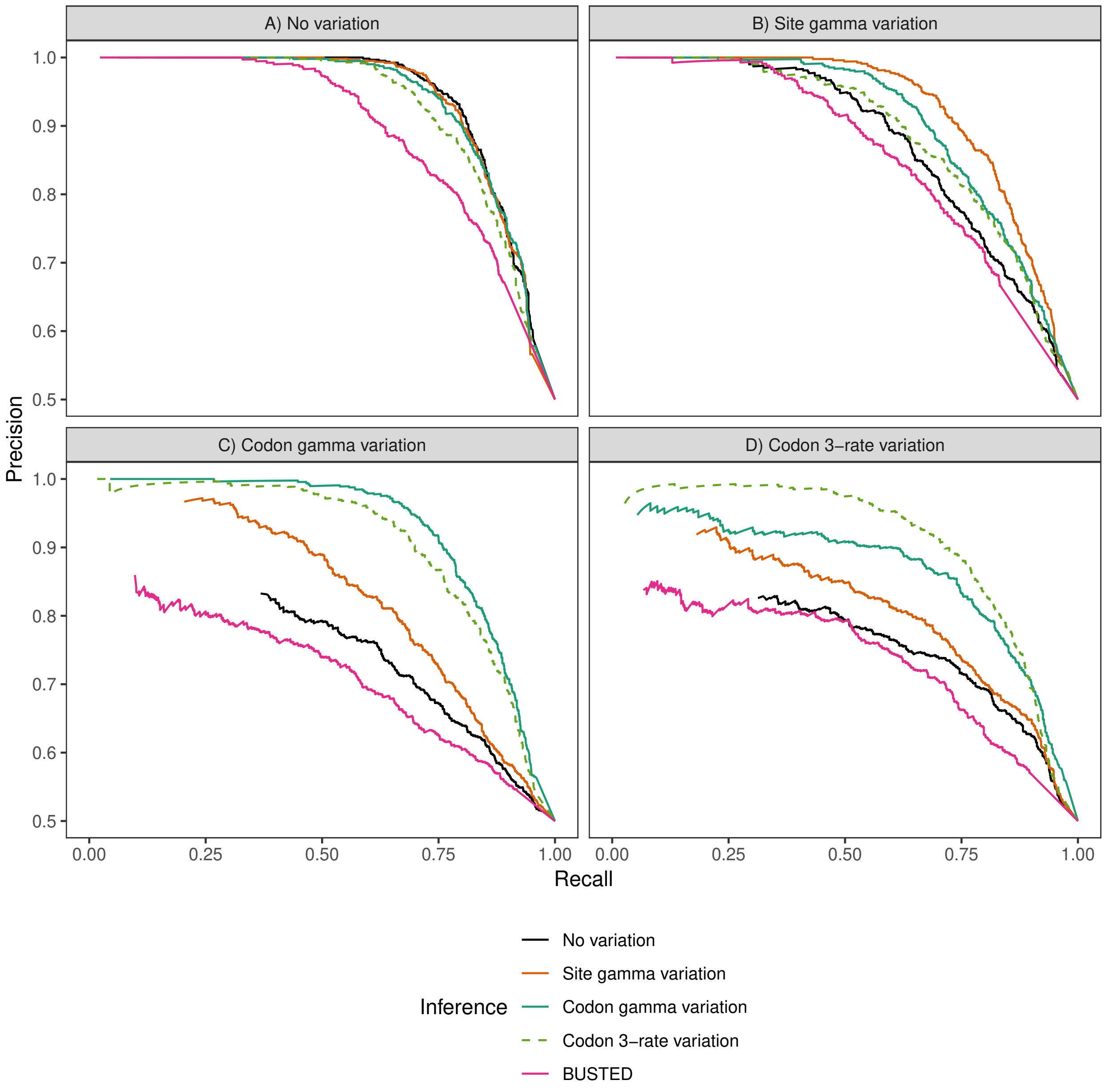

Figure S1: Precision as a function of recall for four M8-based models (M8 with no rate variation, M8 with site rate variation, M8 with codon gamma rate variation, and M8 with codon 3-rate variation) and for BUSTED on datasets A) without rate variation, B) with site rate variation, C) with codon gamma rate variation, and D) with codon 3-rate variation.

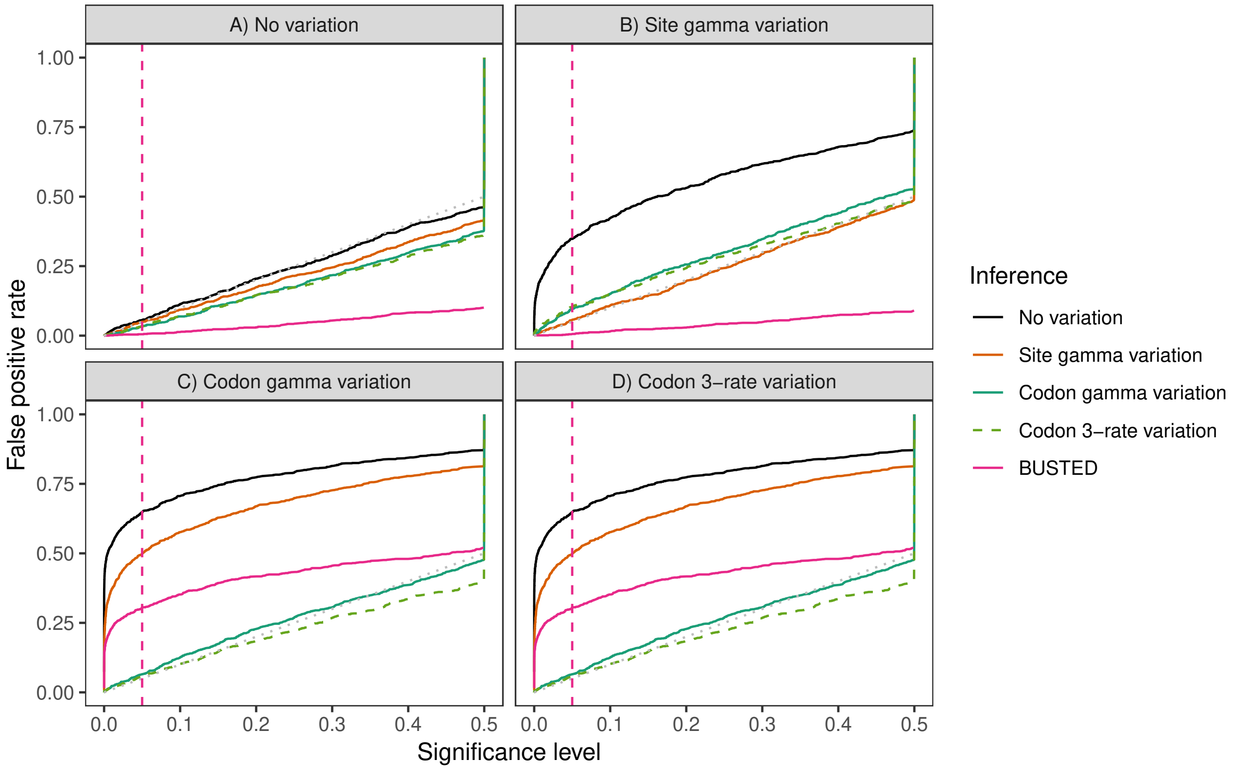

Figure S2: False positive rate as a function of significance level for four M8-based models (M8 with no rate variation, M8 with site rate variation, M8 with codon gamma rate variation, and M8 with codon 3-rate variation) and for BUSTED on datasets A) without rate variation, B) with site rate variation, C) with codon gamma rate variation, and D) with codon 3-rate variation. Vertical dashed line indicates 0.05 threshold. Dashed diagonal line corresponds to the expected false-positive rate under the null hypothesis.

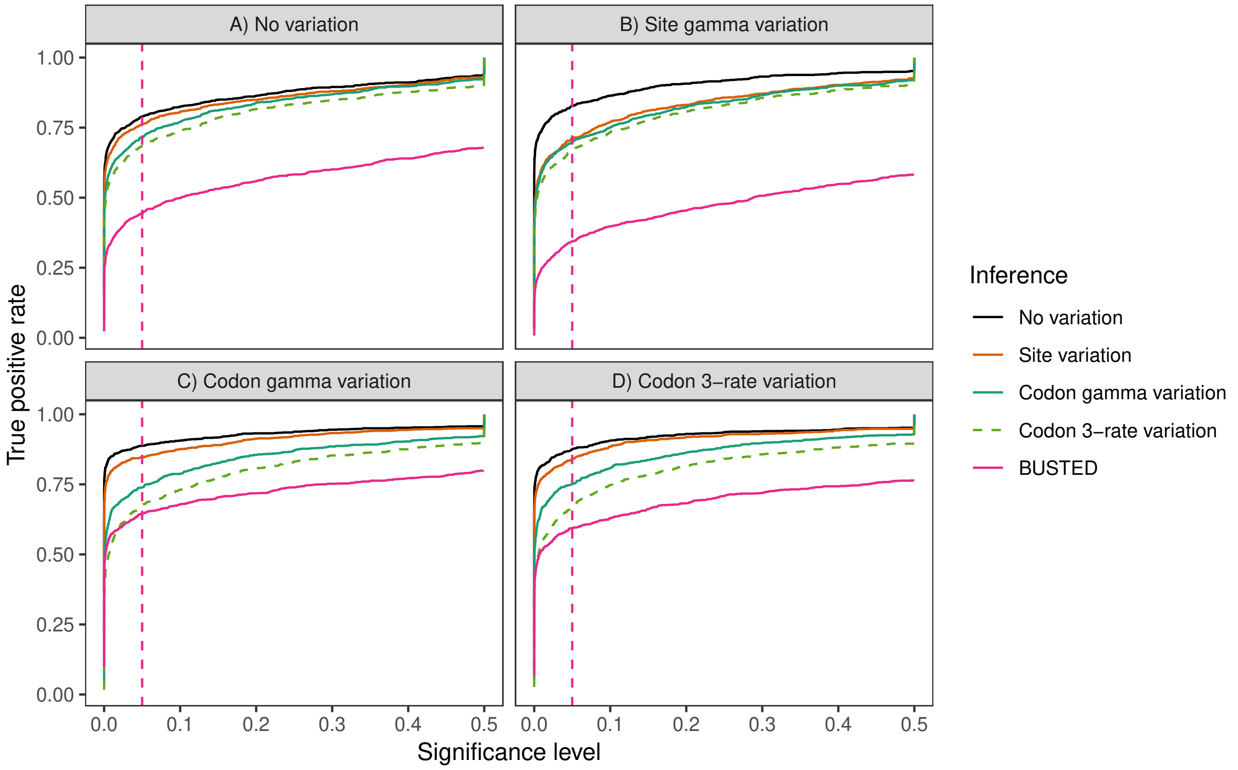

Figure S3: True positive rate (probability of detection) as a function of significance level for four M8-based models (M8 with no rate variation, M8 with site rate variation, M8 with codon gamma rate variation, and M8 with codon 3-rate variation) and for BUSTED on datasets A) without rate variation, B) with site rate variation C) with codon gamma rate variation, and D) with codon 3-rate variation. Vertical dashed line indicates 0.05 threshold.

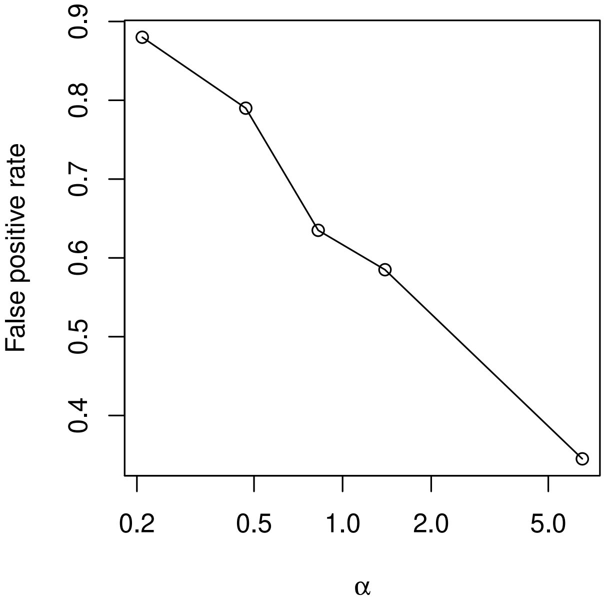

Figure S4: False positive rate as a function of $\alpha$ (gamma distribution parameter, log scale). Smaller $\alpha$ corresponds to stronger rate variation. The data was simulated using the M8 model with codon rate variation and estimated using the M8 model with no rate variation. The data was split into five equal parts according to quantiles of $\alpha$.

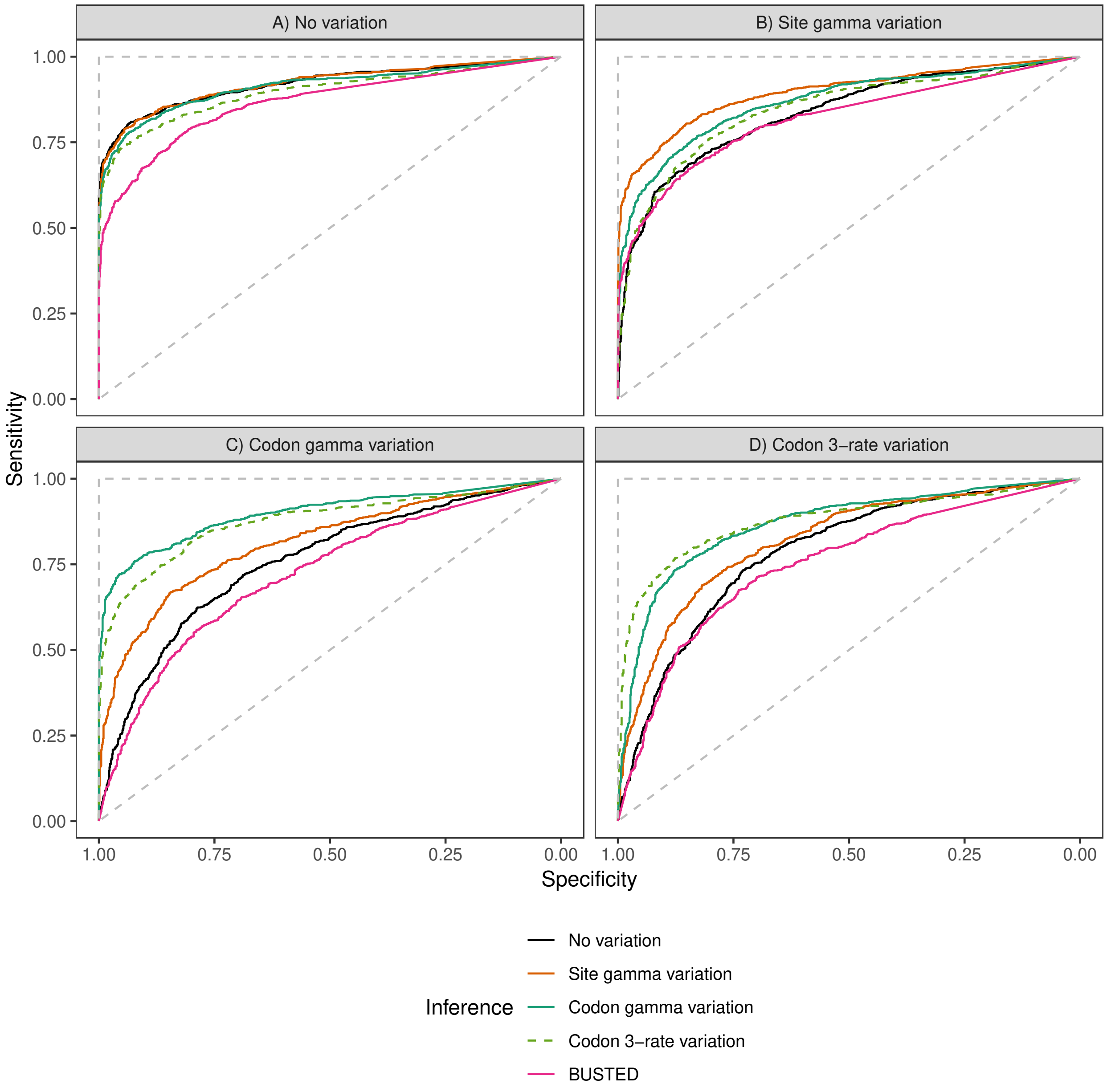

Figure S5: Performance (ROC) of four M8-based models (M8 with no rate variation, M8 with site rate variation, M8 with codon codon rate variation, and M8 with codon 3-rate variation) and of BUSTED on datasets A) without rate variation, B) with site rate variation, C) with codon gamma rate variation, and D) with codon 3-rate variation. True branch lengths were used during estimation. The pink dashed line indicates the 0.95 specificity threshold (i.e., false positive rate of 0.05). The dashed diagonal line shows theoretical performance of the random predictor, the dashed vertical and horizontal lines indicate theoretical performance of the perfect predictor.

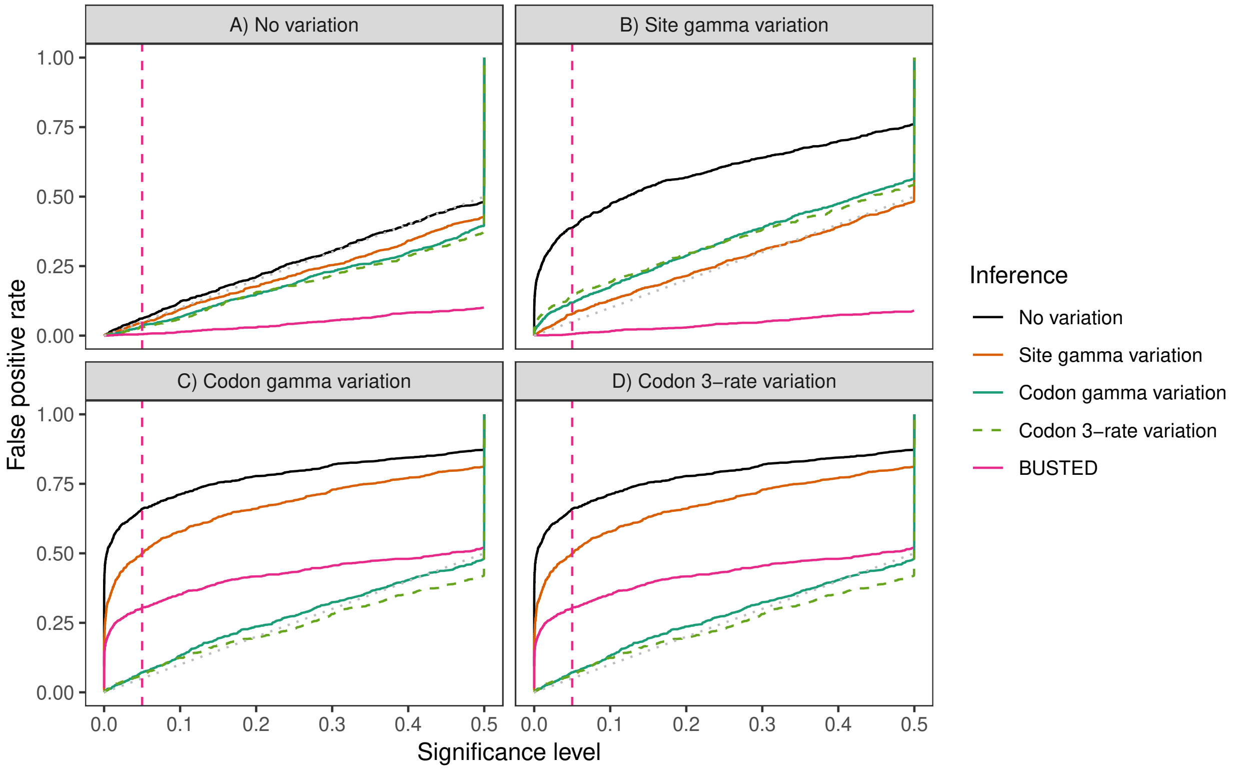

Figure S6: False positive rate as a function of significance level for four M8-based models (M8 with no rate variation, M8 with site rate variation, M8 with codon gamma rate variation, and M8 with codon 3-rate variation) and for BUSTED on datasets A) without rate variation, B) with site rate variation, C) with codon gamma rate variation, and D) with codon 3-rate variation. True branch lengths were used during estimation. Vertical dashed line indicates 0.05 threshold. Dashed diagonal line corresponds to the expected false-positive rate under the null hypothesis.

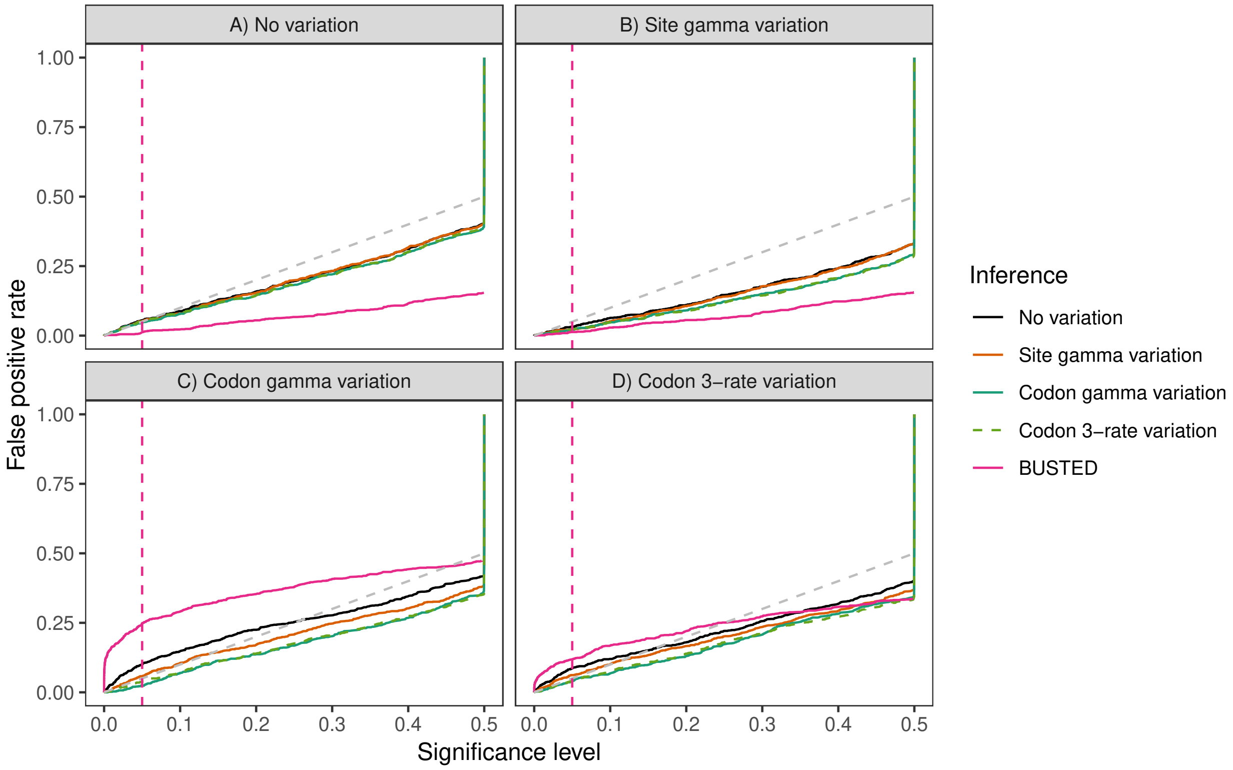

Figure S7: False positive rate as a function of significance level for four branch-site models (branch-site with no rate variation, branch-site with site rate variation, branch-site with codon gamma rate variation, and branch-site with codon 3-rate variation) and for BUSTED on datasets A) without rate variation, B) with site rate variation, C) with codon gamma rate variation, and D) with codon 3-rate variation. Vertical dashed line indicates 0.05 threshold. Dashed diagonal line corresponds to the expected false-positive rate under the null hypothesis.

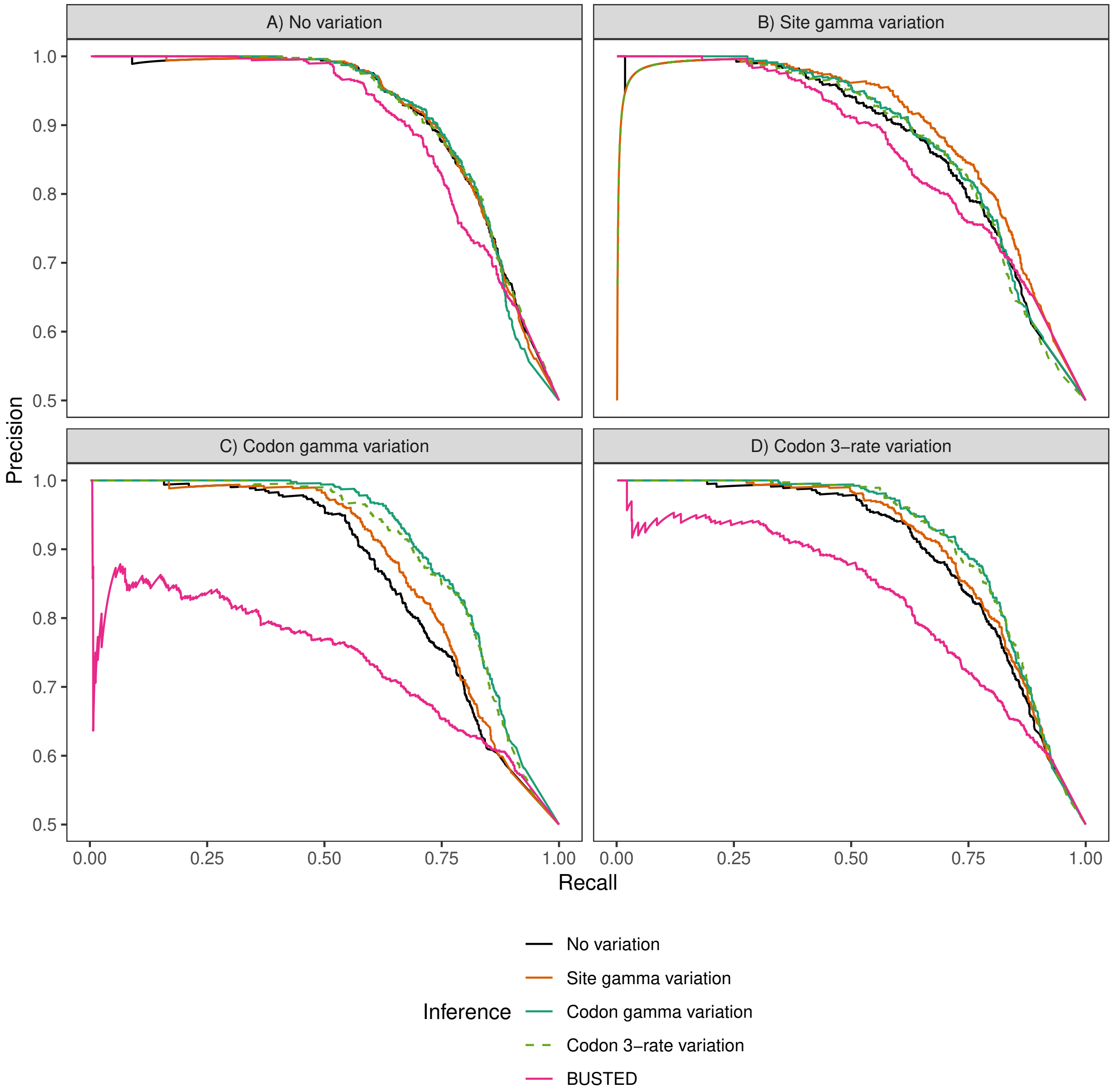

Figure S8: Precision as a function of recall for four branch-site models (branch-site with no rate variation, branch-site with site rate variation, branch-site with codon gamma rate variation, and branch-site with codon 3-rate variation) and for BUSTED on datasets A) without rate variation, B) with site rate variation, C) with codon gamma rate variation, and D) with codon 3-rate variation.

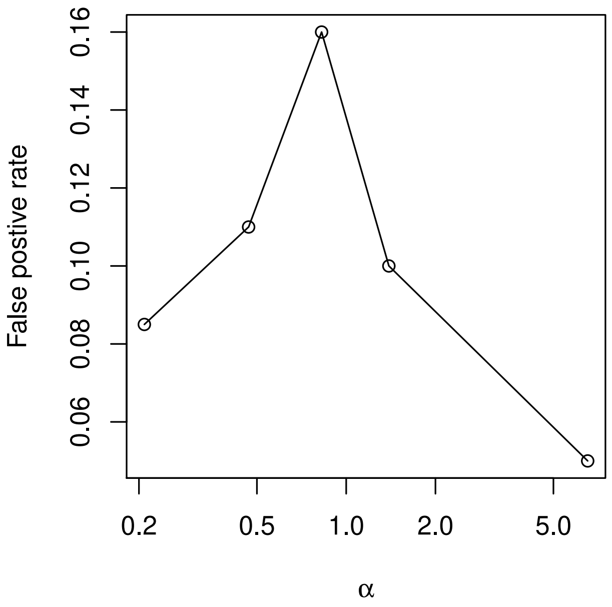

Figure S9: False positive rate as a function of $\alpha$ (gamma distribution parameter, log scale). Smaller $\alpha$ corresponds to stronger rate variation. The data was simulated using the branch-site model with codon rate variation and estimated using the branch-site with no rate variation. The data was split into five equal parts according to quantiles of $\alpha$.

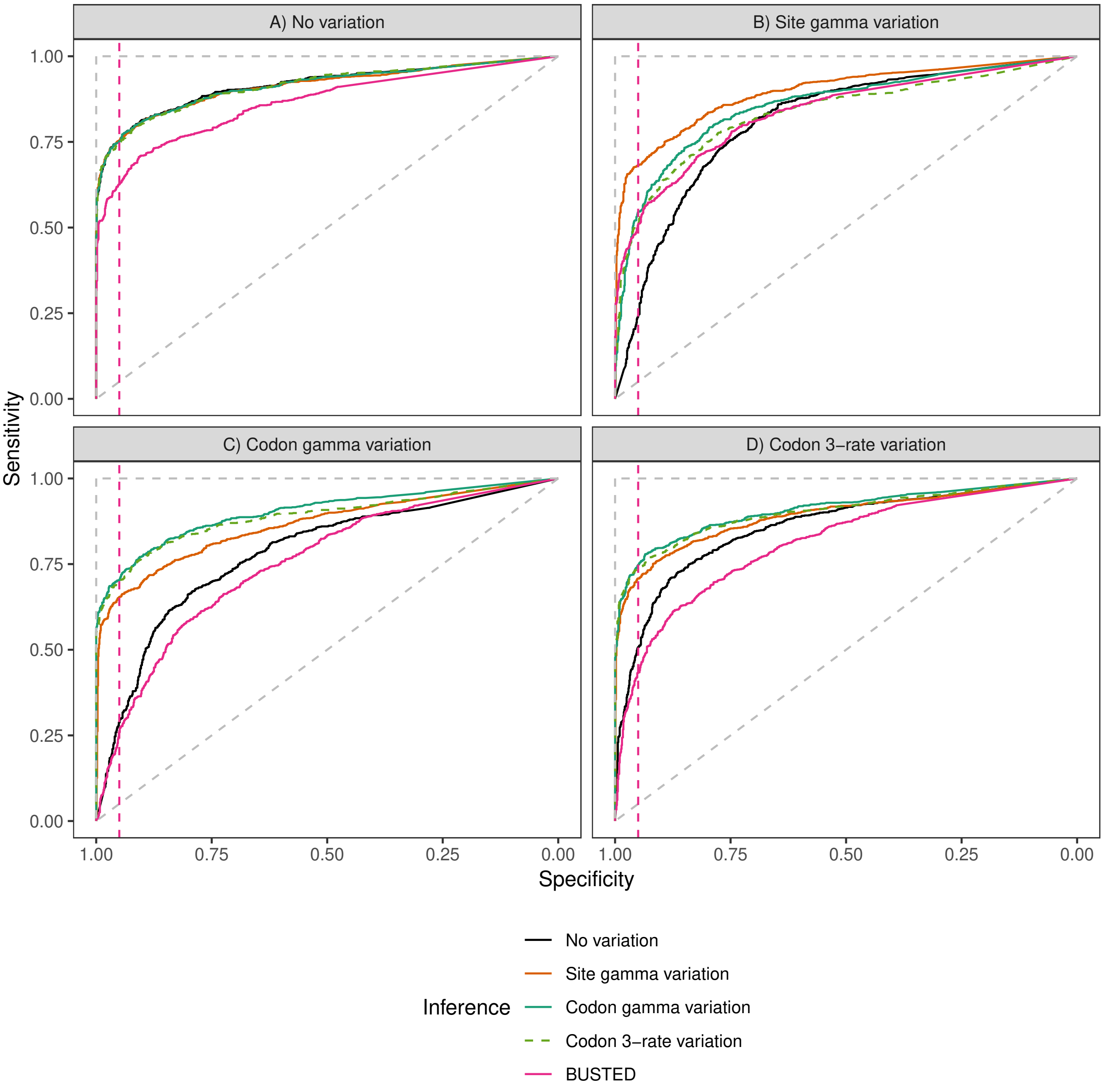

Figure S10: Performance (ROC) of four branch-site-based models (branch-site with no rate variation, branch-site with site rate variation, branch-site with codon 3-rate variation, and branch-site with codon gamma rate variation) and of BUSTED on datasets A) without rate variation, B) with site rate variation, C) with codon gamma rate variation, and D) with codon 3-rate variation. True branch lengths were used during estimation. The pink dashed line indicates the 0.95 specificity threshold (i.e., false positive rate of 0.05). The dashed diagonal line shows theoretical performance of the random predictor, the dashed vertical and horizontal lines indicate theoretical performance of the perfect predictor.

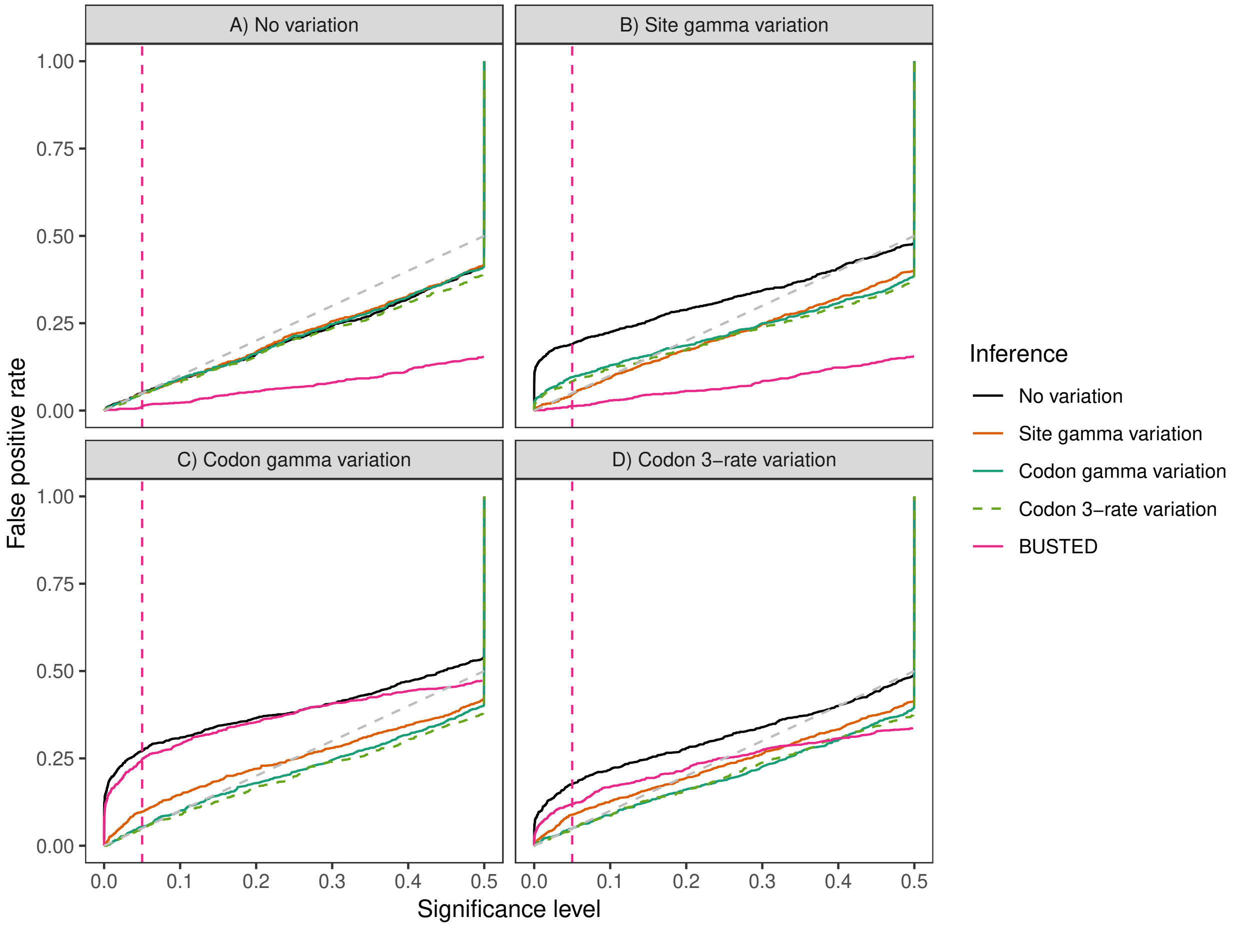

Figure S11: False positive rate as a function of significance level for four branch-site-based models (branch-site model with no rate variation, branch-site model with site rate variation, branch-site model with codon gamma rate variation, and branch-site model with codon 3-rate variation) and for BUSTED on datasets A) without rate variation, B) with site rate variation, C) with codon gamma rate variation, and D) with codon 3-rate variation. True branch lengths were used during estimation. Vertical dashed line indicates 0.05 threshold. Dashed diagonal line corresponds to the expected false-positive rate under the null hypothesis.

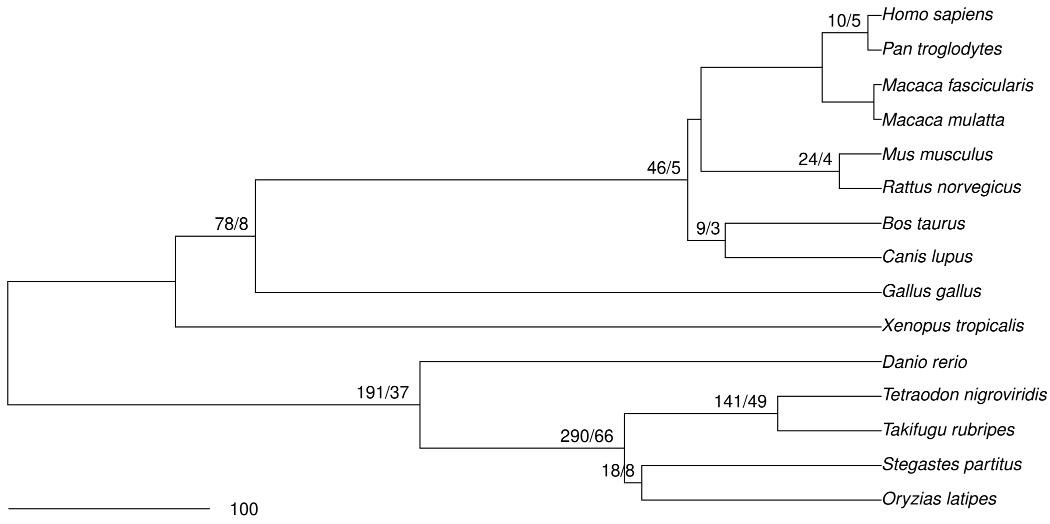

Figure S12: Phylogenetic tree of vertebrate species used in the study. Tree topology and branch lengths in millions of years were acquired from TimeTree (`http://www.timetree.org/`). Labels indicate the number of genes identified as evolving under positive selection at a particular branch (FDR threshold 0.1). The first numbers correspond to the branch-site model without rate variation, the second numbers correspond to the branch-site model with codon gamma rate variation. Only tests performed on branches compatible between the gene tree and the species tree are displayed.

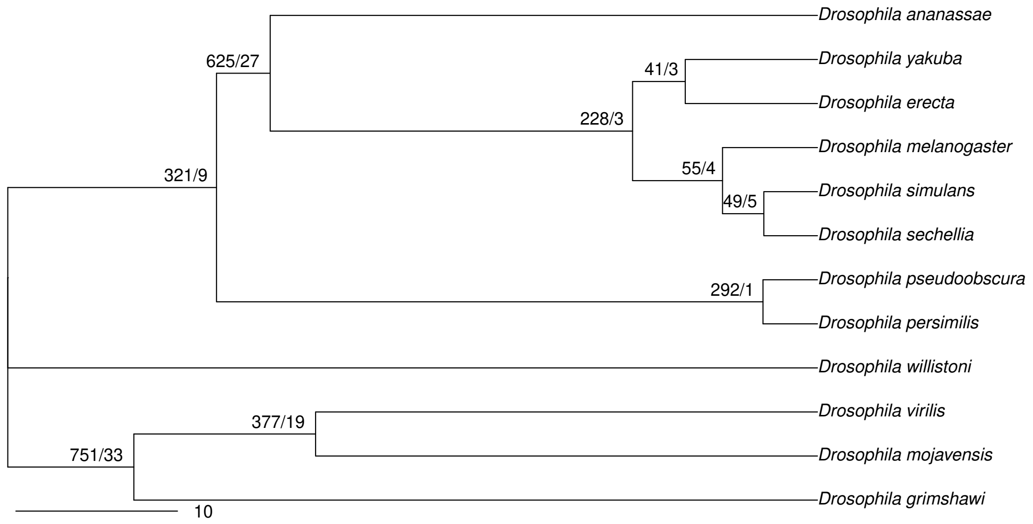

Figure S13: Phylogenetic tree of Drosophila species used in the study. Tree topology and branch lengths in millions of years were acquired from TimeTree (`http://www.timetree.org/`). Labels indicate the number of genes identified as evolving under positive selection at a particular branch (FDR threshold 0.1). The first numbers correspond to the branch-site model without rate variation, the second numbers correspond to the branch-site model with codon gamma rate variation. Only tests performed on branches compatible between the gene tree and the reference tree are displayed.

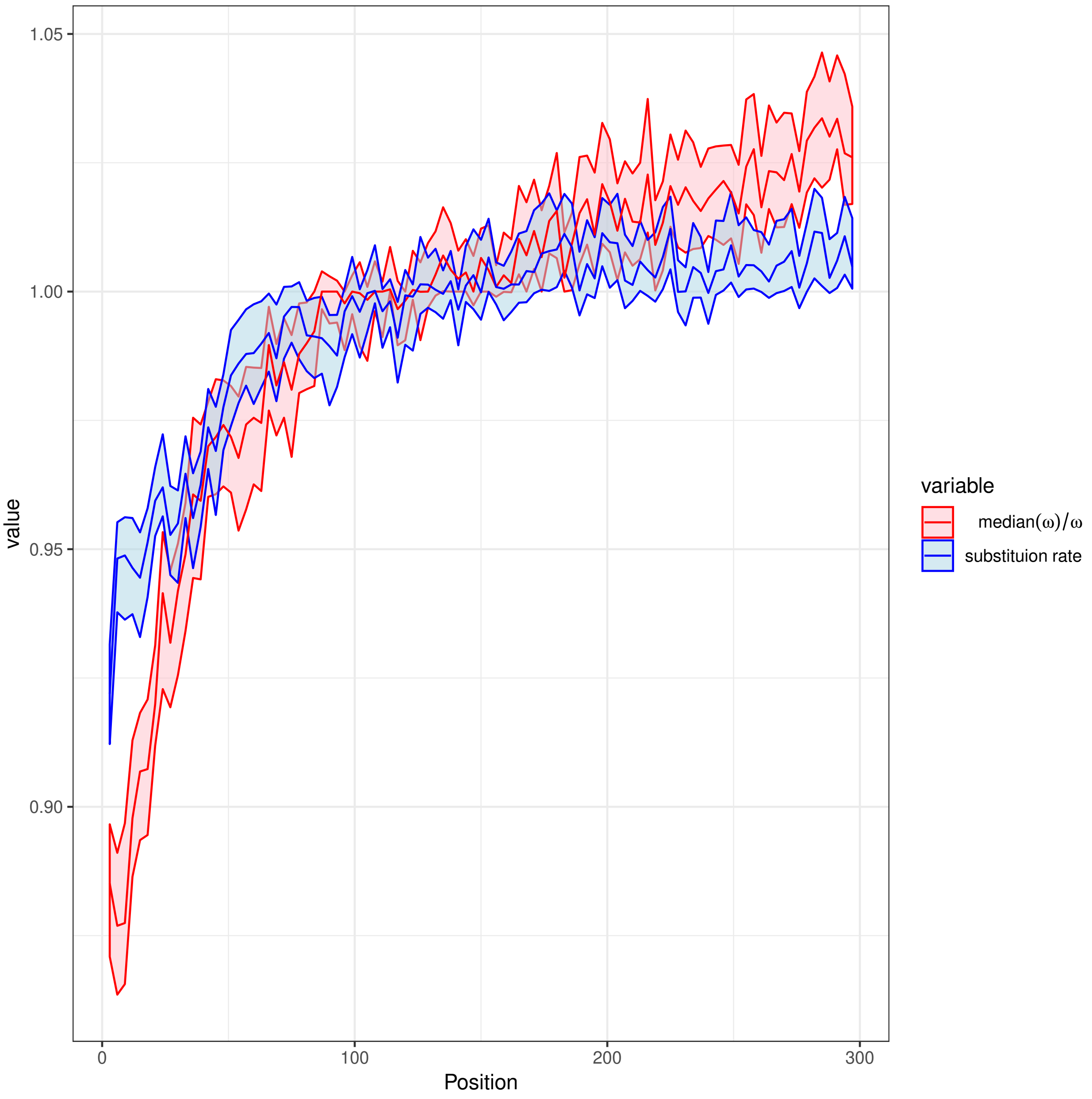

Figure S14: Median posterior estimates of nucleotide substitution rate and inverse protein substitution rates. The latter are defined as $\frac{median(\omega)}{\omega}$; median in the nominator is computed per gene alignment. Displayed as a function of distance from the start codon expressed in the number of nucleotides in Drosophila. The model M8 with codon gamma rate variation was used to estimate both parameters simultaneously. A value of 1 corresponds to the average (nucleotide substitutions) or median (protein substitutions) rate. The ribbons indicate 98% confidence intervals of median estimates. Start codons and alignment positions with less than three sequences were excluded from the plot.

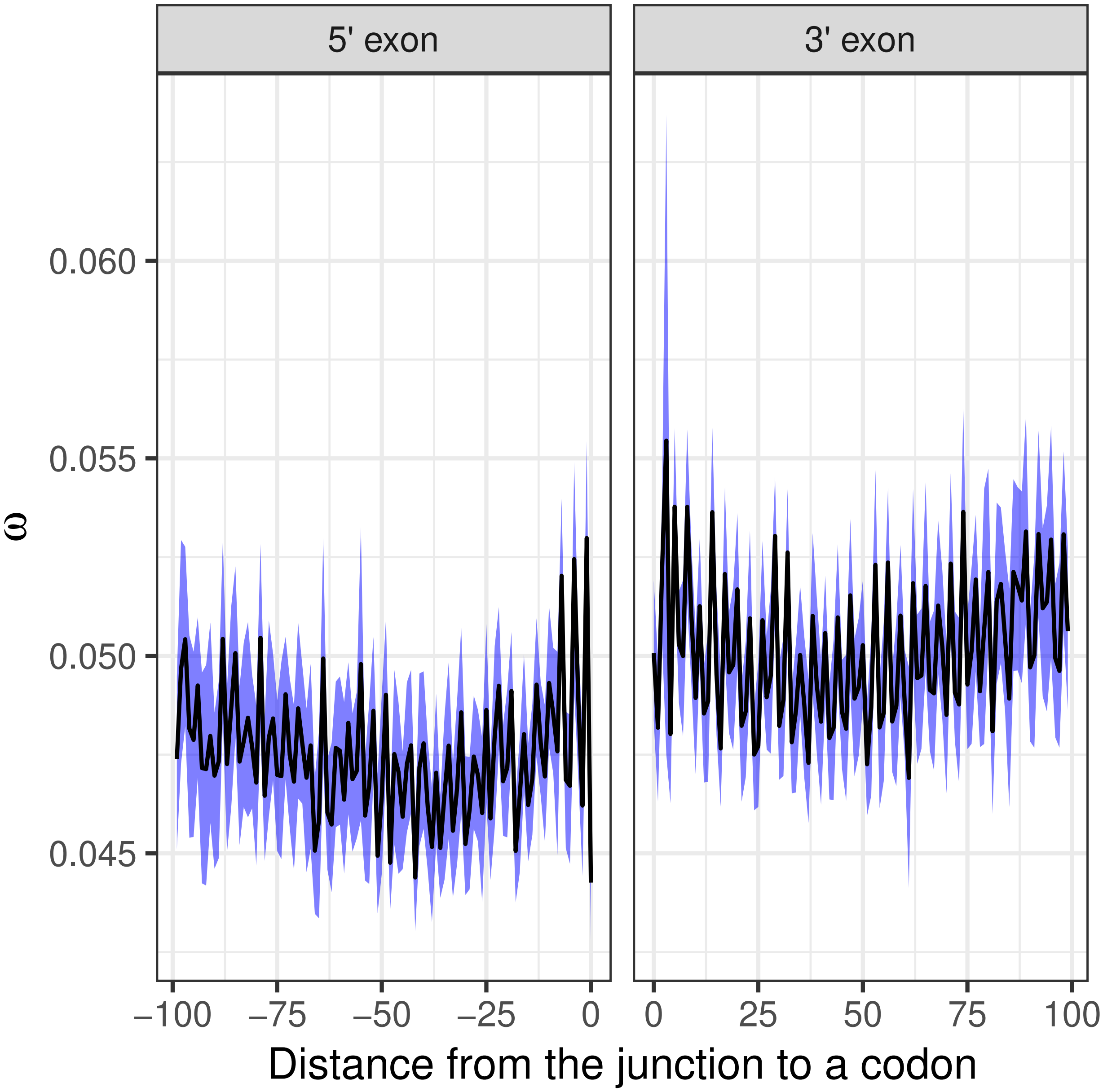

Figure S15: Selection parameter $\omega$ as a function of proximity to the exon-intron and intron-exon junction in the Drosophila dataset. The left panel depicts rates in 5' exon (prior to the exon-intron junction, negative distances), while the right panel depicts 3' exon (rates after the intron-exon junction, positive distances). The blue ribbon indicates 98% confidence interval of mean estimate. Only alignment positions with less than 30% of gaps were used in the plot.

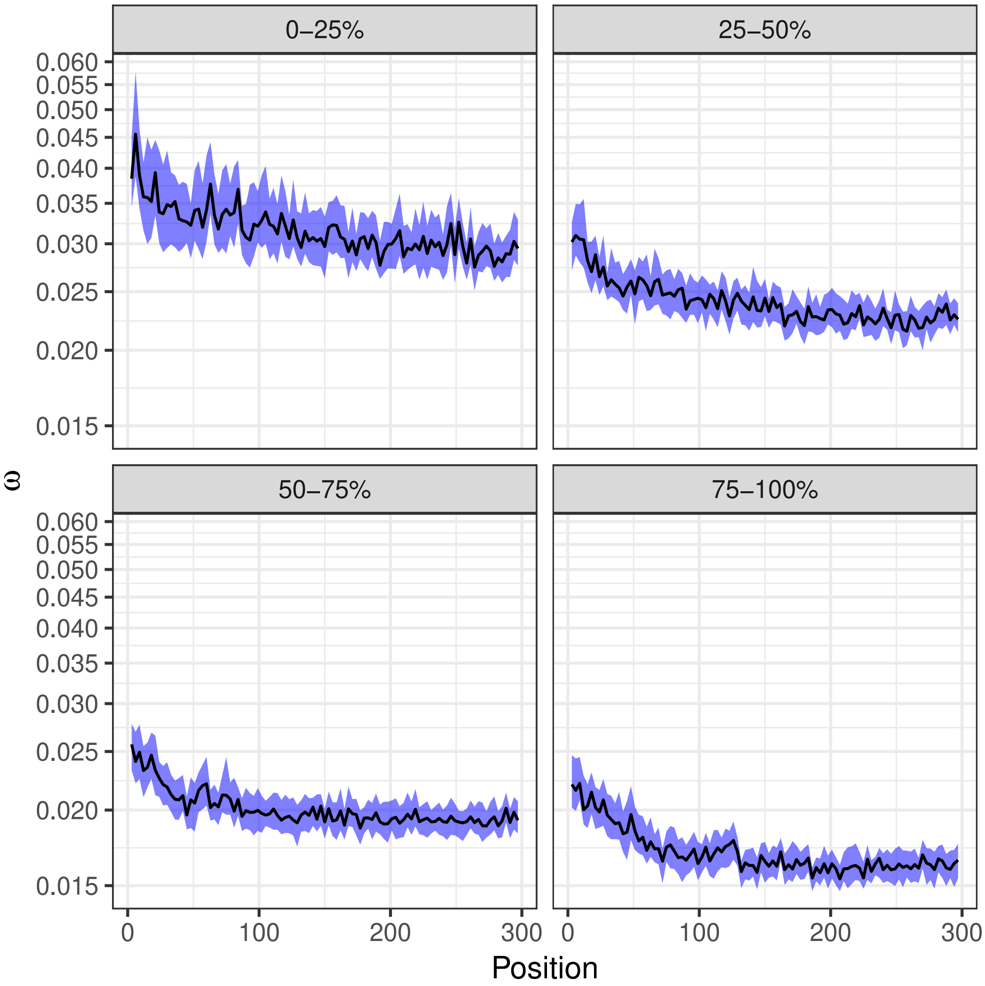

Figure S16: Posterior estimates of median $\omega$ as a function of distance from the start codon expressed in the number of nucleotides. The plot is split into four panels according to mean gene expression quantiles ($[0, 0.25)$, $[0.25, 0.5)$, $[0.5, 0.75)$ and $[0.75, 1]$, percentiles indicated on top of the panels). Logarithmic scale is used for the y axis. The M8 with codon rate variation model was used to estimate codon position specific $\omega$. Smaller values of $\omega$ indicate stronger negative selection acting on the protein sequence. The blue ribbons indicate 98% confidence interval of median estimate. Start codons and alignment positions with less than three sequences were excluded from the plot.

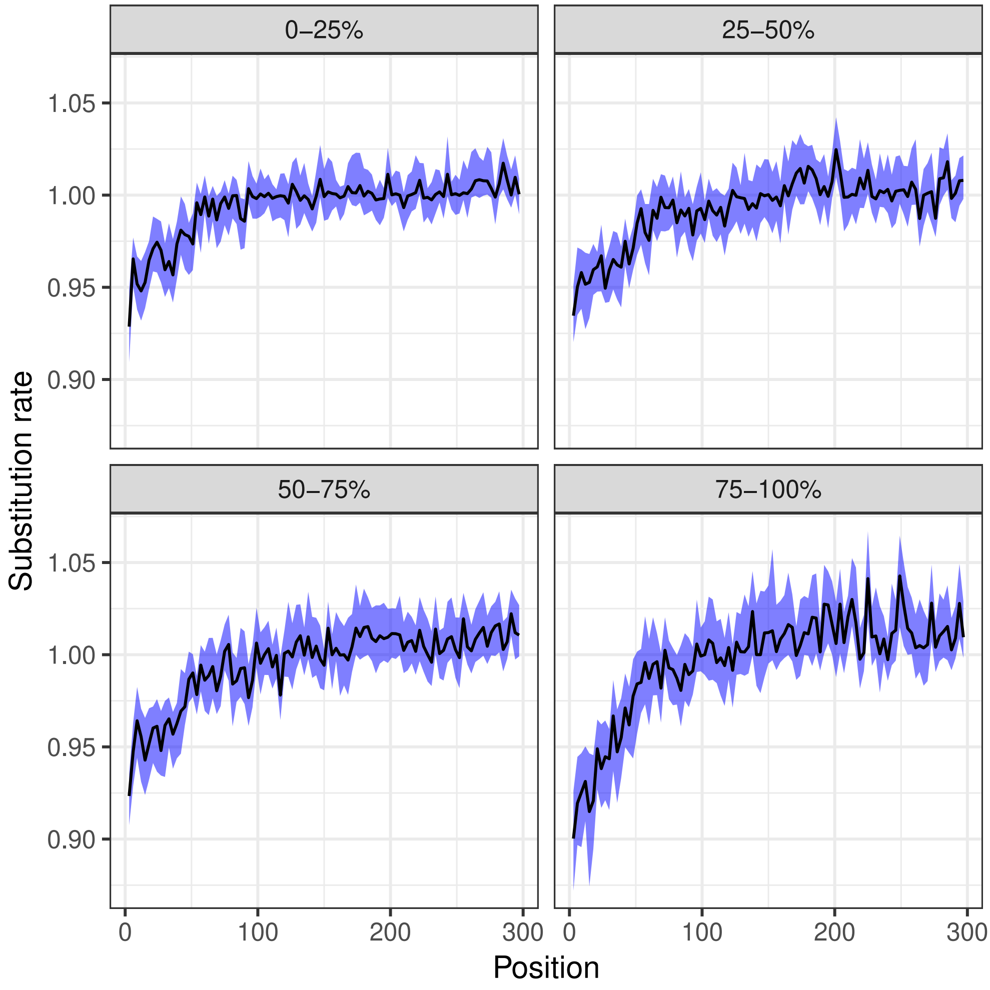

Figure S17: Posterior estimates of median codon substitution rate as a function of distance from the start codon expressed in the number of nucleotides. The plot is split into four panels according to mean gene expression quantiles ($[0, 0.25)$, $[0.25, 0.5)$, $[0.5, 0.75)$ and $[0.75, 1]$, percentiles indicated on top of the panels). The M8 with codon rate variation model was used to estimate the codon position rate. A substitution rate of 1 corresponds to the average rate of substitution over the gene; thus values above 1 do not indicate positive selection, but simply a rate higher than average for this gene. The blue ribbons indicate 98% confidence interval of median estimate. Start codons and alignment positions with less than three sequences were excluded from the plot.

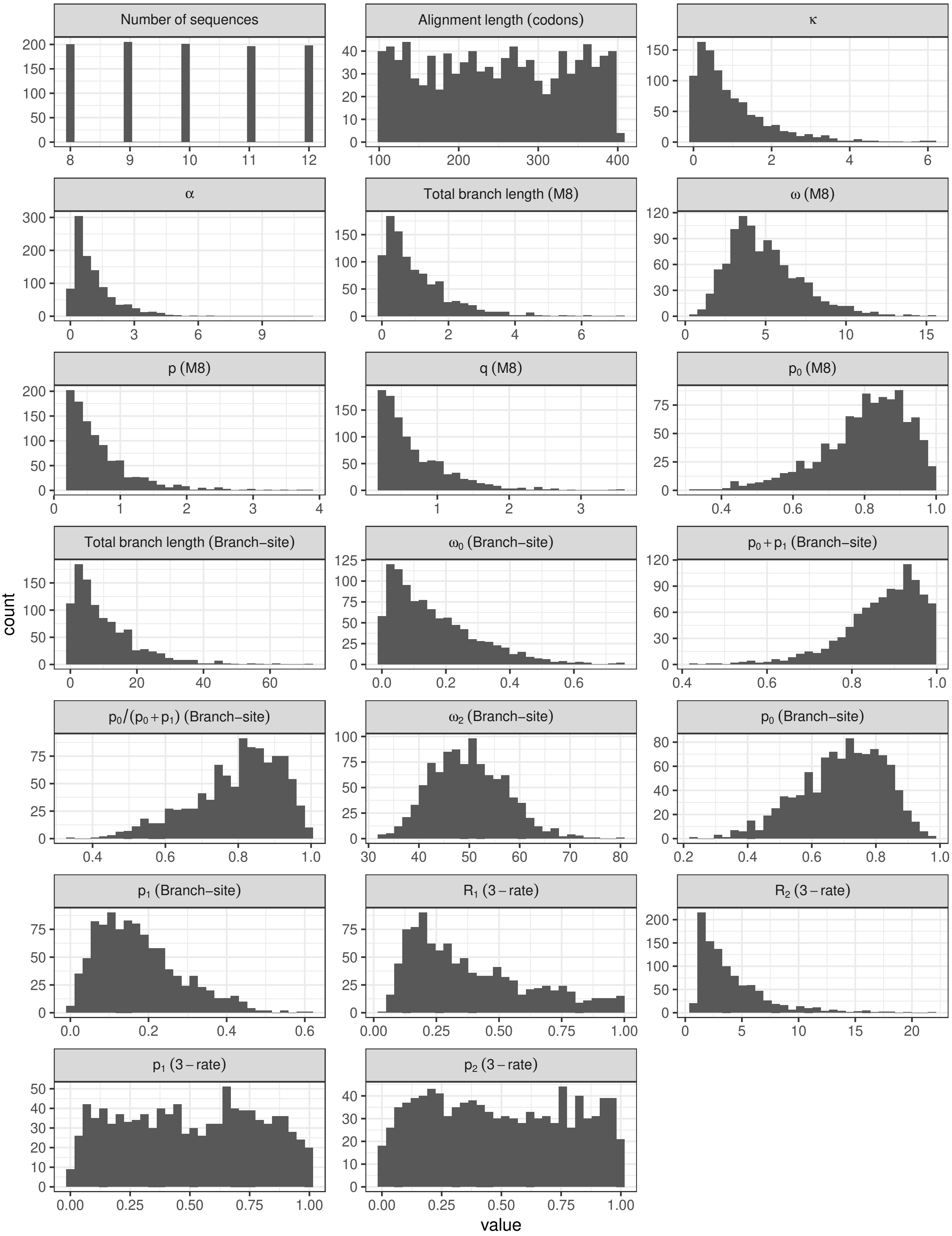

Figure S18: Distribution of simulation parameters.

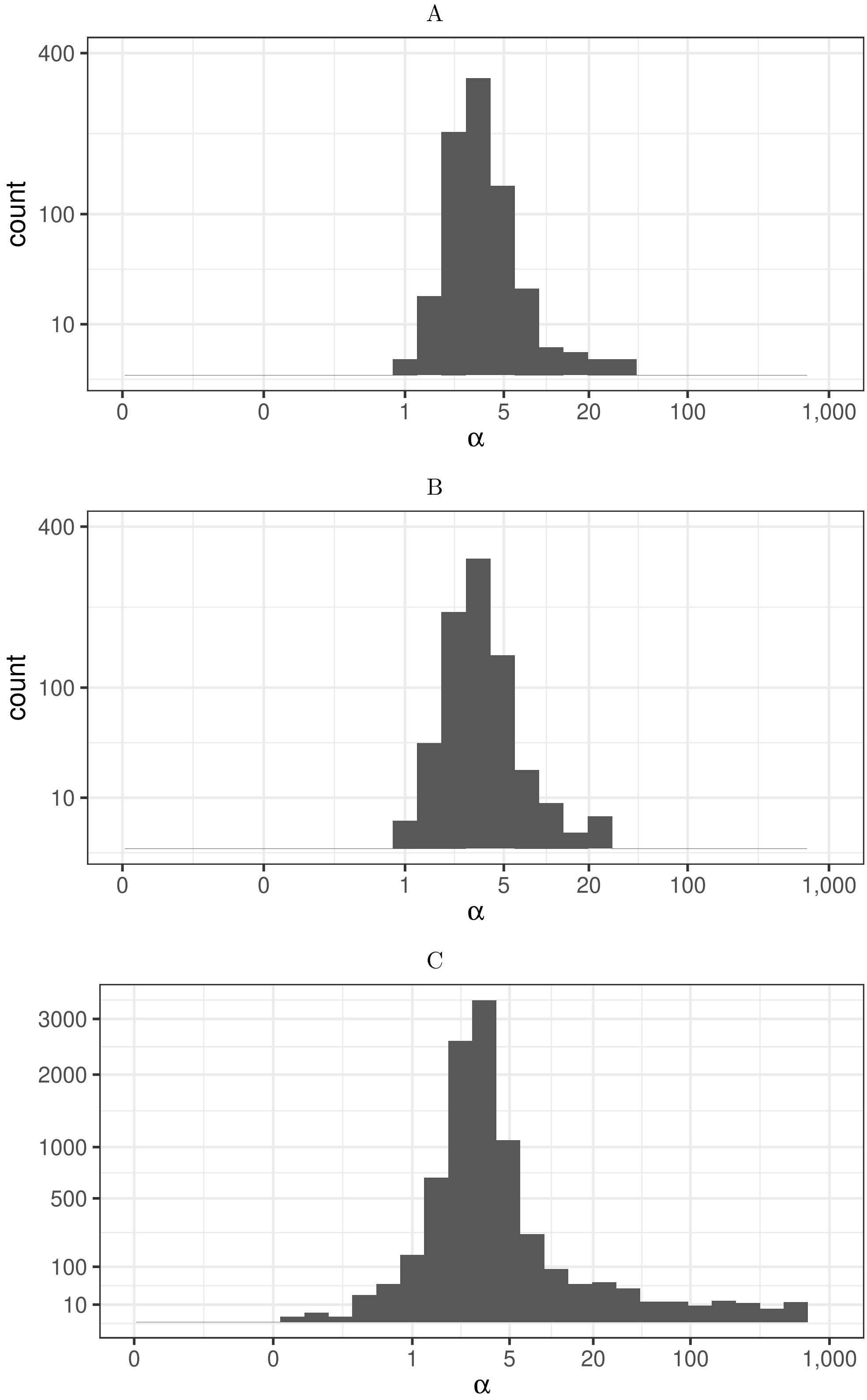

Figure S19: Estimates of $\alpha$ for: A) The vertebrate dataset, codon rate variation; B) the vertebrate dataset, site rate variation; C) the Drosophila dataset. The x axis is log-scaled, the y axis is scaled using the square root.

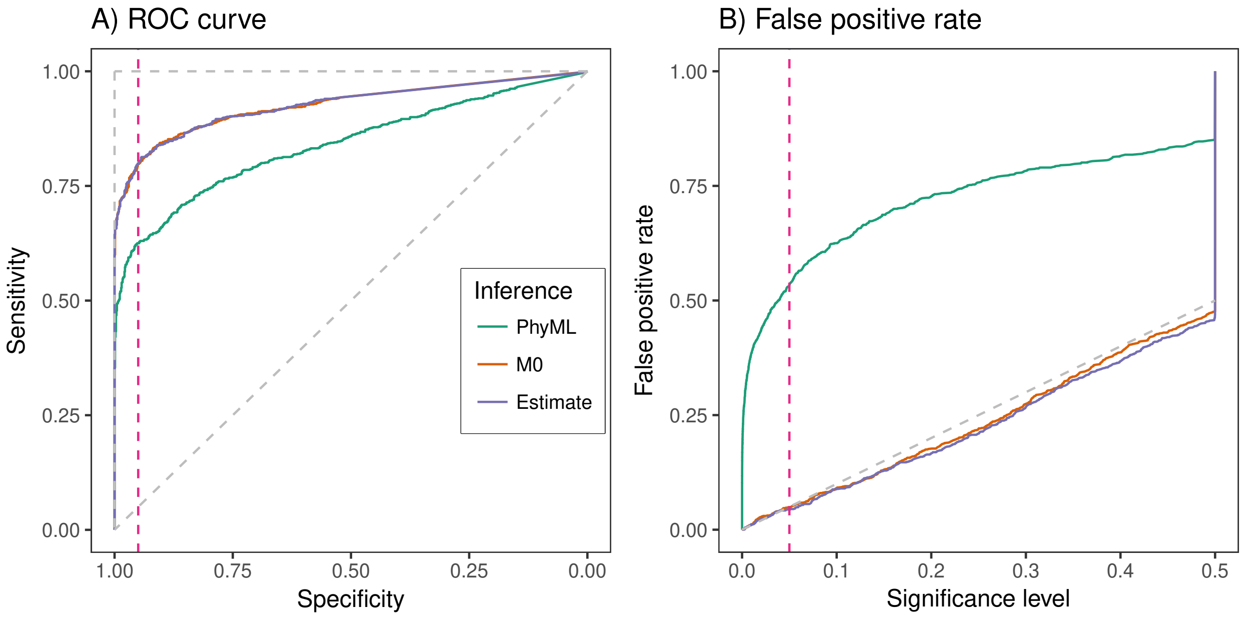

Figure S20: Effect of branch length estimation on positive selection prediction for the M8 model (no rate variation), data simulated under the same model. Three approaches were used to branch length estimation: keep PhyML estimates, estimate using M0, estimate branch lengths and model parameters simultaneously (see legend). A) ROC curve. B) False positive rate as function of significance level. The diagonal dashed line corresponds to the A) performance of a random predictor B) expected false positive rate under the null hypothesis. The vertical pink line corresponds to A) specificity of 95% (false positive rate of 5%), B) significance level of 5%.

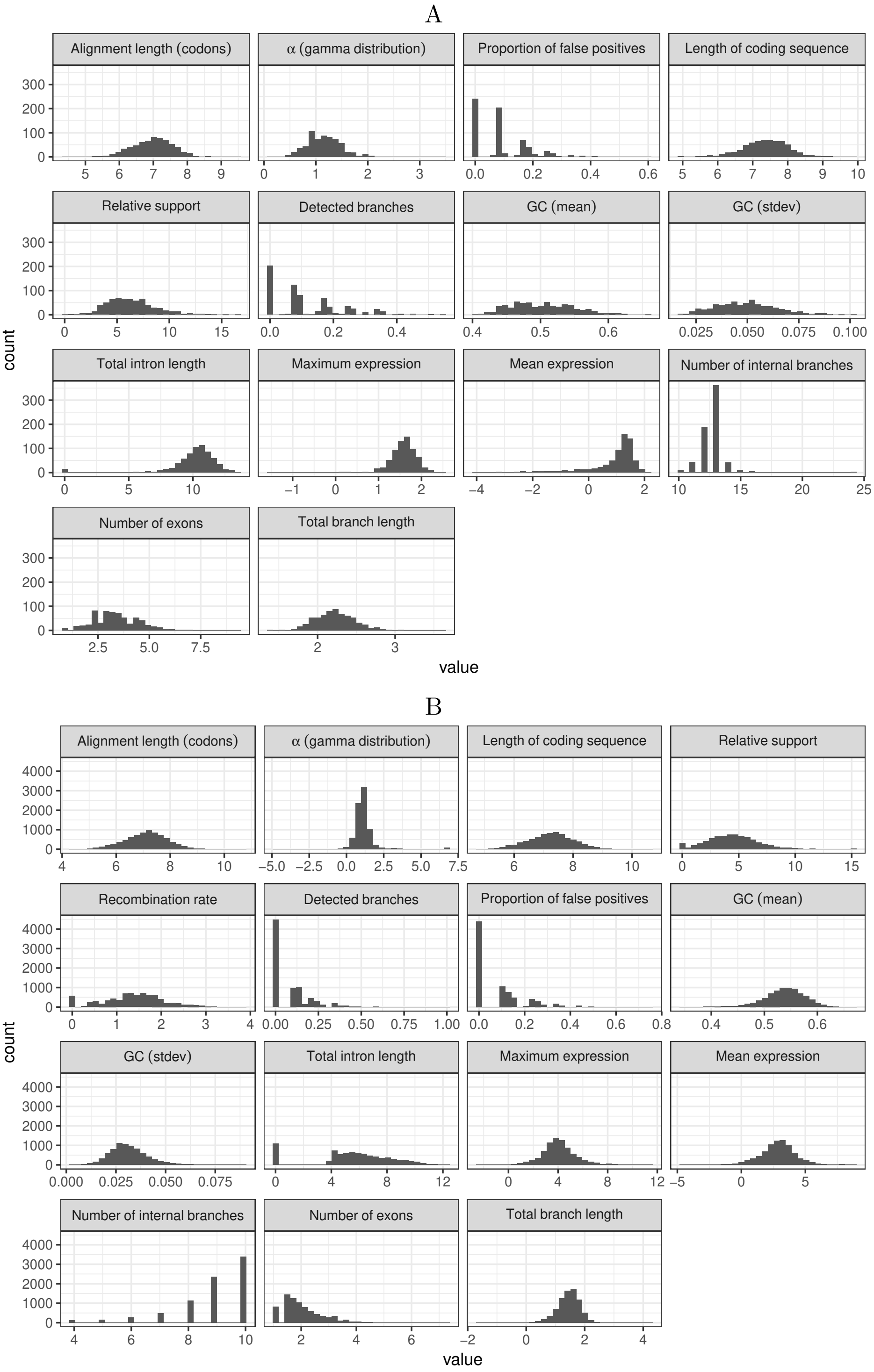

Figure S21: Distribution of transformed variables for the linear model. A) Vertebrate dataset, B) Drosophila dataset. Detected branches is proportion of branches detected using the branch-site model with codon gamma rate variation; proportion of false positives is proportions of branches which are detected by the model without rate variation but not detected by the model with codon gamma rate variation.

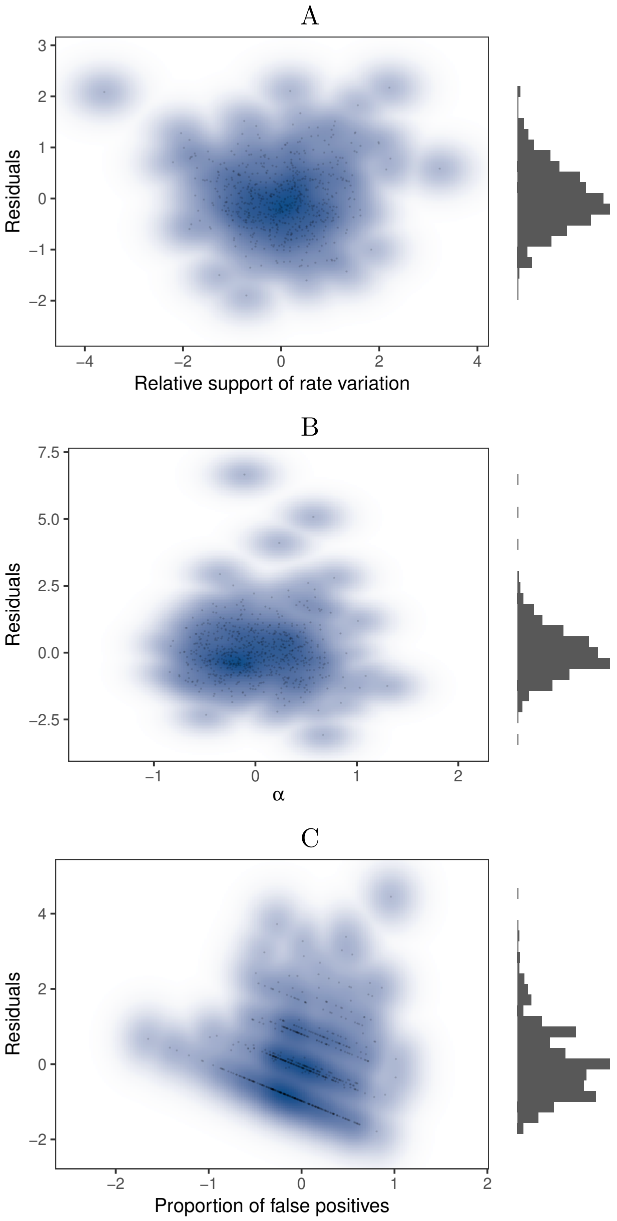

Figure S22: Linear model residuals as a function of the response variable for the vertebrate dataset: A) Relative codon rate variation model support, B) $\alpha$ parameter of gamma distribution, C) Proportion of branches detected by the model without rate variation but not detected by the model with codon gamma rate variation. Response variables were transformed (see methods). Residual histograms are depicted on the right. Points density is indicated with the blue background.

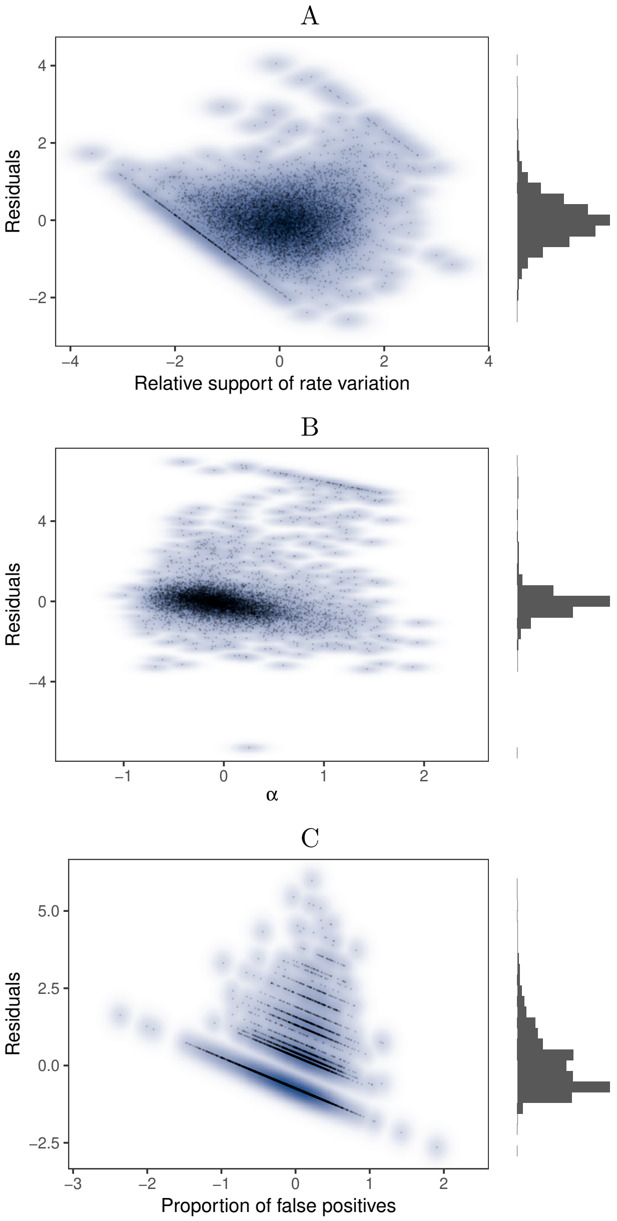

Figure S23: Linear model residuals as a function of the response variable for the Drosophila dataset: A) Relative codon rate variation model support, B) $\alpha$ parameter of gamma distribution, C) Proportion of branches detected by the model without rate variation but not detected by the model with codon gamma rate variation. Response variables were transformed (see methods). Residual histograms are depicted on the right. Density is indicated with the blue background.
